# Genome sequences of four *Ixodes* species expands understanding of tick evolution

**DOI:** 10.1101/2024.02.29.581698

**Authors:** Alexandra Cerqueira de Araujo, Benjamin Noël, Anthony Bretaudeau, Karine Labadie, Matéo Boudet, Nachida Tadrent, Benjamin Istace, Salima Kritli, Corinne Cruaud, Robert Olaso, Jean-François Deleuze, Maarten Voordouw, Caroline Hervet, Olivier Plantard, Aya Zamoto-Niikura, Thomas Chertemps, Martine Maïbèche, Frédérique Hilliou, Gaëlle Le Goff, Jindrich Chmelar, Vilém Mazák, Mohammed Amine Jmel, Michalis Kotsyfakis, José María Medina, Michael Hackenberg, Ladislav Šimo, Fotini A. Koutroumpa, Patrick Wincker, Petr Kopacek, Jan Perner, Jean-Marc Aury, Claude Rispe

## Abstract

Ticks, hematophagous acari, pose a significant threat by transmitting various pathogens to their vertebrate hosts during feeding. Despite advances in tick genomics, high-quality genomes were lacking until recently, particularly in the genus *Ixodes*, which includes the main vectors of Lyme disease. Here, we present the complete genome sequences of four tick species, derived from a single female individual, with a particular focus on the European species *Ixodes ricinus*, achieving a chromosome-level assembly. Additionally, draft assemblies were generated for the three other *Ixodes* species, *I. persulcatus, I. pacificus* and *I. hexagonus*. The quality of the four genomes and extensive annotation of several important gene families have allowed us to study the evolution of gene repertoires at the level of the genus *Ixodes* and of the tick group. We have determined gene families that have undergone major amplifications during the evolution of ticks, while an expression atlas obtained for *I. ricinus* reveals striking patterns of specialization both between and within gene families. Notably, several gene family amplifications are associated with a proliferation of single-exon genes. The integration of our data with existing genomes establishes a solid framework for the study of gene evolution, improving our understanding of tick biology. In addition, our work lays the foundations for applied research and innovative control targeting these organisms.

## Introduction

Ticks are one of a few groups of arthropods that have independently evolved a unique lifestyle of blood-feeding on vertebrates. Present in most terrestrial ecosystems, they represent a threat to companion and farm animals and to humans by transmitting diverse pathogens and parasites (Jongejan & Uilenberg 2004). For example, ticks in the genus *Ixodes* transmit the spirochetes that cause Lyme borreliosis, which is the most common tick-borne disease in the Northern Hemisphere.

Ticks evolved more than 250 million years ago (Mans et al. 2016) and belong to the Parasitiformes, which together with the Acariformes form the Acari (ticks and mites), in the subphylum Chelicerata. Although the Parasitiformes and Acariformes are both monophyletic, the monophyletic status of the Acari has been debated and remains difficult to resolve (Dunlop 2010; Sharma et al. 2014; Lozano-Fernandez et al. 2019; Ballesteros et al. 2019; Zhang et al. 2019; Sharma et al. 2021). Ticks themselves clearly form a monophyletic order (Ixodida), which comprises three families, the Ixodidae (hard ticks), Argasidae (soft ticks) and Nuttalliellidae (Guglielmone et al. 2010; Mans et al. 2011, 2012, 2016). The Ixodidae are further subdivided into the Prostriata, which contains the genus Ixodes, and the Metastriata, which includes several tick genera.

Throughout their history, ticks have evolved remarkable traits to ensure the success of their blood-feeding lifestyle. Molecular level interactions between ticks and their hosts are an essential condition for the success of the blood-feeding strategy of ticks. These interactions principally take place at two host-tick interfaces, represented by the feeding site in the host skin (Mans 2019) and by the ingested blood meal in the tick midgut. Molecules of hosts and ticks interact at these interfaces; for example, tick compounds neutralize host immune and haemostatic responses and thus facilitate tick attachment and blood-feeding on the host (Medina, Jmel, et al. 2022). Tick saliva is mainly produced by the tick salivary glands and facilitates tick-host interactions at the feeding site (Šimo et al. 2017) whereas the tick midgut is the main organ responsible for the digestion of host blood components. These two interfaces represent points of interactions between tick genomes and the hemostatic and immune responses of their vertebrate hosts, exerting a strong selective pressure on ticks and driving a diversification of the tick genetic toolbox.

Gene duplication is believed to shape the major innovations in tick biology, and the duplicate genes would facilitate the evolution of the metabolic potential of these organisms (Mans et al. 2017). To understand the evolution of tick gene repertoires and the importance of tick-specific duplications in particular, a comprehensive comparative study of tick genomes is necessary, which is now possible thanks to the growing number of available genome sequences both in ticks and in other Chelicerata. In comparison with insects, tick genomics has developed late, due to the relatively large genome sizes in this group (several times larger than most insects) (Geraci et al. 2007). The first complete tick genome was published for *Ixodes scapularis* in 2016 (Gulia-Nuss et al. 2016), followed by the genomes of six other tick species, including *Ixodes persulcatus* and five species belonging to a monophyletic group of non-*Ixodes* hard tick species, known as the Metastriata (Jia et al. 2020). Two high quality genome sequences of *I. scapularis* have been published recently (De et al. 2023; Nuss et al. 2023) that used the newer generation of long-read high-throughput sequencing.

The purpose of our study was to improve our knowledge of tick genomics, especially in the genus *Ixodes* which includes some of the most important vectors of tick-borne disease in Europe, North America, and Asia. We therefore generated the genome sequences and gene catalogs of four species, *I. ricinus*, *I. pacificus*, *I. persulcatus*, and *I. hexagonu*s. *Ixodes ricinus*, *I. pacificus*, *I. persulcatus*, and *I. scapularis* are closely related to each other and are part of the *Ixodes ricinus* species complex, whereas *I. hexagonus* represents an outgroup (Charrier et al. 2019; Keirans et al. 1999; Xu et al. 2003). The following species have distinct distributions: *I. scapularis* (Eastern USA), *I. pacificus* (West-coast USA), *I. persulcatus* (Eurasian) and *I. ricinus* (Europe). This ensemble could thus represent an example of vicariant species, corresponding to species that have diverged from a common ancestor in different regions, where they have conserved similar ecological characteristics, as found for the tick genus *Hyalomma* (Sands et al. 2017). Our comprehensive study of the tick gene repertoires in a comparative and phylogenetic framework provides insight into the major aspects that shaped the evolution of tick genomes in the genus *Ixodes* and ticks in general, pointing gene families that have evolved and expanded differently from the rest of Chelicerata, or between different branches within the tick group.

## Results

### Genome assembly and annotation

To enhance our understanding of the genomic characteristics of ticks, particularly in the genus *Ixodes*, we sequenced and assembled the genome of four species: *I. ricinus*, *I. pacificus*, *I. persulcatus* and *I. hexagonus*. The genome sequencing for these four species involved the use of linked reads (10X Genomics library sequence with Illumina short-reads), and for *I. ricinus*, we also incorporated Hi-C libraries to achieve chromosome-level assembly.

In the case of *I. ricinus*, we identified fourteen major scaffolds corresponding to 13 autosomes and the X sex chromosome, which totalled 93% of the total assembly (Hi-C map of interaction, Fig. 1 A). This result is consistent with the established haploid chromosome count of 14 for this species, as in *I. scapularis* (Oliver 1977). We therefore assume that the 14 largest scaffolds of the *I. ricinus* genome, accounting for 95.2% of the assembly size and 98.2% of the predicted genes, represent these 14 chromosomes. By comparison, the 14 largest scaffolds of the *I. scapularis* genome (De et al. 2023) represented 87.0% of the assembly size and 88.2% of the predicted genes, indicating a slightly more fragmented assembly. The genome assemblies of the other three species were organized by aligning them with the chromosomal structure of *I. ricinus*. Standard metrics of the four *Ixodes* genome assemblies sequenced in the present study are given in Table 1.

**Figure 1:**
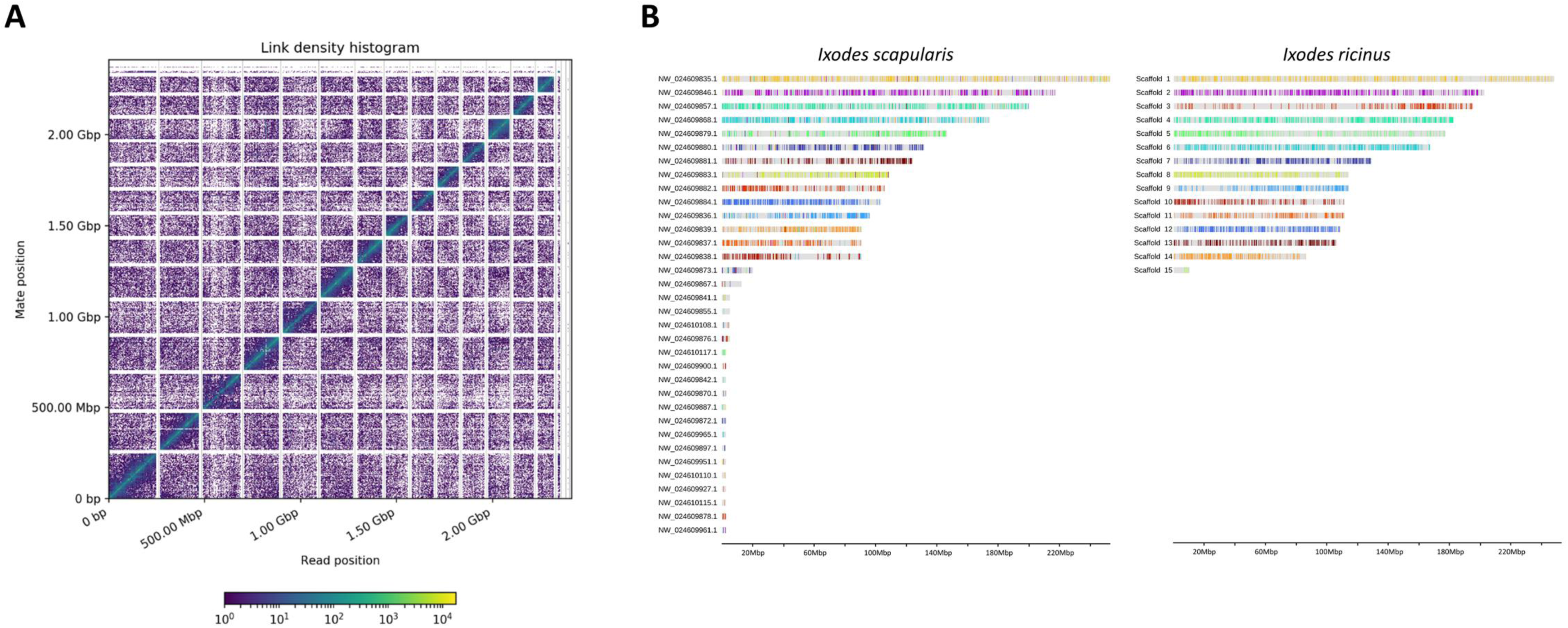
Continuity of the *I. ricinus* genome assembly and synteny with the *I. scapularis* genome. **A** Hi-C map of interactions for the *I. ricinus* genome assembly, showing 14 major scaffolds. The x and y axes give the mapping positions of the first and second read in the read pair respectively, grouped into bins. The color of each square gives the number of read pairs within that bin. Scaffolds less than 1 Mb are excluded. **B** Synteny between the genomes of *I. scapularis* and *I. ricinus*. Horizontal bars represent the major scaffolds of each genome, while syntenies between the two species are indicated by identical colors.

**Table 1:** Metrics of genome assemblies of the four *Ixodes* species, and of their gene catalogs. For *I. ricinus*, the metrics are given prior to HiC scaffolding, and gene counts are prior to manual curation (i.e. for the OGS1.0 version of the prediction).

|  | *Ixodes ricinus* | *Ixodes persulcatus* | *Ixodes pacificus* | *Ixodes hexagonus* |
| --- | --- | --- | --- | --- |
| Estimated genome size | 2.6 Gb | 2.2 Gb | 1.8 Gb | 2.7 Gb |
| Cumulative size | 2,266,064,099 | 2,508,899,395 | 2,295,733,707 | 2,627,922,774 |
| # scaffolds | 76.37 | 178.628 | 199.32 | 143.052 |
| N50 (L50) | 293,124 (1,505) | 79,428 (5,227) | 32,483 (15,538) | 196,001 (2,765) |
| Average scaffold size | 29.672 | 14.045 | 11.517 | 18.37 |
| Max scaffold size | 5,276,202 | 3,084,009 | 903.275 | 4,222,030 |
| Merqury score | 42.2973 | 42.9407 | 42.6649 | 43.9246 |
| Number of predicted genes (OGS 1.0) | 22.486 | 24.979 | 29.201 | 19.511 |
| Average number of exons per gene | 5.65 | 4.67 | 4.06 | 5.44 |
| CDS avg/med length (bp) | 1,124/789 | 1,042/732 | 899/630 | 1,224/918 |
| Genes av/med length (bp) | 30,176/8,214 | 14,586/3,941 | 9,526/3,324 | 21,158/5,354 |
| Introns avg/med length (bp) | 5,301/1,665 | 3,169/1,483 | 2,474/1,313 | 3,964/1,851 |
| ENA accession numbers | PRJEB67792 | PRJEB67789 | PRJEB67791 | PRJEB67790 |

The four genome assemblies were annotated, and genes were predicted by using homologies with proteins of closely related species and RNA-Seq data. Manual curation was performed exclusively for *I. ricinus* resulting in the OGS1.1 gene catalog, which resulted in the correction of 1,569 genes (see supplementary Table S1). The most notable change was the prediction of 500 entirely new gene models (supplementary Table S2). The completeness (% of complete BUSCOs) of the four new gene catalogs generated in this study fell within the range of recently sequenced tick genomes as shown in Table 2. Completeness was lowest in *I. pacificus* (81%), and highest in *I. ricinus* and *I. hexagonus* (about 91%), which is somewhat lower than the 98% observed for the recently improved genome of *I. scapularis* (De et al. 2023). For *I. pacificus*, we also note a relatively high percentage of “duplicated” genes in the BUSCO analysis, suggesting that heterozygosity might have not been fully resolved and that our assembly still contains duplicate alleles, which is supported by the higher heterozygosity estimate for this genome (supplementary Fig. S1). Finally, two tick genomes from another study (Jia et al. 2020), *Hyalomma asiaticum* and *Haemaphysalis longicornis*, showed significantly lower completeness (65% and 56% of complete BUSCOs, respectively), whereas the completeness for *I. persulcatus* in that study was lower compared to our study (79.6% versus 88.0%). For subsequent analyses involving *I. persulcatus*, we utilized our assembly as the reference genome.

**Table 2:**
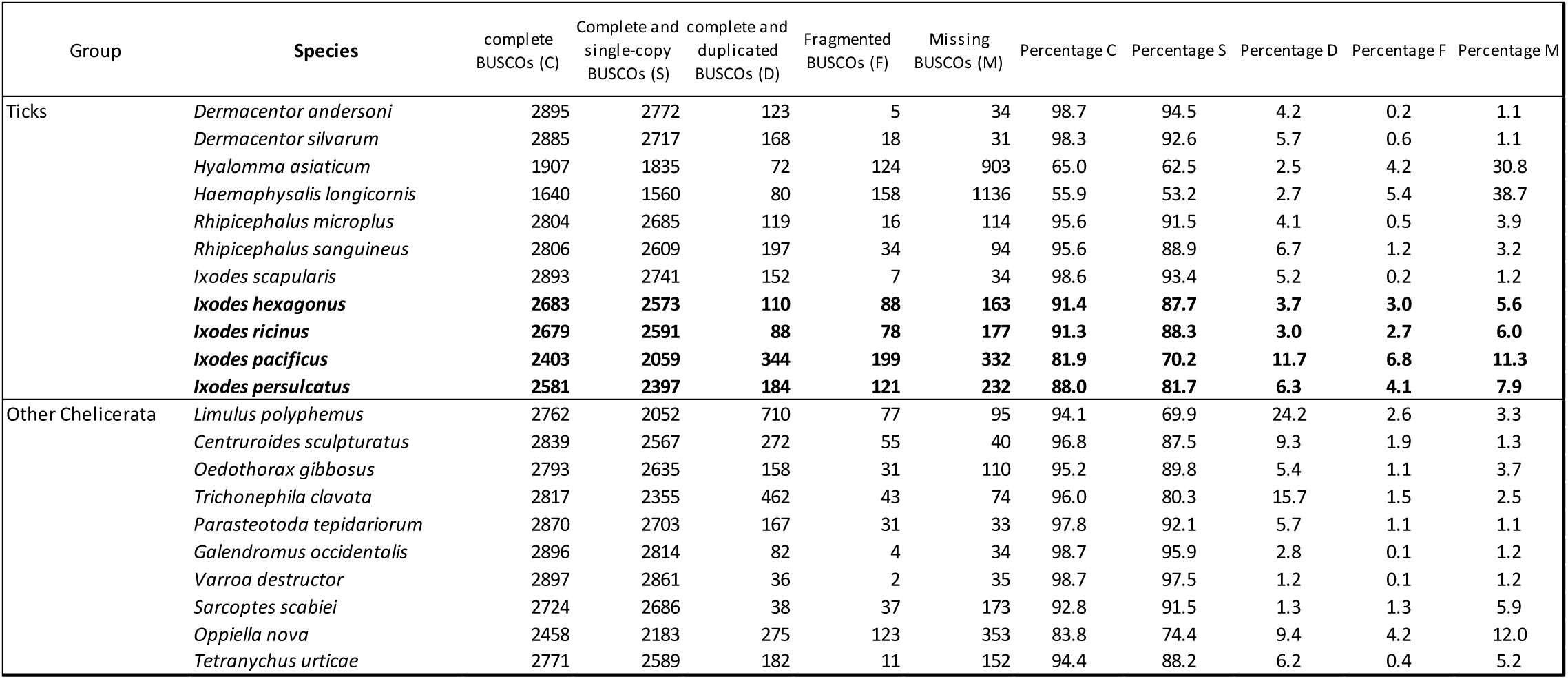
Completeness of the species of Chelicerata included in our comparative study. The four *Ixodes* genomes sequenced in this study are in bold character. The search of conserved genes was made using BUSCO with the Arachnida odb_10 database (search of 2934 conserved genes). Group (ticks or other Chelicerata) and species name, Numbers (five next columns) or Percentages (five last columns) of different categories of BUSCO genes, as detailed in the headers.

### Transposable elements in ticks

In *I. ricinus, I. hexagonus, I. pacificus and I. persulcatus*, repeated elements represented between 57.3% (*I. ricinus*) and 67.9% (*I. hexagonus*) of the genome, the majority being transposable elements (Table 3). Most of the transposable elements (TEs) identified are unclassified (79.83% of the total elements, and covering ∼43% of the tick genomes). The most abundant TEs found in these tick species were long interspersed nuclear elements (LINEs), with 397,287 elements on average (∼7% of the elements identified). Interspersed sequences represented only 9.5% of the genome of *Amblyomma maculatum* (Ribeiro et al. 2023), whereas they accounted for over 50% of the genome of all *Ixodes* ticks. However, the relative frequencies of each TE family were similar between the genomes of *A. maculatum* (Jose M. C. Ribeiro et al. 2023) and *I. scapularis* (De et al. 2023). For example, the Gypsy/DIRS family in the long terminal repeat (LTR) class has one of the highest coverages in both *A. maculatum* (Ribeiro et al 2023) and *I. scapularis*, (Nuss et al 2023, De et al 2023), and represent ∼5% of the genome in the four *Ixodes* species sequenced in the present study.

**Table 3:** Repeated elements in the genomes of the four *Ixodes* species (*I. ricinus*, *I. hexagonus*, *I. pacificus* and *I. persulcatus*). *: Most repeats fragmented by insertions or deletions have been counted as one element.

| Element family | *Ixodes ricinus* |  | *Ixodes hexagonus* |  | *Ixodes pacificus* |  | *Ixodes persulcatus* |  |
| --- | --- | --- | --- | --- | --- | --- | --- | --- |
|  | Number of elements* | Percentage of genome coverage | Number of elements* | Percentage of genome coverage | Number of elements* | Percentage of genome coverage | Number of elements* | Percentage of genome coverage |
| Retroelements (class I): | 467 260 | 11.74 % | 563 828 | 14.96 % | 606 344 | 14.30 % | 677 841 | 14.45 % |
| SINEs: | 692 | 0.01 % | 795 | 0.00 % | 935 | 0.01 % | 773 | 0.00 % |
| LINEs: | 318 541 | 7.14 % | 385 768 | 9.00 % | 429 800 | 9.31 % | 455 040 | 8.53 % |
| Penelope | 68 082 | 1.10 % | 52 969 | 0.76 % | 94 523 | 1.29 % | 74 359 | 1.04 % |
| L2/CR1/Rex | 158 040 | 3.39 % | 138 618 | 2.66 % | 200 079 | 4.04 % | 227 456 | 3.98 % |
| R1/LOA/Jockey | 31 715 | 1.22 % | 97 692 | 3.18 % | 55 214 | 2.22 % | 54 525 | 1.69 % |
| R2/R4/NeSL | 70 | 0.00 % | 0 | 0.00 % | 141 | 0.00 % | 619 | 0.01 % |
| RTE/Bov-B | 17 064 | 0.38 % | 24 517 | 0.72 % | 24 545 | 0.56 % | 22 648 | 0.56 % |
| L1/CIN4 | 9 668 | 0.24 % | 4 572 | 0.13 % | 6 385 | 0.14 % | 12 171 | 0.21 % |
| LTR elements: | 148 027 | 4.60 % | 177 265 | 5.96 % | 175 609 | 4.98 % | 222 028 | 5.91 % |
| BEL/Pao | 6 689 | 0.23 % | 5 431 | 0.24 % | 9 471 | 0.28 % | 11 330 | 0.27 % |
| Ty1/Copia | 4 200 | 0.05 % | 148 | 0.00 % | 6 571 | 0.06 % | 3 480 | 0.03 % |
| Gypsy/DIRS1 | 125 677 | 4.10 % | 169 153 | 5.68 % | 155 980 | 4.61 % | 198 078 | 5.52 % |
| Retroviral | 1 721 | 0.02 % | 1 959 | 0.02 % | 809 | 0.01 % | 2 575 | 0.03 % |
| DNA transposons (class II): | 224 170 | 3.49 % | 349 135 | 4.94 % | 303 130 | 4.03 % | 286 973 | 3.92 % |
| TIR elements: |  |  |  |  |  |  |  |  |
| hobo-Activator | 100 154 | 1.39 % | 212 903 | 2.82 % | 143 436 | 1.65 % | 133 077 | 1.49 % |
| Tc1-IS630-Pogo | 38 954 | 0.60 % | 54 536 | 0.88 % | 49 829 | 0.65 % | 45 241 | 0.61 % |
| PiggyBac | 1 478 | 0.03 % | 2 492 | 0.04 % | 1 580 | 0.02 % | 2 548 | 0.07 % |
| Tourist/Harbinger | 9 998 | 0.14 % | 9 633 | 0.10 % | 14 154 | 0.14 % | 18 704 | 0.36 % |
| Other (Mirage, P-element, Transib) | 26 806 | 0.55 % | 7 947 | 0.14 % | 25 637 | 0.45 % | 30 809 | 0.39 % |
| Rolling-circles (Helitrons): | 29 289 | 0.39 % | 57 465 | 0.72 % | 75 418 | 0.51 % | 43 503 | 0.35 % |
| Unclassified: | 3 912 196 | 40.68 % | 4 588 330 | 46.42 % | 4 704 132 | 44.66 % | 4 738 687 | 43.12 % |
| Total interspersed repeats: |  | 55.91 % |  | 66.31 % |  | 62.99 % |  | 61.49 % |
| Small RNA: | 38 630 | 0.37 % | 36 622 | 0.37 % | 50 953 | 0.54 % | 32 867 | 0.31 % |
| Satellites: | 0 | 0.00 % | 4 420 | 3.00 % | 3 946 | 0.04 % | 5 945 | 0.05 % |
| Simple repeats: | 319 274 | 0.55 % | 250 286 | 0.41 % | 17 318 | 0.06 % | 16 351 | 0.05 % |
| Low complexity: | 35 265 | 0.07 % | 21 427 | 0.04 % | 0 | 0.00 % | 0 | 0.00 % |
| Total | 5 026 081 | 57.30 % | 5 871 508 | 67.88 % | 5 761 216 | 64.14 % | 5 802 151 | 62.25 % |

Interestingly, Bov-B LINE retrotransposons were found in our tick genomes: 160 elements in *I. persulcatus*, 155 in *I. ricinus*, 1 in *I. pacificus*, and none in *I. hexagonus* (see Discussion).

### Macro-syntenies between hard ticks

The genomes of *I. ricinus* and *I. scapularis* were found to be largely syntenic (Fig. 1 B). Comparison of these two tick species suggests very little gene shuffling (homologous genes remained in the same blocks). However, the length of the largest scaffolds (chromosomes) varied between the two species and their ranking in size was slightly different. These differences may be due to different amounts of repeated elements, and/or to the state of assembly of these elements. We note that the sequence NW_0240609873.1 representing the 15th largest scaffold in *I. scapularis* was included in scaffold 7 of *I. ricinus*. Conversely, scaffold 15 from *I. ricinus* matched with NW_0240609883.1, the 8th largest scaffold from *I. scapularis*). In both species, the smaller scaffolds that were not included in the 14 putative chromosomes were relatively gene poor and could correspond to regions with a high proportion of repeated elements, which are difficult to assemble. The plot comparing the two genome assemblies (supplementary Fig. S2 A) indicated several inversions (especially for scaffolds 1 and 6 of *I. ricinus*). It was not possible to determine whether these inversions are real or the result of post-assembly errors. The comparison between *I. ricinus* and *Dermacentor silvarum* also revealed the correspondence of major scaffolds between the two assemblies (supplementary Fig. S2 B). *Dermacentor silvarum* has 11 major scaffolds, which corresponds to its chromosomal number. Despite a low level of micro-synteny, there was a substantial proportion of shared content in the chromosomes, with eight exact matches between chromosomes of these two tick species. In addition, scaffold NC_051157.2 of *D. silvarum* had non-overlapping matches with scaffolds 3 and 14 of *I. ricinus*, *D. silvarum* scaffold NC_051154.1 matched with scaffolds 6 and 12 of *I. ricinus*, and *D. silvarum* scaffold NC_051155.1 matched with scaffolds 7 and 11 of *I. ricinus*. Thus, depending on the ancestral karyotype, there were only three chromosome fission or fusion events in the two branches leading from a common ancestor to the two extant species, after which macro-synteny remained remarkably stable. In the two comparisons (*I. ricinus* versus *I. scapularis* and *I. ricinus* versus *D. silvarum*) we did not observe evidence of multiple regions from one species matching two different regions from the other species. This indirectly suggests that no large-scale supplication events occurred in a common ancestor of ticks.

### Gene families in ticks and the Chelicerata

Analysis of 497,214 protein sequences from 21 species of Chelicerata, including 11 tick species, resulted in 11,331 gene families comprising a total of 332,365 protein sequences (supplementary Table S3). For some gene families, we found unexpected large differences in gene abundance even among closely related tick species. These gene families with differential gene abundance were associated with gene ontology (GO) terms (or domains, result not shown) typically indicating transposable elements. For example, gene family FAM0000061 had 6,896 genes in the spider *Trichonephila clavata* versus 0 genes in some of the Acari genomes, and 1,266 genes in *I. scapularis* but only 113 in *I. ricinus* (supplementary Table S4). Gene annotation strategies may differ among genomes with respect to the masking of repeated regions and hence the putative TEs. For the subsequent analyses on gene family evolution, we therefore masked gene families with typical transposon domains (e.g. reverse transcriptase, transposase, recombination activating gene) and gene families with atypical size variation – families where the number of genes in *I. scapularis* was more than five times that of *I. ricinus*. The resulting distribution of gene families is illustrated in Fig. 2, showing for example that 620 families were present in each of the 21 species analyzed, whereas 139 families were present in all 11 tick species, but in none of the other species.

**Figure 2:**
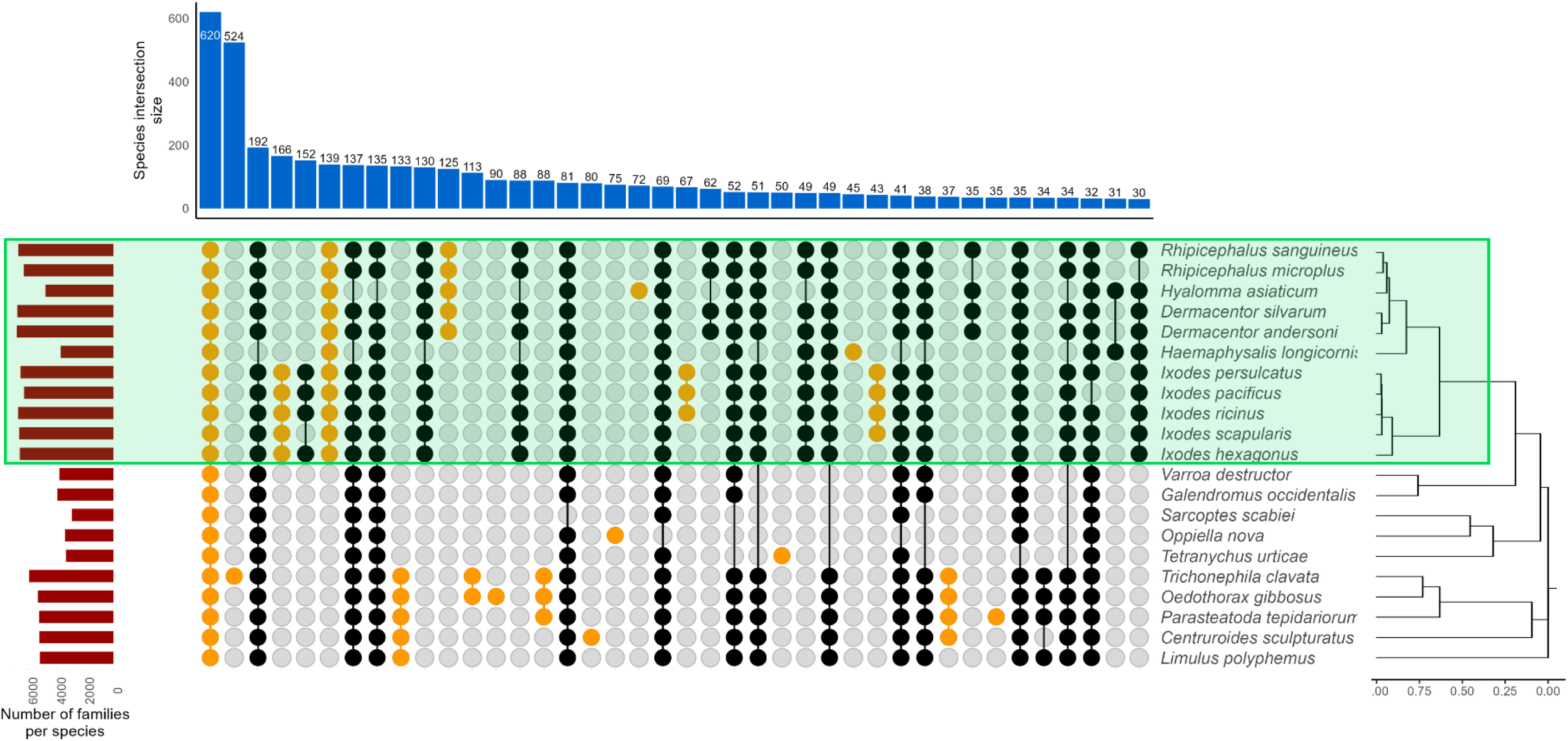
Distribution of gene families among the genomes of twenty-one different species of Chelicerata. Families identified as putative transposable elements were filtered out. The top bar plot represents the number of families shared in a given intersection, the left bar plots gives the number of families per species. Species were ordered according to their phylogeny (right tree) and intersections with a phylogenetic relevance are indicated in orange. Tick (Ixodida) species are highlighted in green

### Phylogeny based on single copy orthologs

Our phylogenetic tree based on 107 single-copy orthologs (Fig. 3) showed high support for the Acari (Parasitiformes and Acariformes) being a monophyletic group. Whether the Acari are a monophyletic group has been debated in the recent literature (Lozano-Fernandez et al. 2019; Zhang et al. 2019; Van Dam et al. 2019). This question, which was not central to our study, will need more complete sequence data to be fully resolved, especially regarding taxon sampling and filtering of sites/genes. The grouping of the Mesostigmata with the Ixodida, and the monophyletic grouping of ticks were both strongly supported, confirming all previous phylogenetic analyses, and this was the main justification for our comparison of gene family dynamics in ticks. Within the Ixodidae (hard ticks), the phylogenetic relationships in our study are consistent with recent studies based on transcriptomes (Charrier et al. 2019) or whole genomes (Jia et al. 2020). Our study confirms that the group of four species belonging to the *Ixodes ricinus* species complex are very close genetically. Within the genus *Ixodes*, an analysis based on a higher number of shared sequences allowed us to obtain a finer resolution of the phylogenetic relationships (supplementary Fig. S3). The unrooted tree showed that *I. ricinus* and *I. persulcatus* are sister clades, as are *I. scapularis* and *I. pacificus*. The first and second species pairs are found in Eurasia and North America, respectively, which suggests a pattern of phylogeographic divergence. We note however that these four species diverged at nearly the same time.

**Figure 3:**
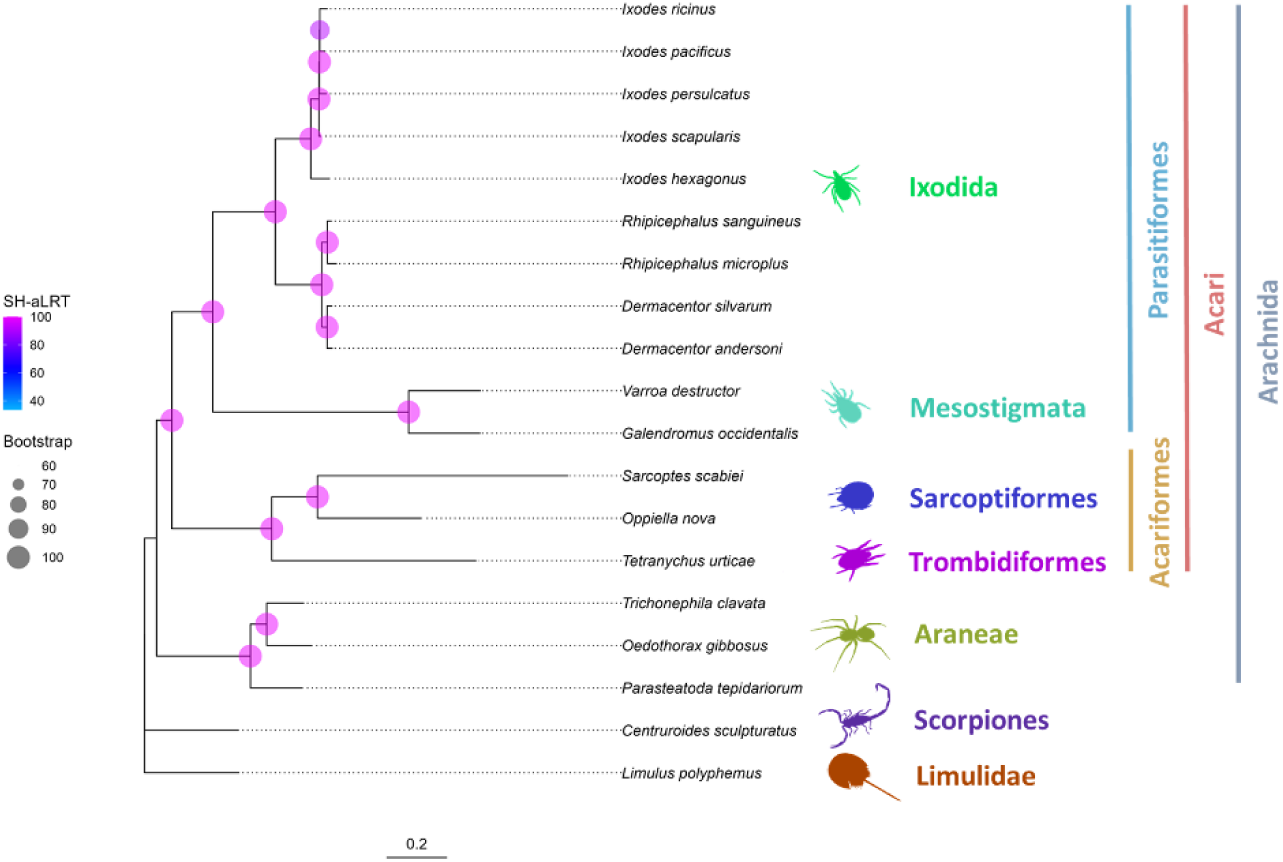
Phylogenetic tree of Chelicerata, based on complete genomes. This analysis was restricted to species with high genome completeness (e.g. without *Hya. marginatum* and *Hae. longicornis*). The tree was built by IQ-TREE 2 using a concatenation of 107 single-copy protein sequences, shared by all represented species. Branch support is shown by bootstrap values and Shimodaira-Hasegawa approximate likelihood ratio test (SH-aLRT) values.

### Dynamics of gene family expansions in ticks

To analyze gene gain/loss dynamics across Chelicerata species, we ran CAFE5 (v5.0), using a lambda of 0.451 predicted from a first round of Base model. This program filtered gene families not present at the root of the tree, and retained 4,525 gene families, of which 497 gene families were found to be either significantly expanded or contracted during the evolution of the Chelicerata. Gene family expansions and contractions were quantified on each branch of the phylogenetic tree of Chelicerata (Fig. 4 and Fig. 5).

**Figure 4:**
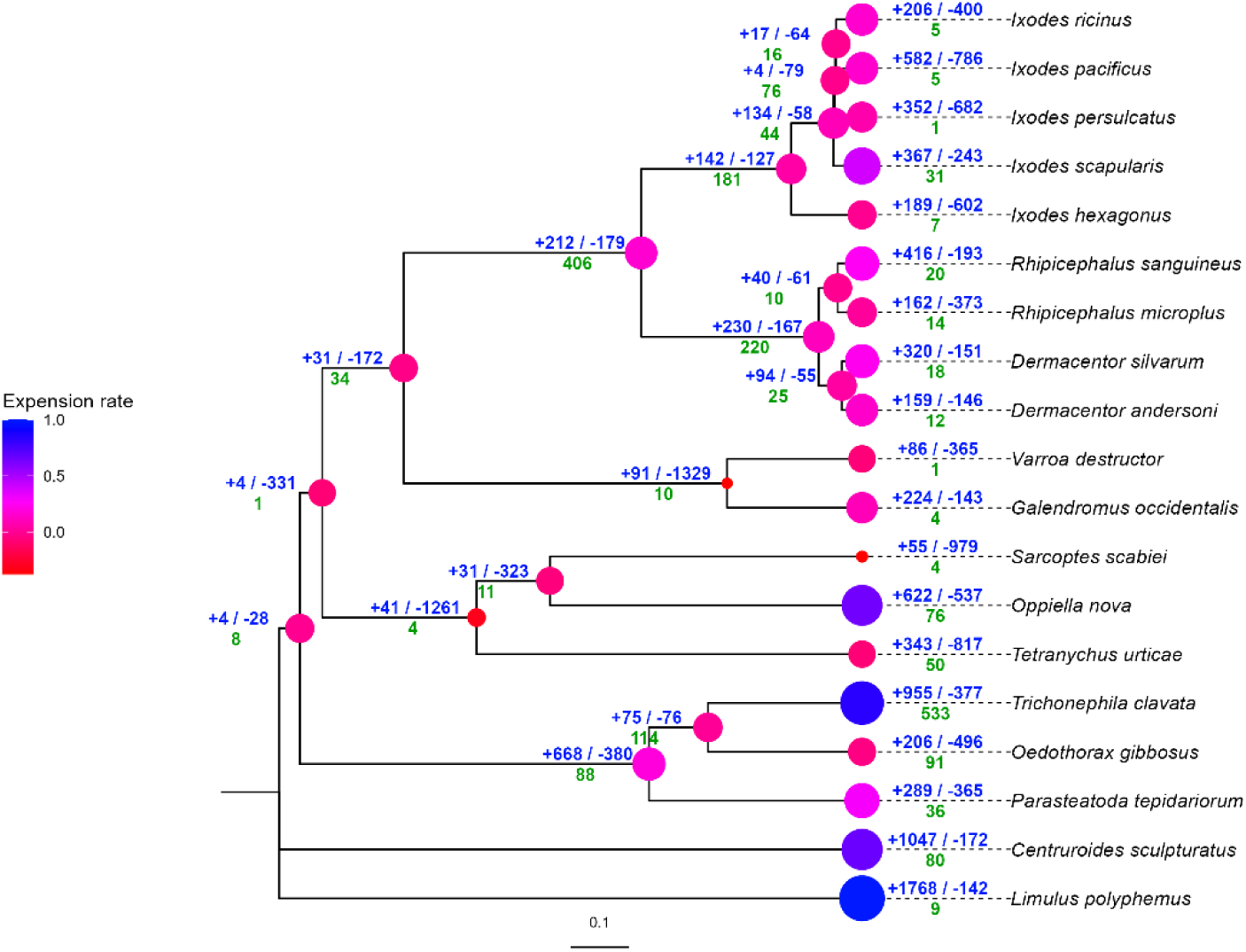
Gene expansion and contraction dynamics in Chelicerata species, analyzed with CAFE. The expansion rate per node is given by the size and the color of the points. The number of expanding (+) or contracting (-) gene families for each node is in blue and above the branches. The number of new families per node is in green. The tree was built by IQ-TREE 2 using 107 protein sequences, before being transformed into an ultrametric tree using phytools and ape R packages.

**Figure 5:**
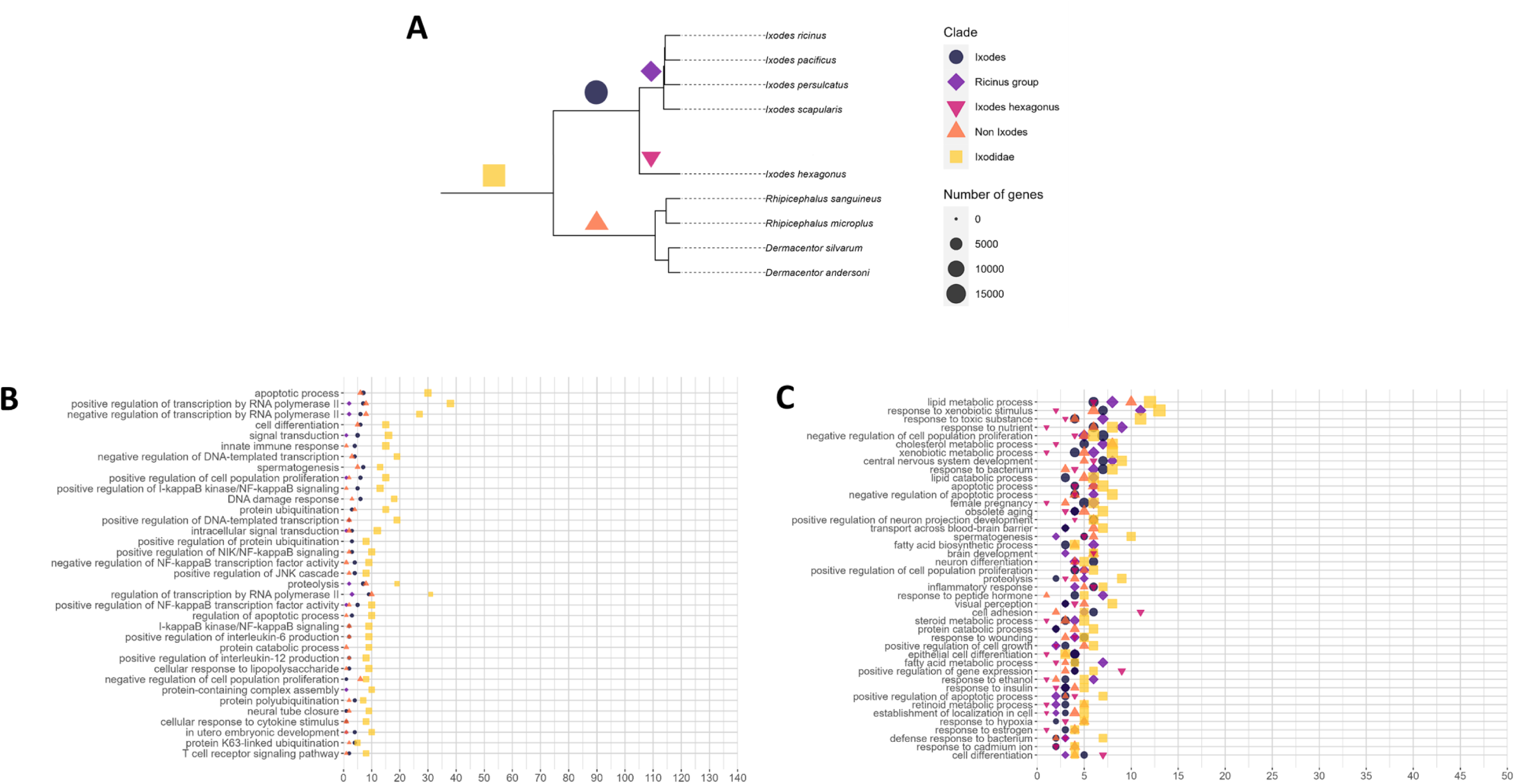
Enriched gene ontology terms (GOs) in gained and expanded families during the evolution of ticks. **A** phylogenetic tree of the tick species (extracted from the complete phylogenetic tree of Chelicerata). The “non *Ixodidae*” clade refers to the Metastriata species. The “ricinus group” is a group of closely related *Ixodes* species. **B** and **C** show the most represented Gene Ontology terms associated with biological processes in gained and expanded families, respectively.

Expanded gene families in the *Ixodidae* included were involved in lipid metabolism and xenobiotic detoxification (Fig. 5 C). The principal GO terms for molecular functions associated with tick expansions were heme binding, transferase, hydrolase, oxidoreductase, metalloendopeptidase/protease and transmembrane transporter activities. The list of the significantly expanded gene families in ticks (supplementary Table S5) shows that some of the most important expansions during the evolution of ticks concern genes associated with detoxification processes, for example cytosolic sulfotransferases (SULTs), carboxylesterases, and Glutathione S-transferases (see below for more detailed analyses). Other important gene family expansions include genes known for their importance in tick metabolism such as metallopeptidases, or serpins. Other gene family expansions were unexpected and not previously described, such as acylcoA synthases and fatty acid elongases.

The number of gene families gained per branch was also estimated by summing all gene families present in a specific clade but absent from species outside of this clade. We identified 406 gene families that were gained in the common ancestor of ticks (i.e., these gene families were absent in all other Chelicerata, Fig. 5 B). These results must be interpreted with caution; we noted that the largest gene family in this category, FAM007521, contains genes annotated as tumor necrosis factor receptor (TNFR)-associated factors (TRAF), which are widespread in the Metazoa. Tick genes from this family show a low level of conservation since best hits have ∼25% identities in BlastP comparisons with non-tick organisms. This suggests that this gene family contains highly divergent genes and would explain why this gene family did not have any potential orthologs in the Chelicerata.

### Structure of genes, importance of mono-exonic genes

As many as 20.7% of genes in *I. ricinus* were predicted to be mono-exonic. This result is interesting because in eukaryotic genomes mono-exonic genes are usually at much lower frequencies. The percentage of mono-exonic genes was high in gene families tagged as putative TEs (35% on average in families containing > 10 genes in *I. ricinus*), but with a large range (0% to 86%). The percentage of mono-exonic genes was also high in other gene families (18% for families containing > 10 genes in *I. ricinus*), again with a large range (0% to 100%). Some of the gene families with high a percentage of mono-exonic genes corresponded to the most expanded gene families in ticks, such as the fatty acid elongases (FAM002111, 82%), cytosolic sulfotransferases (FAM000226, 52%), serpins (FAM001806, 45%), and M13 metallopeptidases (FAM000666, 34%), suggesting that a proliferation of mono-exonic genes contributed to tick-specific gene family expansions. However, other large gene families had few or no mono-exonic copies, such as the MFS transporters (FAM000149).

### Structural and regulatory non-coding elements

Extensive analysis to identify and annotate putative structural and regulatory non-coding elements in the *I. ricinus* genome revealed a total of 21,792 non-coding RNAs (ncRNAs) and cis-regulatory elements, including small nuclear RNAs, small nucleolar RNAs, ribosomal RNAs, long non-coding RNAs (lncRNAs), transfer RNAs, microRNAs, ribozymes or riboswitches, among others (Table 4). Annotating the lncRNAs was difficult due to the paucity of information on lncRNAs in the species studied here and the lack of previous complete annotation of lncRNAs in *I. ricinus*. To accurately annotate the lncRNAs, we used a published lncRNA dataset that included both putative and consensus lncRNAs for *I. ricinus* (Medina, Abbas, et al. 2022). Consensus lncRNAs from this dataset were considered as high-confidence annotations. Before aligning the putative lncRNAs to our *I. ricinus* genome, we confirmed their non-coding properties. We identified 13,287 lncRNAs with alignment scores above 90, indicating significant matches with the genome. To ensure the reliability of our results, we deleted 2,591 lncRNAs that showed overlap with coding RNAs. As a result, we annotated a total of 10,696 lncRNAs in the *I. ricinus* genome, of which 2,433 were classified as high confidence lncRNAs. Our annotation and characterization of ncRNAs and cis-regulatory elements in the *I. ricinus* genome therefore contributes to a comprehensive view of the structural and regulatory non-coding elements of this species.

**Table 4:** Distribution of structural and regulatory non-coding elements in the genome of *I. ricinus*.

| Non-coding element | Class | Number |
| --- | --- | --- |
| Transfer RNA | Non-coding RNA | 9815 |
| Ribosomal RNA | Non-coding RNA | 267 |
| Small nuclear RNA | Non-coding RNA | 192 |
| Small nucleolar RNA | Non-coding RNA | 21 |
| Binding small RNA | Non-coding RNA | 1 |
| microRNA | Non-coding RNA | 63 |
| Long non-coding RNA | Non-coding RNA | 10696 |
| Signal recognition particle | Non-coding RNA | 16 |
| Ribozyme | Non-coding RNA | 3 |
| Riboswitch | Cis-regulatory element | 35 |
| UTR stem-loop | Cis-regulatory element | 662 |
| Iron response element | Cis-regulatory element | 5 |
| Potassium channel RNA editing signal | Cis-regulatory element | 5 |

### Neuropeptides in the *I. ricinus* genome

A total of 41 different orthologs of invertebrate neuropeptide genes were found in the *I. ricinus* genome (supplementary Table S6). Including protein isoforms, 45 neuropeptide precursors contain more than 100 active peptide forms (supplementary Fig. S4). Most of the neuropeptides identified in *I. ricinus* have clear counterparts in other hard tick species (Donohue et al. 2010; Waldman et al. 2022), whereas agotoxin-like peptide and IDLSRF-like peptide were identified for the first time in ticks in the current study. Interestingly, the precursors of ecdysis triggering hormone, which occurs in the tick *Rhipicephalus microplus* (Waldman et al. 2022) and neuropeptide F found in some hard ticks were not detected in the genome of *I. ricinus*. Similarly, the pigment dispersing factor (PDF), which is common in insects and some Chelicerata, was not identified.

### Tick cystatins and Kunitz-domain proteins

The cystatins are a family of cysteine protease inhibitors. Iristatin has been characterized as a secreted immunomodulator from tick salivary glands found at the tick-host interface (PMID: 30747251, (Kotál et al. 2019)). Our phylogenetic analysis of these gene families (supplementary Fig. S5) found independent expansions of cystatins in *I. scapularis* (ISCP_027970 clade) and *I. hexagonus* (Ihex00005714 clade) and expansions of iristatins in both *I. pacificus* and *I. ricinus*, suggesting that these gene family expansions were important for the biology of these species. In terms of gene structure, the cystatins and iristatins have a typical three-exon structure, and they are principally clustered in two genomic regions on scaffold 3 and scaffold 9, respectively (detailed list provided in supplementary Table S7A). Cystatins correspond to family FAM006825 in the SiLiX analysis, which expanded significantly in the common ancestor of ticks, but not in branches deriving from this node (supplementary Table S5). Iristatins have high expression either in hemocytes, or in ovaries, malpighian tissues, salivary glands, and synganglion respectively (supplementary Fig. S6).

Tick Kunitz-domain proteins constitute a major group of serine protease inhibitors that function in blood feeding (Jmel et al. 2023). Kunitz-type inhibitors are divided into subgroups based on the number of Kunitz domains, which varies from one to five and defines the monolaris, bilaris, trilaris, tetralaris and penthalaris groups, respectively. The numbers of Kunitz-type inhibitors in each category were quite variable (detailed list in supplementary Table S7 B), even among closely related species within the genus *Ixodes* (supplementary Table S7 C). The most and least abundant categories were the monolaris and trilaris, respectively. Several species-specific expansion events in the monolaris family (supplementary Fig. S7 A) were observed in the genomes of *I. scapularis* and *I. ricinus*, and to a lesser extent in the other species. The bilaris family, which is the second most abundant group, and includes two well-studied genes *Boophilin* and *Ixolaris*, showed few species-specific expansions (supplementary Fig. S7 B), with the exception of an expansion of *Ixolaris*-like genes in *I. scapularis*. Ixolaris has been characterized as a tick salivary anticoagulant localized at the tick-host interface (Francischetti et al. 2002; Nazareth et al. 2006; De Paula et al. 2019), and our phylogeny indicates that *I. scapularis* has five similar gene copies in its genome. Phylogenies for the trilaris, tetralaris, and penthalaris groups are available in supplementary Fig. S7 C, D, E). By contrast with other gene groups analyzed in this study, determining groups of orthologs for Kunitz-domain proteins within the genus *Ixodes* was difficult, due to a patchy representation of arthropod species in each gene family, and unequal patterns of gene duplication. This could be explained by highly dynamic evolution of this gene family, and high sequence divergence. Alternatively, incomplete annotation in the different arthropod species could also be a factor, given the structure of these genes; most sequences (especially in the monolaris group) are short with ∼80 residues and have a typical 4-exon structure with some of the exons being very short). For the three species that were not manually curated in our study or for other tick species, annotation might not be complete and correct. Finally, we note that the automatic clustering by SiLiX assigned most of the Kunitz peptides from the five groups into a single family (FAM000015), which significantly expanded in *Ixodes* ticks, but not in their common ancestor. In summary, Kunitz-domain proteins exhibit dynamic evolution in ticks and other groups in the Chelicerata.

### Serpins

Serpins are protease inhibitors involved in the regulation of many physiological processes in vertebrates and invertebrates (Huntington 2011), and even viruses (Spence et al. 2021). In ticks, serpins are salivary components responsible for anti-hemostatic, anti-inflammatory and immunomodulatory effects in the vertebrate host (Abbas et al. 2022). Serpins from *I. ricinus* can be divided into two groups, Iripins (*I. ricinus* serpins) and IRIS-like serpins, which refers to the first described tick serpin IRIS (Leboulle et al. 2002). Iripins bear signal peptides and are secreted from the cell, whereas IRIS-like serpins are most likely intracellular or secreted non-canonically. A total of 61 Iripins and 21 IRIS-like serpin sequences were found in the genome of *I. ricinus*. Gene expression differed among tick tissues and the number of exons ranged from 1 to 7 (supplementary Table S8). Serpins were generally classified in the same SiLiX family (FAM001806), which was significantly expanded in the common tick ancestor. Our phylogenetic analysis found several clades of Iripins and one clade of IRIS genes (Fig. 6 B), in agreement with a previous phylogenetic study (Spence et al. 2021). Several serpins form clusters of closely related genes in the genome, suggesting they have arisen through tandem duplication (e.g., a cluster of 22 Iripins within a region of 600 Kbp on scaffold 14 of the *I. ricinus* genome, whereas most IRIS serpins form a cluster on scaffold 9). Mono-exonic genes were common in the Iripins (53% were intronless), whereas three other Iripins only had two exons, the first one being 5’ untranslated region (UTR) only. This gene structure suggests initial events of retroposition, followed by re-exonization of some genes, in addition to tandem duplication. Some serpins are expressed constitutively, while others are upregulated or downregulated by the blood meal (Fig. 6 A). Many serpin genes, especially mono-exonic ones, had no or negligible levels of gene expression, which could explain why they were not found in transcriptomic studies. By contrast, the most highly expressed serpins, such as Iripin-01, –02, –03, –05 and –08 and IRIS-1, have been previously studied and characterized as anti-hemostatic, anti-inflammatory and/or immunomodulatory proteins (Leboulle et al. 2002; Chmelar et al. 2011; Chlastáková et al. 2021, 2023; Kascakova et al. 2021, 2023; Kotál et al. 2021). High numbers of silent serpins located in clusters on the tick genome suggest high rates of gene duplication and gene recombination for this gene family. Highly dynamic gene families can generate new gene copies that are redundant with older ones and that will ultimately be eliminated by selection. Such a scenario would fit with low or no gene expression, and we tagged 14 of the serpins as potential pseudogenes, based on the absence of expression and their incomplete gene model. However, some of the newly generated serpin copies may undergo mutations, be positively selected, and lead to new serpins with novel functions.

**Figure 6:**
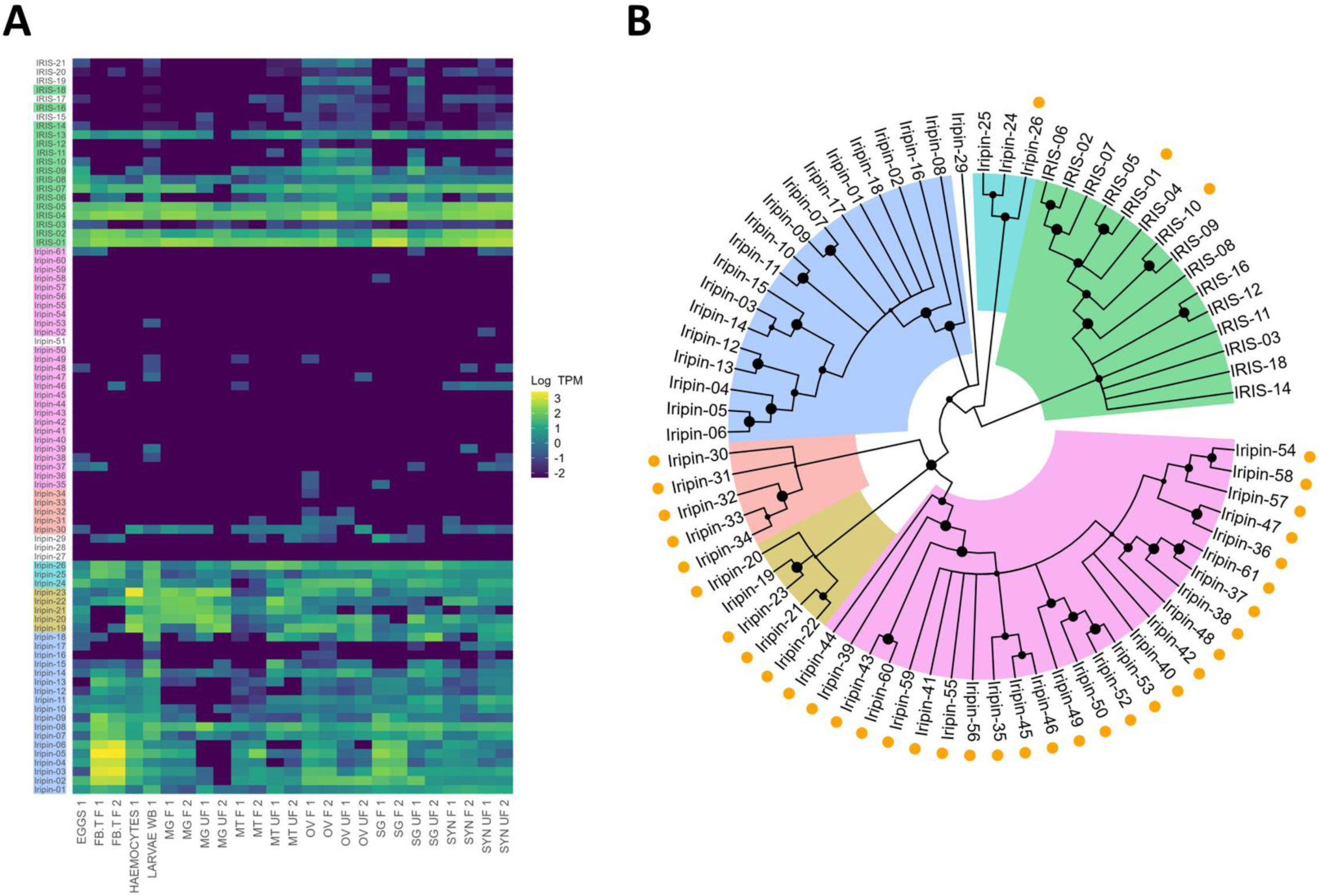
Evolution of serpins in the tick *I. ricinus*. **A** Serpin expression profile. The expression heatmap is based on log10(TPM) (transcripts per million) calculated for the respective transcriptomes: SYN – synganglion; SG – salivary glands; OV – ovary; MT – Malphigian tubules; MG – midgut; FB.T – fat body/trachea; UF – unfed females; F – fully fed females; WB – whole body. **B** Consensus phylogenetic tree of serpins from *I. ricinus*. Sequences were aligned as proteins, signal peptides and variable reactive center loops were removed before the analysis as well as non-informative positions. Edited protein sequences were analyzed by Maximum likelihood method and JTT matrix-based model and bootstrap method with 1000 replications was used to calculate the reliability of tree branches. Only branches with bootstrap value equal or higher than 50% are shown. Mono-exonic serpins are shown with an orange dot. Specific clades are represented by colored areas in the phylogenetic tree, using the same background color for sequence labels in the heatmap.

### Incomplete heme pathways and heme-independent iron inter-tissue trafficking

Heme is an essential molecule for living organisms, involved in multiple processes, and necessary for successful reproduction in ticks (Perner, Sobotka, et al. 2016). The *I. ricinus* genome only contains genes coding for the last three enzymes of the heme biosynthetic pathway: *cpox*, *ppox*, and *fech*, which code for coproporphyrinogen oxidase, protoporphyrinogen oxidase, and ferrochelatase, respectively. Ticks from the Metastriata group have lost the *cpox* gene (Fig. 7, supplementary Fig. S8). Finally, soft ticks in the genus *Ornithodoros* have *cpox*, *ppbox* and *fetch* but also carry the conserved genes *pbgs* and *urod*, which code for porphobilinogen synthase and uroporphyrinogen decarboxylase (supplementary Table S9 and supplementary Fig. S8). The absence of several genes in the heme pathway strongly suggests a loss of heme biosynthetic activity in all ticks, implying they only obtain haem from the blood meal, which agrees with previous studies on *I. ricinus* and other tick species (Perner, Sobotka, et al. 2016; Perner, Provazník, et al. 2016; Perner et al. 2019; Jia et al. 2020). The heme pathway genes *cpox*, *ppox*, and *fech*, have tissue-specific patterns of expression (Perner, Sobotka, et al. 2016), suggesting functional transcripts. The function of these three proteins remains elusive, but the retention of their mitochondrial target sequence indicates a function in mitochondrial biology (Fig. 7). Finally, no sequence homologous to heme oxygenase was found in ticks, nor in other Chelicerata, as shown by the metabolic pathway reconstructed for porphyrin metabolism (supplementary Fig. S8).

**Figure 7:**
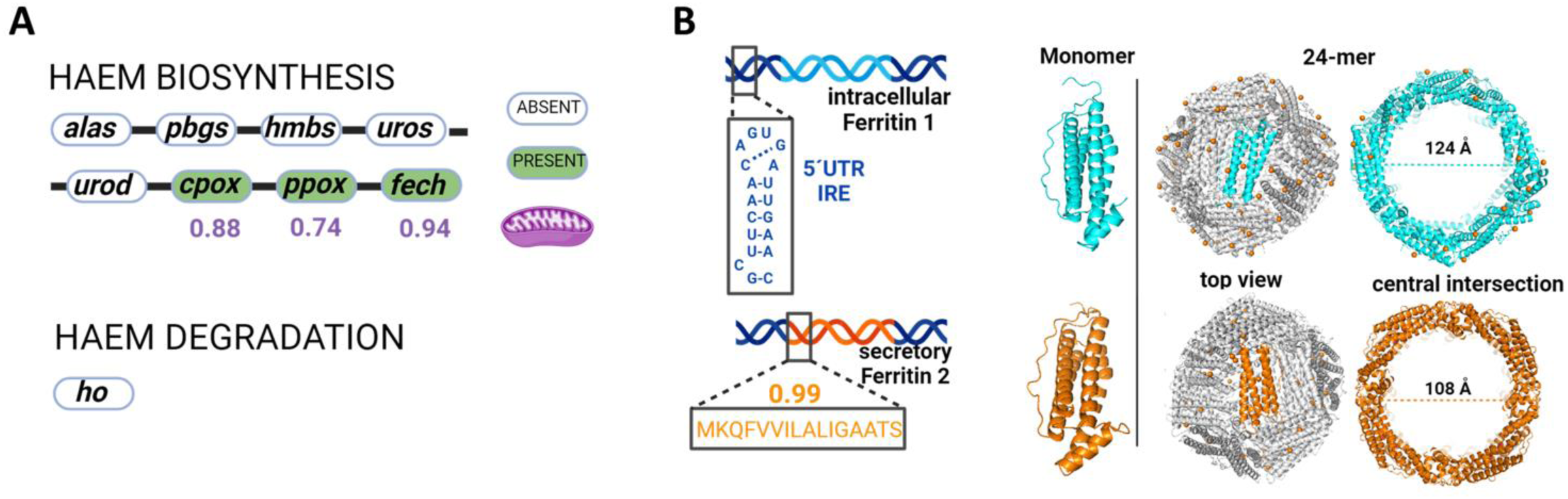
Genomic prediction for heme and iron biology. **A** Gene loss in the heme biosynthesis pathway in the genome of *I. ricinus*. The green color indicates the presence of the homologous gene in the *I. ricinus* genome, with predicted mitochondrial targeting of their protein products (DeepLoc8 prediction values in purple are shown below the enzyme name). **B** Two ferritin genes have been identified in the *I. ricinus* genome: ferritin 1 contains 5’UTR iron-responsive element with the “head” part of the stem-loop structure and complementary bases forming the stem (the blue inset), while ferritin 2 contains a signal peptide (the orange inset) with high SignalP9 probability (shown above the inset). The 3-D reconstruction confirms the conservation of monomeric folding and assembly towards a 24-mer multimers of > 10 nm in external diameter.

Ferritins are an essential actor of iron metabolism in ticks. Unlike in other arthropods, the absence of heme oxygenase in ticks means that iron homeostasis is indeed decoupled from haem homeostasis, and is only ensured by these cytosolic proteins that store iron. Our study confirms previous findings that ticks genomes contain two ferritin genes (Hajdusek et al. 2009) that respectively encode for intracellular Ferritin 1 (without a signal peptide) and secretory Ferritin 2 (with a signal peptide). The sequence of the *ferritin 1* gene is preceded (5’UTR to the gene sequence) by a regulatory region called the iron responsive element (IRE) (Kopáček et al. 2003) (Fig. 7). Both tick ferritins fold into conserved four-helix bundle monomers that self-assemble into higher-order 24-homomeric structures (Fig. 7). The mechanisms of cellular export of tick Ferritin 2 nanocages (Oh & Jung 2023) or their uptake by peripheral tissues remains unknown. The presence of two types of ferritin, allows ticks to store intracellular iron in tissues using Ferritin 1, and traffic non-haem iron between tissues using Ferritin 2 (Perner et al. 2022). Binding of the iron regulatory protein to the IRE prevents expression of Ferritin 1 under low iron conditions (Perner, Sobotka, et al. 2016; Hajdusek et al. 2009). While these ferritin cages are often several nanometers in diameter, the secretory Ferritin 2 was identified within larger (∼ 100 nm) hemolymphatic extracellular vesicles in *Rhipicephalus haemaphysaloides* and *Hya. asiaticum* ticks (Xu et al. 2023).

### Metallopeptidases

The M13 metalloproteases are ubiquitous in bacteria and animals, indicating their evolutionary significance. Mammalian M13 metalloproteases, exemplified by neprilysin, consist of a short N-terminal cytoplasmic domain, a single transmembrane helix, and a larger C-terminal extracellular domain containing the active site. Invertebrate M13 metalloproteases include transmembrane proteins, and secreted soluble proteins (Meyer et al. 2021). Some invertebrate genomes have expanded gene copies for M13 metalloproteases, but most of them are secreted and catalytically inactive (Meyer et al. 2021; Bland et al. 2008). We screened the *I. ricinus* genome to assess the expansion history of the M13 metalloprotease-encoding genes. We identified 88 genes, which is one of the largest recorded expansions of this gene family. The M13 gene family (FAM000666 according to the SiLix clustering) was the second most-expanded gene family in ticks compared to other Chelicerata and was identified as significantly expanded (CAFE analysis) in both the common ancestor of all ticks, and in the common ancestor of the Metastriata. This reflects even larger numbers of gene copies in the non-*Ixodes* species of hard ticks (∼130 in *Rhipicephalus*, and ∼220 in *Dermacentor* versus ∼90 in *Ixodes* species). The genes display diverse exon arrangements (supplementary Fig. S9); most of the multi-exonic genes are expressed in salivary glands or synganglia, whereas most of the mono-exonic genes are transcribed in tick ovaries (supplementary Fig. S9). To predict conservation of protein functions, we searched for protein motifs known to be important for zinc ion binding (HExxH), protein folding (CxxW), and substrate binding (NA(F/K)Y) (Meyer et al. 2021). The very diverse combinations of conserved/substituted residues among the whole collection of sequences present in the *I. ricinus* genome suggests a full spectrum of proteins with diverse functions, including neo-functionalized proteins, possible scavengers and inactive enzymes. The conservation of the zinc-binding motif is always coupled with a secretion signal peptide, and not with transmembrane domains. Also, proteins containing conserved active domains often have low expression in the tick tissues, suggesting that these proteins play discrete functions across multiple tissues. The functions of highly expressed, possibly inactive, M13 homologs in ticks are currently unknown.

### Gustatory receptors and ionotropic receptors for chemosensation

Non-insect arthropods, including ticks, use two major gene families for chemosensation, gustatory receptors (GRs) (Robertson et al. 2003) and ionotropic receptors (IRs) (Rytz et al. 2013). The IRs are involved in the perception of both volatile odorants and tastes (Joseph & Carlson 2015). Ticks possess olfactory receptors that are part of the IR family and gustatory receptors that are part of either the IR family or the distinct GR family. Insects possess a third family of chemoreceptors, odorant receptors (ORs), which probably evolved in ancient insects from GRs (Missbach et al. 2014). While ticks seem to lack homologs of the insect ORs, their IRs and GRs are related to the insect IRs and GRs, respectively (Eyun et al. 2017). Arthropod chemosensory receptors are characterized by high sequence divergence, gene duplication, and gene loss (Robertson et al. 2003; Robertson & Wanner 2006; McBride 2007; McBride & Arguello 2007). This has been well demonstrated in insect groups for which a large number of species genomes have been sequenced and annotated. For this reason, annotation of tick chemoreceptors also requires careful curation, which we performed here for *I. ricinus*.

The *I. ricinus* genome contains 159 IR and 71 GR genes, whereas 125 IR and 63 GR genes have been described for *I. scapularis* (Josek et al. 2018) (Table 5). For *I. ricinus*, 65 of the 90 full-length IRs (72.2%), and 10 of the 51 full-length GRs (19.6%) were intronless. Acari GRs are highly divergent including the most functionally and genetically conserved GRs in insects, such as the receptors that detect sugar, fructose, carbon dioxide (CO2) and bitter taste (Kent & Robertson 2009; Shim et al. 2015; Kumar et al. 2020). Consequently, our phylogenetic studies only included sequences from the closely related *I. scapularis*, and the predatory phytoseid mite, *Galendromus occidentalis*. We identified three clades of GRs in the Acari (Fig. 8 A), one clade was specific to *G. occidentalis* and another to ticks, and both clades showed relatively large group-specific gene expansions. A third clade contained sequences from both mites and ticks, suggesting that it represents more conserved sequences. Several (but not all) of the *I. ricinus* gene copies in this third clade were intronless, suggesting retroposition events. The fact that the *G. occidentalis* homologs are all multi-exonic in this clade suggests that the multi-exonic status was ancestral, and that independent retroposition events occurred repeatedly in *I. ricinus*. For both GRs and IRs, we most often identified co-orthologs between *I. ricinus* and *I. scapularis*, although some expected orthologs were absent, or there were specific amplifications (e.g., there were 6 copies of a gene in *I. ricinus* that was co-orthologous to IscaGR13F in *I. scapularis*). We note that these differences may result from incomplete annotation in either species.

**Figure 8:**
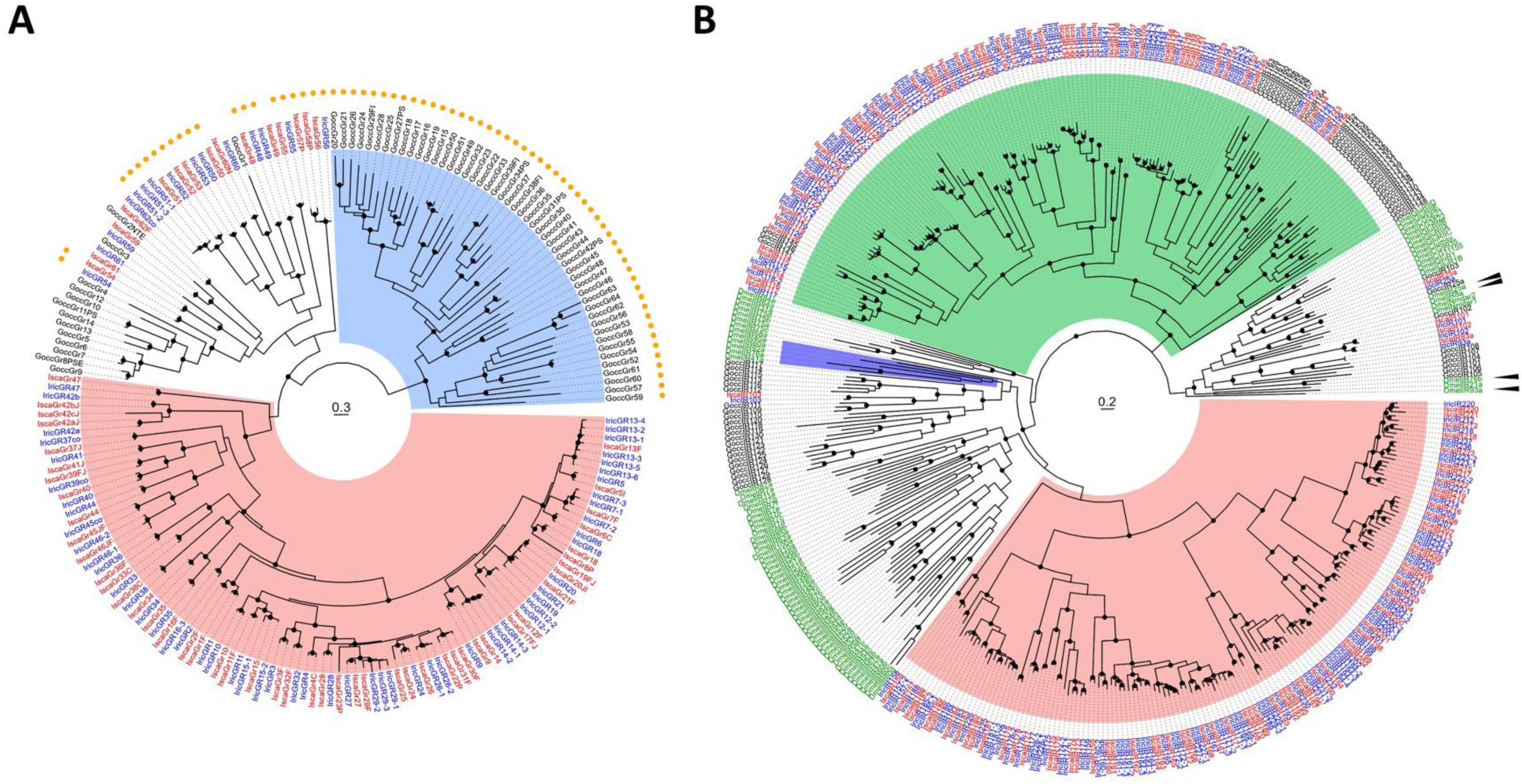
Evolution of tick chemosensory proteins. Trees have been midpoint rooted. Specific clades are represented by colored areas. **A** Maximum-likelihood phylogenetic tree of tick gustatory receptors (GRs). The tree was built using GR repertoires of the ticks *I. ricinus* (IricGRX, labels in blue) and *I. scapularis* (IscaGRX, labels in red), and of the mite *G. occidentalis* (GoccGRX, labels in black). Colored ranges indicate a tick-specific clade (red) and a mite-specific-clade (blue). Clades supported by an aLRT value over 0.9 are indicated by a black dot. Exterior circle: orange dots indicate mono-exonic genes. **B** Maximum-likelihood phylogenetic tree of tick ionotropic receptors (IRs). Color correspondence of labels: *I. ricinus* (IricIRX, in blue), *I. scapularis* (IscaIRX, in red), *G. occidentalis* (GoccIRX, in black) and *Drosophila melanogaster* (DmelX, in green). The ionotropic glutamate receptor clade was used as an outgroup. Black arrowheads show phylogenetic positions of the IR coreceptors found in *D. melanogaster* (IR25a, IR76b, IR8a and IR93a). The clades highlighted in red and green are respectively tick specific and acari specific. The clade highlighted in blue comprises *D. melanogaster* receptors involved in ammonia, amines and humidity detection. Clades supported by an aLRT value over 0.9 are indicated by a black dot.

**Table 5:** Number of chemosensory receptor genes for eight Arthropoda species, including *I. ricinus* (genome sequenced and annotated in this study).

| Class | Species | IRs | GRs | ORs |
| --- | --- | --- | --- | --- |
| Insect | *Drosophila melanogaster* | 63 | 68 | 62 |
| Insect | *Bombyx mori* | 31 | 76 | 49 |
| Insect | *Tribolium castaneum* | 22 | 207 | 259 |
| Insect | *Apis mellifera* | 10 | 10 | 163 |
| Acari | *Ixodes ricinus* | 159 | 71 | 0 |
| Acari | *Ixodes scapularis* | 125 | 63 | 0 |
| Acari | *Galendromus occidentalis* | 65 | 64 | 0 |
| Acari | *Tetranychus urticae* | 18 | 447 | 0 |

The phylogenetic study of IRs also found several clades based on gene structure and conservation (Fig. 8 B). A large clade of sequences was found only in *Ixodes* spp., all of which were intronless. A second large clade contained sequences from both the mite *G. occidentalis* and *Ixodes* spp. (all multi-exonic). Another 5 IR sequences of *I. ricinus* were found in clades more conserved between insects and Acari. In a separate phylogenetic analysis, we compared only these ‘conserved’ IRs between insects and Acari (supplementary Fig. S10), which allowed usto identify orthologs in *I. ricinus* for two IR coreceptors, IR25a and IR93a, which are conserved in all arthropods. Interestingly, IR25a and IR93a were co-linear in the genomes of arachnids, such as *I. ricinus*, *I. scapularis*, *G. occidentalis* (mite), and *Argiope bruennichi* (spider), but not in Lepidopteran insects, such as *Spodoptera littoralis* (Meslin et al. 2022) and *Heliconius melpomene* (van Schooten et al. 2016) (not shown). We also found a pair of closely related sequences, IR101 and IR102, which cluster with the IR93a and IR76b clades (being basal to this group of sequences), and could therefore function as co-receptors in ticks. Finally, another sequence, IR103, was distantly related to the *D. melanogaster* IR proteins that detect humidity, heat, ammonia or amines (Min et al. 2013; Hussain et al. 2016; Knecht et al. 2016, 2017). These stimuli are known attractants for hematophagous (blood feeding) arthropods and are potentially important for the biology *of I. ricinu*s. We note that homologs of IR103 in *G. occidentalis* have undergone multiple duplications. Due to their fast evolution, our SiLiX clustering divided the IR and GR families into several gene families, and some of these contained only tick sequences. This was especially the case for the more divergent mono-exonic clades (e.g. FAM013836, FAM015933, FAM021535 for IRs). However, the majority of multi-exonic IR sequences were contained in one gene family (FAM000240), which was classified as significantly expanded in the tick common ancestor. In summary, we found strong evidence of expansion of GR and IR genes in ticks. Detailed lists for GRs and IRs are given in supplementary Tables S10 and S11, respectively.

### Defensins

Defensins are small cationic antimicrobial peptides (AMPs) that are part of the innate immune system in both hard and soft tick species (Kopácek et al. 2010; Wu et al. 2022). A characteristic feature of these secreted effector molecules is their small molecular size of about 4 kDa and the conserved pattern of six cysteine residues forming three intrachain disulfide bridges (Wang & Zhu 2011). Two types of defensin-related genes were considered, prepro-defensins and defensin-like peptides. Prepro-defensins were defined using three criteria for canonical tick defensins: (i) conserved pattern of cysteine residues in the C-terminal part of the molecule, (ii) presence of the furin cleavage motif (R)VRR, and (iii) mature peptide of ∼4 kDa. Sequences that differed in any of these criteria were termed defensin-like peptides. With these criteria, we identified 14 genes encoding prepro-defensins 1-14 (def1-14), and 8 defensin-like peptides (DLP1-8). Most prepro-defensins (def1-12) were located in a cluster (range 28 Mbp) on scaffold 7, while the two remaining sequences (def13-14) were on scaffold 9, and adjacent. For the DLPs, 3 sequences (DLP1-3) were located on scaffold 7, in the same cluster as def1-12, while the other DLPs were located on scaffold 6 (supplementary Table S12). The absence of intron-less copies in this gene family suggests repeated tandem duplications without retroposition events, unlike other gene families. This is also supported by the fact that closely related sequences in our phylogeny are genes that are physically close in the genome as for example DLP4-8 (supplementary Fig. S11). Ortholog groups within *Ixodes* species were not easy to determine from our phylogeny because there were many ‘gaps’ (subclades where one or more of the five *Ixodes* species were missing), which could be explained by either gene loss/expansion or incomplete annotation (an increased risk for these very short peptides, with a three-or four-exon structure). The predicted mature peptides of DLP1 and DLP2 have only four conserved cysteine residues, while *DLP3* encodes a mature defensin with an extended C-terminal portion, and the *DLP8* gene sequence is longer due to an internal insertion (Fig. 9 A). The sequences of DLP4-7 lack furin cleavage motifs, but both canonical defensins and DLPs contain an N-terminal signal peptide, indicating that they are secreted peptides. Defensins def1 and def3 (which have identical amino acid sequences) were by far the most expressed genes in this family, especially in the hemocytes and midgut of fed ticks (Fig. 9 B). Most defensins were more expressed in the hemocytes than in other tick tissues. Exceptions included def2 and def4 (which differ only in their prepro-domains but have identical mature sequences) that were preferentially expressed in the fat body-trachea complex, def13 was more expressed in the salivary glands of fed ticks, DLP3 was more expressed in the ovaries of fed ticks, and DLP1,2 and 7 were more expressed in the synganglion of males.

**Figure 9:**
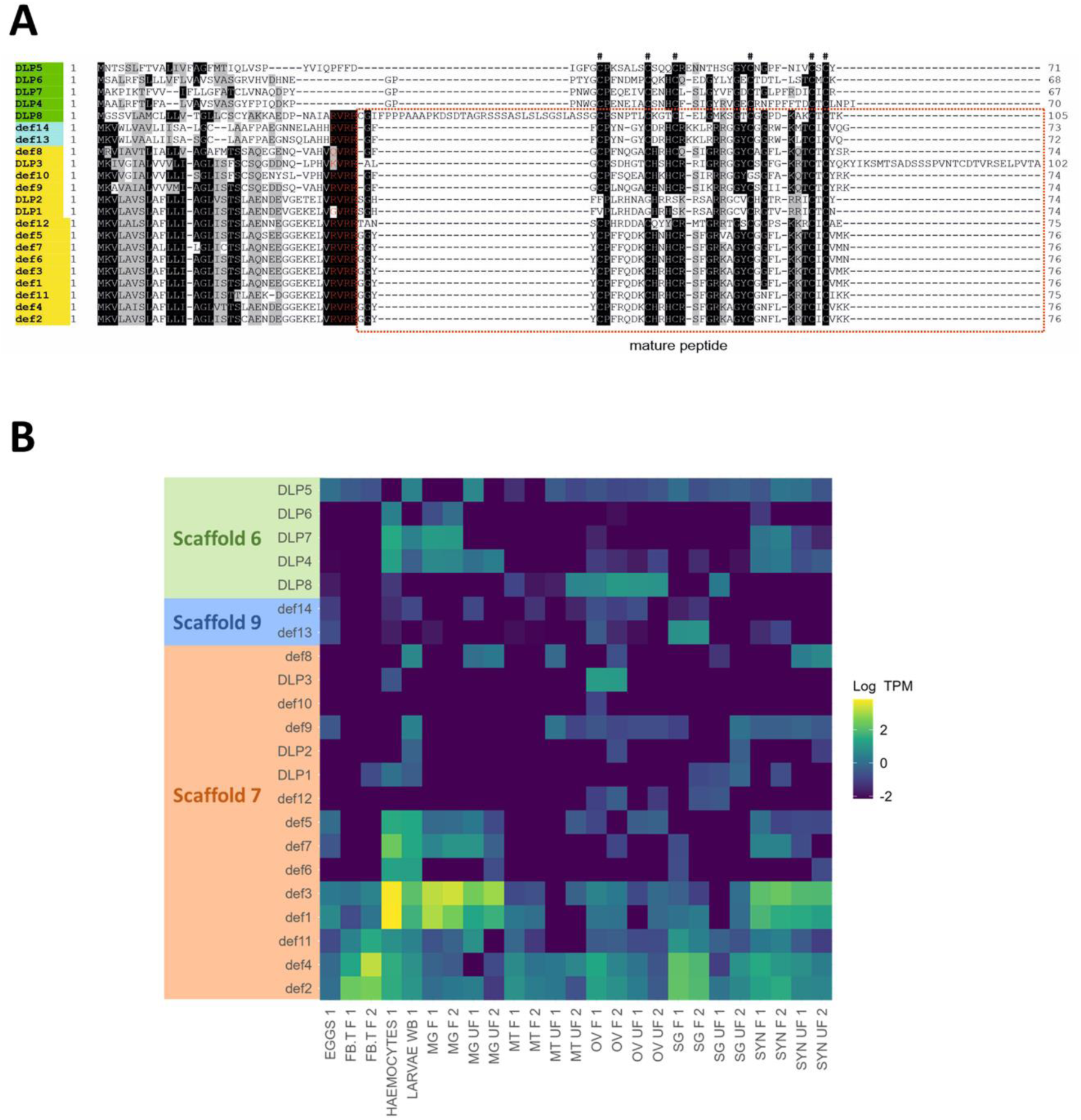
Multiple sequence alignment and tissue expression heatmap of identified *I. ricinus* prepro-defensins and defensin-like peptides. **A** Multiple amino-acid sequence alignment of identified prepro-defensins (def1-def14) and defensin-like peptides (DLP1-DLP8). Highlighted in yellow – genes located on scaffold 7; in blue – genes located on scaffold 9; in green – genes located on scaffold 6; red letters – furin cleavage motif; red dashed frame – predicted mature peptides; # – conserved cysteine residues. **B** Tissue expression heatmap based on TPM (transcripts per million) in respective transcriptomes using log transformation log10(TPM). SYN – synganglion; SG – salivary glands; OV – ovary; MT – Malphigian tubules; MG – midgut; FB.T – fat body/trachea; UF – unfed females; F – fully fed females; WB – whole body.

### Tick gene repertoires for detoxification

Ticks, like other arthropods, have developed a variety of mechanisms to cope with a wide range of endogenous or exogenous compounds. Detoxification processes occur in three main phases (Després et al. 2007). In phase 1, toxic substances are chemically modified to make them more reactive. In phase 2, metabolites are conjugated to hydrophilic molecules to facilitate their excretion. In phase 3, conjugated metabolites are transported out of the cells. Phase 1 typically involves cytochromes P450 (CYPs) and carboxylesterases (CCEs), while phase 2 mainly involves glutathione S-transferases (GSTs), UDP-glycosyltransferases (UGTs) and cytosolic sulfotransferases (SULTs). Phase 3 is carried out by cellular transporters, such as ATP-binding cassette (ABC) transporters, which use the energy from ATP breakdown to transport molecules across lipid membranes. CYPs and CCEs are involved in various physiological processes such as digestion, reproduction, behavioral regulation, hormone biosynthesis, xenobiotic detoxification, and insecticide and acaricide resistance (Oakeshott et al. 2005; Rewitz et al. 2006; Beugnet & Franc 2012; Nauen et al. 2022). Of note, in *I. ricinus*, transcripts for several phase 2 detoxification enzymes were shown to be regulated by ingested heme from the host blood meal (Perner, Provazník, et al. 2016).

The *I. ricinus* genome contains all components of the classical three-phase detoxification system. We did not find homologs for UGTs in the tick genomes, which agrees with the demonstrated loss of this gene family in the common ancestor of the Chelicerata (Ahn et al. 2014). A summary of gene counts is given in Table 6 and detailed lists of genes are given for CYPs, CCEs, GSTs, SULTS, and ABCs (supplementary Tables S13 and S17, respectively).

**Table 6:**
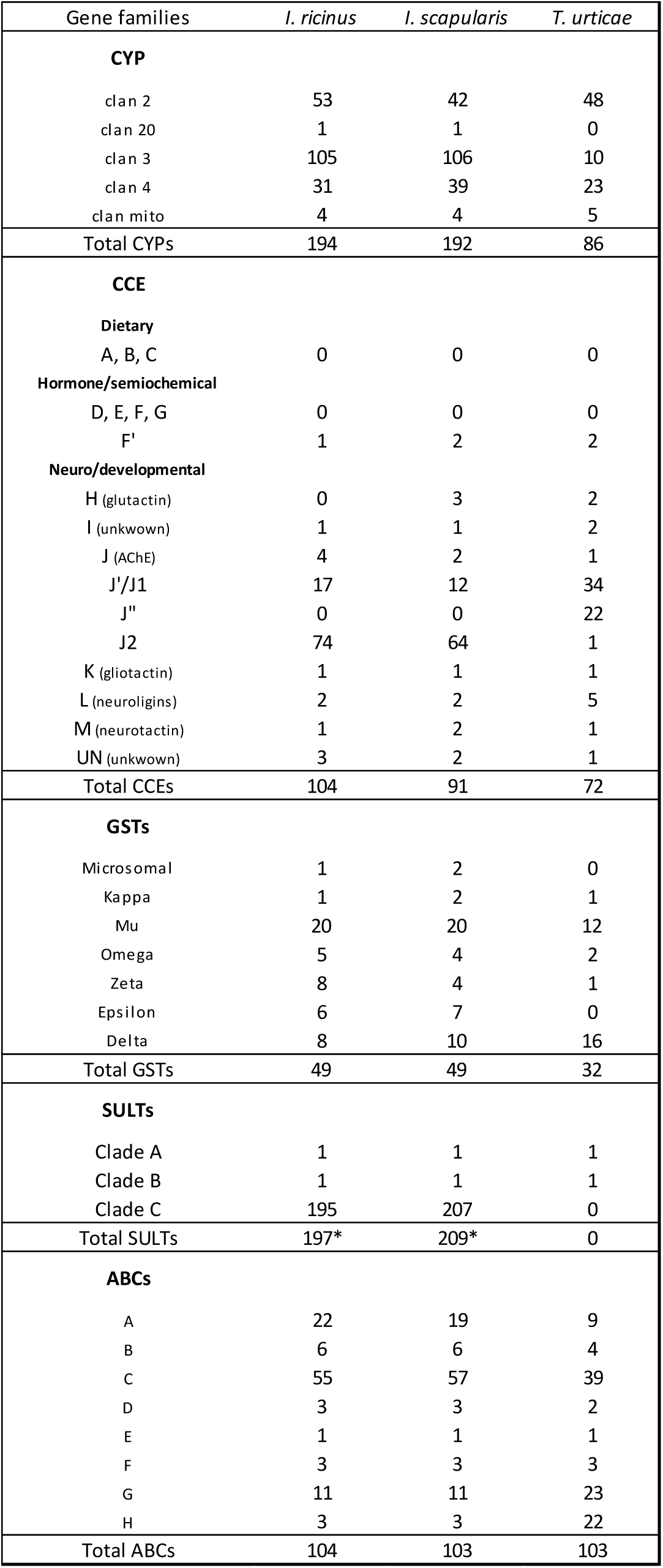
Gene counts for proteins involved in detoxification processes. Counts are given for two tick species (*I. ricinus* and *I. scapularis*) and another Acari (*T. urticae*). For cytosolic sulfotransferases (SULTs), the numbers given correspond to gene counts in the SiLiX family FAM00226 (pre-manual curation).

### Cytochrome P450s

A total of 194 CYP genes were identified in *I. ricinus*, including 131 complete open reading frames (ORFs). The IricCYPs were all named following the Nelson nomenclature (Dr D. Nelson, University of Tennessee Health Science Center, Memphis). The number of IricCYPs and their distribution into five clans is similar to that of *I. scapularis* (Dermauw et al. 2020), in line with the fact that many of the genes are one-to-one orthologs in both species (supplementary Fig. S12). Three members of the mitochondrial clan (CYP302A, CYP314A and CYP315A) involved in ecdysteroid biosynthesis have orthologs in mites and ticks as well as in insects and crustaceans (Dermauw et al. 2020). In mites and ticks, clan 2 genes have been implicated in pesticide detoxification, especially when expressed in the midgut (De Rouck et al. 2023). In *I. ricinus*, four CYP families (3001B, 3001M, 3001N and 3003A) of clan 2 are clustered in tandem duplications, each at different locations, that are also rich in other detoxification gene families (CCEs, SULTs, ABCs). A cluster of 21 CYPs from clan 3 was found on scaffold 13 with genes from CYP3009 (A, B and D) and from CYP3006 (E, F and G), some of which are highly expressed in the midgut of unfed ticks, whereas other groups of genes had high gene expression in other tissues or stages, especially eggs or larvae (supplementary Fig. S13). Six SULT genes were also found at this location on scaffold 13, as well as 2 ABC transporters, suggesting a possible role of this cluster in xenobiotic detoxification. The CYPs essentially corresponded to the SiLiX family FAM000045, which was found to be significantly expanded in several species of ticks, but only in the terminal branches of the evolutionary tree and not in the common ancestor of ticks. This can be explained by other independent large expansions of CYPs in other species of Chelicerata.

### Carboxylesterases

A total of 104 IricCCEs were identified in *I. ricinus*, including 73 complete ORFs. This count is similar to that of *I. scapularis* (Gulia-Nuss et al. 2016; De et al. 2023) indicating that these genes also have one-to-one orthology between the two species. Insect CCEs have been classified into three main functional groups (Claudianos et al. 2006), including the dietary/detoxification group, for which we found no genes in ticks or other Acari (supplementary Table S14, sheet B), and the hormonal/semiochemical group represented by a single gene both in *I. ricinus* and *I. scapularis*. In contrast, the third group of neurodevelopmental CCEs was highly developed in ticks (Fig. 10 A), with a strong expansion of two new clades closely related to AChEs, as already observed in mites (Grbić et al. 2011; Bajda et al. 2015). Similar to the CYPs, we observed a strong differentiation of expression profiles within this gene family and a relatively large number of genes were highly expressed in larvae (supplementary Fig. S14). The CCEs essentially corresponded to the SiLiX family FAM000444, which was significantly expanded in the common ancestor of ticks, and also in the common ancestors of Metastriata and of *Ixodes* species, respectively.

**Figure 10:**
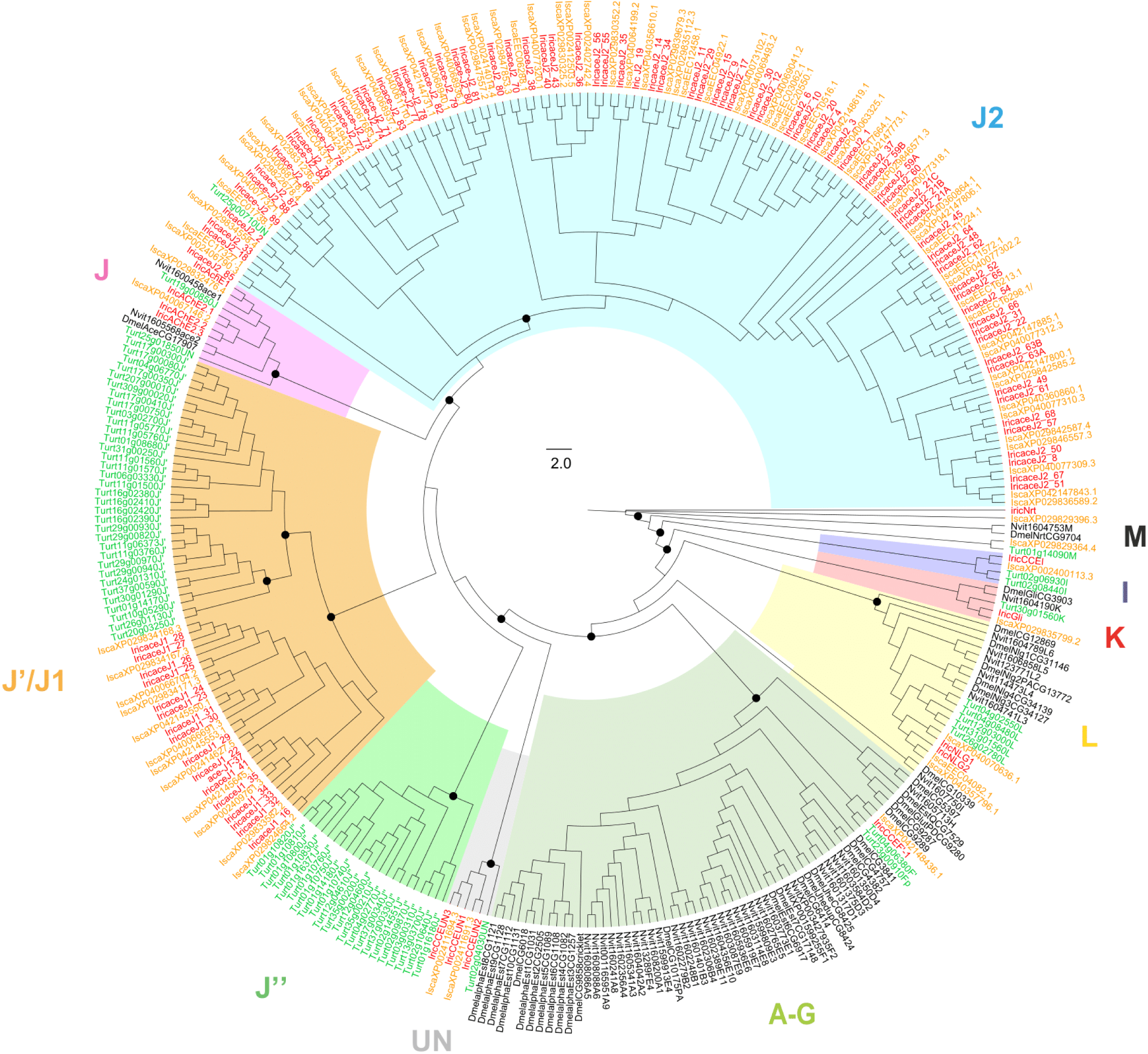
Phylogenetic tree of CCEs in ticks and other representative arthropod genomes. CCEs included arefrom *I. ricinus* (Iric in red), *I. scapularis* (Isca in orange), *Tetranychus urticae* (Turt in green), *Nasonia vitripennis* (Nvit in black) and *D. melanogaster* (Dmel in black). Dots on the tree correspond to bootstrap values above 0.87. Scale is given in the middle of the tree. Ticks show important expansion of the J2 clade.

### Glutathione S-transferases

A total of 49 GST sequences were identified for both *I. ricinus* and *I. scapularis*. Interestingly, more than half of the IricGSTs – including the heme-responsive GST, here named GSTD2/IricT00009278 (Perner et al. 2018) – are located on scaffold 2, a cluster already identified in *I. scapularis* (De et al. 2023). This distribution of GSTs may reflect numerous clade-specific duplication events at the root of GST diversity. Our phylogenetic analysis indicated some patterns of lineage-specific duplication within ticks; for example, several duplications in the Zeta class were specific to the genus *Ixodes*, and several duplications in the Epsilon class were specific to the Metastriata (supplementary Fig. S15 A). Similar to the CYPs and CCEs, several clusters of GSTs were well characterized by distinct gene expression profiles (supplementary Fig. S15 B). The GSTs corresponded almost exactly to FAM000927 in the SiLix clustering, which was found to be significantly expanded in the common ancestor of ticks, and also in the common ancestor of the *I. ricinus* species complex, this GST family had more genes in these four *Ixodes* species compared to *I. hexagonus* or the Metastriata ticks.

### Cytosolic sulfotransferases

The SULT family is one of the most expanded gene families in ticks with ∼200 genes in *I. scapularis* and *I. ricinu*s, and was found to be significantly expanded in both the common ancestor of ticks and in the internal nodes of the tick evolutionary tree (i.e., in the common ancestor of the Metastriata and of *Ixodes* species, respectively, and also in the ancestor of the *I. ricinus* species complex – SiLiX family FAM000226, supplementary Table S5). This suggests a continuous trend of gene family expansion. Our phylogenetic analysis (Fig.11 A) allowed us to distinguish three clades in the SULT gene family. Clades A and B contain sequences found in the Chelicerata and in other Metazoa including humans and houseflies. These two clades have few or no duplications in the Chelicerata. For clade A, only one tick sequence was identified (SFT-142), which was homologous to the four *D. melanogaster* sequences (dmel-St1-4) and to the human ST4A1 sequence. Clade B also contained a single tick sequence (SFT-7), which was homologous to three human sequences (ST1A1, ST2A1 and ST6B1), but no sequences from *D. melanogaster*. Clades A and B have high bootstrap support, so we tentatively propose that an ancient duplication in the common ancestor of arthropods and vertebrates gave rise to these two clades. This would have been followed by secondary gene duplications or gene loss (e.g., loss of the clade B ortholog for *D. melanogaster*). All other SULT sequences form clade C, and are exclusively found in the Chelicerata. Several independent gene expansions occurred within clade C, especially in ticks. While SULT genes in clades A and B have a similar structure and a similar number of exons (6 or 7), the SULT genes in clade C have a much lower number of exons. Most of the SULT genes in clade C are indeed mono-exonic, or have only two exons, the first one often non-coding (i.e., entirely 5’ UTR). In addition, genes with introns are nested between clades of intron-less copies. This suggests an initial retroposition event, which would have generated an intron-less copy, at the origin of clade C – and thus probably in the common ancestor of Chelicerata – followed by a process of re-exonization of some of the copies. A particularly large SULT gene expansion in *I. ricinus* indicates tandem duplication events, as shown by several clusters of adjacent copies. Intriguingly, the conserved multi-exonic copy SFT-7 (clade B) is embedded within a cluster of mono-exonic genes on scaffold 9, although these copies are phylogenetically distant (they belong to clade C) and structurally different. This arrangement is unexpected if it reflects an ancient local duplication event, as tandemly arranged genes are not supposed to arise through retroposition (Pan & Zhang 2008). Finally, the distribution of SULT gene sequences among many different chromosomes or chromosomal regions indicate that additional retroposition events have also occurred.

**Figure 11:**
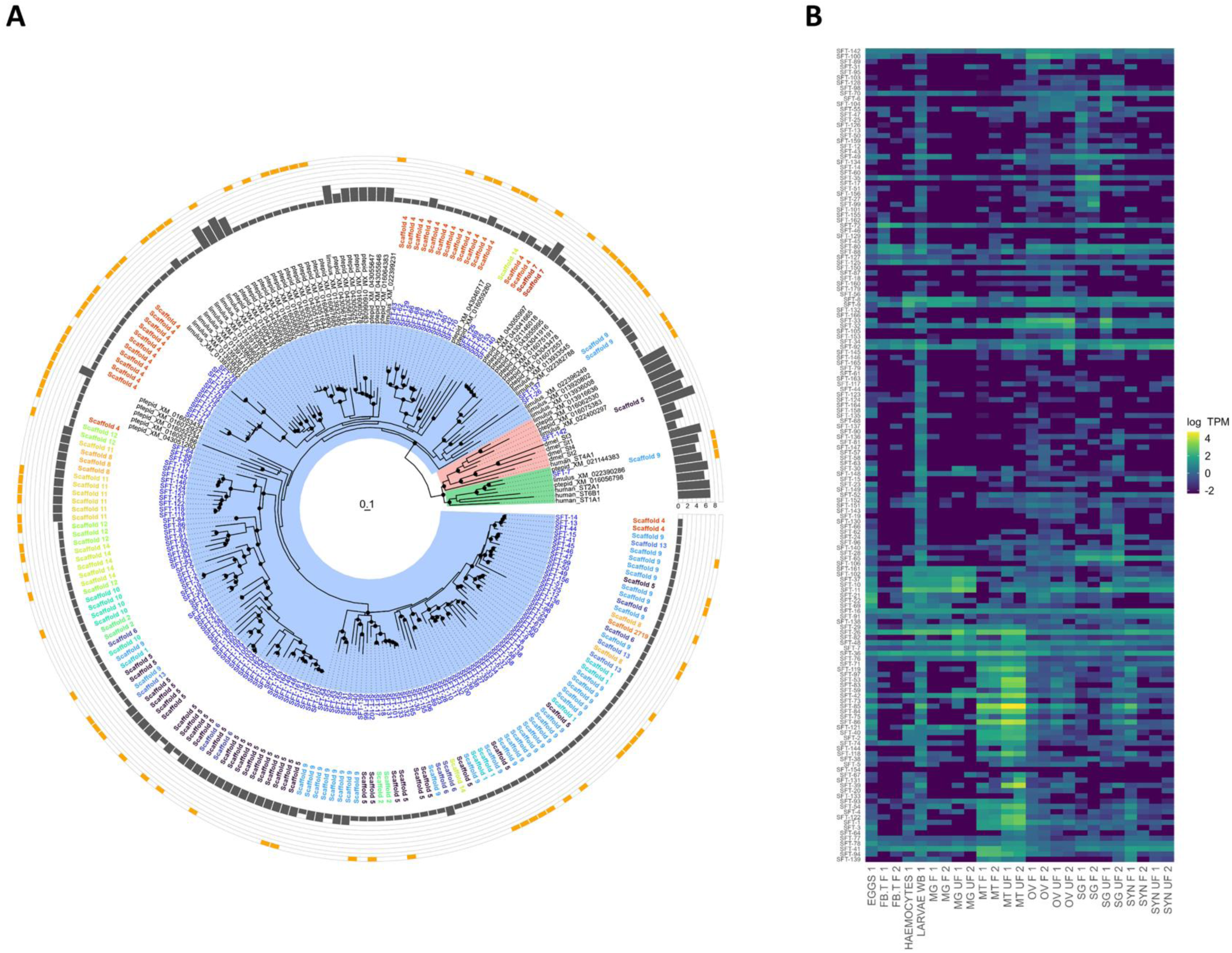
Expansion of the cytosolic sulfotransferases (SULTs) in the the genome of ticks and other Chelicerata, with evidence for both retroposition events and re-exonization. **A** Phylogenetic tree of cytosolic sulfotransferases (SULTs). ML tree using the best-fit LG+I+G4 model of substitution. Sequences from ticks (*I. ricinus*, labels in blue), the spider *Parasteatoda tepidariorum*, the horse-shoe crab *Limulus polyphemus*, human and *D. melanogaster*. Well supported nodes are indicated by dark-filled circles (the width of circles varies with bootstrap values, ranging between 0.85 and 1). The chromosomal (scaffold number) localization is indicated in the first outer circle (each scaffold has its own color label). The next outer circle are bar-charts of the number of coding exons (dark filling) and 5’UTR-only exons (orange filling). The tree allows to define two « conserved » clades (A – green and B – red), with sequences shared between vertebrates and Chelicerata, and a clade (C – blue) with sequences exclusively in Chelicerata. **B** Heatmap of expression of SULTs in *I. ricinus*.

Further evidence that SULTs are a particularly dynamic gene family in tick genomes are the traces of many partial gene copies. However, the absence of complete ORFs or transcript support led us to not annotate these additional hits as *bona fide* genes, and they were not included in our gene counts. With respect to gene expression (Fig.11 B), the conserved copies (SFT-7 and SFT-142) are preferentially expressed in the ovaries of half-fed ticks, whereas many of the other SULTs were dominantly expressed in either the Malpighian tubules of unfed ticks, in larvae, or in salivary glands, with a generally low level of gene expression for the strictly mono-exonic copies.

### ATP-binding cassette transporters

We identified 104 genes encoding IricABCs in *I. ricinus* (the same number as in *I. scapularis*), while 103 genes were found in the red spider mite *Tetranychus urticae* (Dermauw et al. 2013). While most ABC genes in the two *Ixodes* tick species were one-to-one orthologs, the distribution within gene subfamilies was different between ticks and *T. urticae*. ABCs are grouped into 8 subfamilies (A to H) based on similarities in the ATP-binding domain. In *T. urticae*, the most important lineage-specific expansions were found in subfamilies C, G and H, whereas in *I. ricinus* they were found in subfamilies A and C (supplementary Fig. S16). Automatic clustering grouped the B and C subfamilies into a family FAM000065, which was significantly expanded in the genus *Ixodes* and in the common ancestor of the *I. ricinus* species complex, but not in the common ancestor of ticks. This result agrees with the high abundance of ABC genes in other Chelicerata, probably due to independent expansion, and with higher gene counts in *Ixodes* species than in the Metastriata. A second gene family, FAM000381, included the A and G subfamilies, and was found to be significantly expanded in the common ancestor of ticks and in the Metastriata (FAM000381 being more abundant in the Metastriata than the *Ixodes* genus).

Similar to other large gene families, there was strong differentiation in the gene expression profiles of ABCs (supplementary Fig. S17), and the two largest groups of genes were preferentially expressed in eggs and larvae. Mono-exonic copies were found in less than 20 of 104 ABC genes of *I. ricinus*. A group of 11 mono-exonic copies belonging to the C subfamily formed a physical cluster located on scaffold 10, the other 9 intronless copies were dispersed on seven different scaffolds.

## Discussion

### Importance of transposable elements in tick genomes, and definition of cellular genes

Our analysis of transposable elements (TEs) shows that they constitute a large proportion of the genomes in four *Ixodes* tick species, which is typical of large eukaryotic genomes. Interestingly, we identified Bov-B LINEs in the genome of *I. ricinus*, a subclass of retrotransposons abundant in vertebrates, which have also been found in the genomes of reptile ticks, with evidence of horizontal gene transfer between vertebrates and ticks (Walsh et al. 2013; Puinongpo et al. 2020). The presence of Bov-B LINE sequences in several species of *Ixodes* could therefore reflect independent acquisitions of transposons from vertebrate hosts. A recent study of the variation in gene presence-absence in tick genomes showed that most *de novo* genes and genes that are not consistently shared between species are TE-related (Rosani et al. 2023). Due to the unique dynamics of TEs, it is common practice to distinguish TEs from the core genome. However, our comparative study of the Chelicerata genomes found that many of the predicted cellular genes, which sometimes belonged to large gene families, were actually TEs. The number of TEs was high in some genomes, in particular in *I. scapularis* and *Trichonephila clavata* (Joro spider). Following best practices for comparative genomics, we removed these genes from our analyses. Although the role of these TEs is not clear, the acquisition of introns and the high levels of tissue-specific gene expression for some of these genes suggest that they have a functional role in tick biology. More studies are needed to determine whether the large differences in the number of TEs among tick species (including closely related ones) is due to annotation strategies, or to true differences in transposon dynamics.

### Mechanisms of duplication, importance of intronless copies

Gene duplication can result from different mechanisms, with both small-scale events, such as tandem duplications and retropositions, and larger-scale events, such as segmental duplications or whole genome duplication (WGD) (Wolfe & Shields 1997; Kuzmin et al. 2022). In the Chelicerata, previous studies have shown that WGD has occurred independently in the Xiphosura (horseshoe crabs) (Kenny et al. 2016) and in the common ancestor of scorpions and spiders (Schwager et al. 2017; Aase-Remedios et al. 2023), whereas there is currently no evidence of WGD in ticks. A previous study based on the genetic distances between paralogs (Van Zee et al. 2016) suggested that large-scale duplications may have occurred in ancestral ticks. However, the absence of duplicated Hox genes in ticks (Schwager et al. 2017) and our analyses of macro-synteny between tick genomes both suggest that large-scale duplications have not occurred in tick genomes.

Our study identified several large gene families in ticks, often with physical clusters of closely related copies, suggesting that they arose through tandem duplications. Strikingly, many gene families also harbor a high percentage of intronless genes. Intronless copies result from the retroduplication of poly-exonic genes where the mRNA of the ancestral poly-exonic gene is reverse transcribed into DNA and then inserted back elsewhere into the genome (Long et al. 2003; Ohno 1970). Retroposed genes were typically considered as secondary and accidental events, because they generate copies that lack regulatory sequences (and are therefore expected to be eliminated by selection) and because of their small number. However, it has been shown that these retroposed genes can re-acquire new exons, typically in the 5’ untranslated region (UTR), allowing the restoration of a promoter sequence (Fablet et al. 2009; Vinckenbosch et al. 2006). This mechanism could “stabilize” duplicated genes and facilitate their retention in the genome (Micheli & Camilloni 2022). We observed this phenomenon in several gene families, indicating secondary exon acquisition in initially retroposed gene copies. Mono-exonic genes often had low or null levels of expression (with some exceptions), whereas the expression of genes with secondarily acquired introns was in the range of the multi-exonic copies. Thus, our study suggests a high gene duplication rate in tick genomes, driven by a combination of transposon-based retroposition events and tandem duplication. While most duplicate gene copies are probably destined for elimination (e.g., as shown by the many expressionless partial copies for the serpins or SULTs), some copies may be retained by selection and acquire new functions.

### Functional importance of gene expansions

Gene duplication is a major force in the evolution of genomes and in the adaptative potential of organisms (Ohno 1970; Lynch 2002). Although most duplicated genes are eliminated, some new genes undergo sub-functionalization or acquire new functions (Lynch & Conery 2000; Kuzmin et al. 2022). Ticks are no exception and are expected to show specific gene duplications linked with the constraints and particularities of their parasitic life-style. In hematophagous (blood feeding) arthropods, genes involved in host-parasite interactions and blood processing (e.g., salivary gland proteins and proteases) are more likely to show gene expansion (Arcà et al. 2017; Mans et al. 2017; Ruzzante et al. 2019). Ticks possess multigene families involved in various physiological processes, such as detoxification of host molecules, evasion of host immune defenses, and sensory perception (Mans et al. 2017). The co-evolutionary arms race between ticks and their vertebrate hosts exerts particularly strong selection on the tick genes involved in searching for a host, attachment to the host, and blood-feeding. Many of these host-tick interaction genes are expressed in the tick salivary glands, which produce a complex mixture of proteins that have anti-hemostatic, anti-inflammatory, and anti-immunity functions in addition to facilitating tick attachment and tick blood feeding. Thus, the duplication of the tick salivary gland genes contributes to the diversification of these functionally important genes in the frame of the co-evolutionary arms race with their hosts (Chmelař et al. 2016; Francischetti et al. 2008).

Our comparative genomics analysis allowed us to identify important expansions of gene families in tick genomes. Genes associated with detoxification processes showed high expansions in tick genomes (especially in the tick common ancestor) compared to the other Chelicerata. For example, the high number of GST genes identified in tick species may be related to specific adaptations during the rapid evolution of this group towards its modern avian and mammalian hosts (Parola & Raoult 2001), but may also compensate for the absence of UDP-glycosyltransferases in ticks (Ahn et al. 2014). In ticks and mites, several genes of the ABC sub-family C (ABCC) have been shown to confer acaricide resistance (Shakya et al. 2023, Wu et al. 2023). With 55 genes encoding for ABCCs, *I. ricinus* appears to be well equipped to evolve resistance to acaricides, although these genes are probably also involved in physiological processes. For example, the ABC transporter of *R. microplus* (RmABCB10) mediates the transport of dietary acquired heme across cell membranes, and is being studied as an acaricidal target (Lara et al. 2015).

We observed expansions of several gene families that encode metalloproteases, in particular for the M13 metalloprotease family. Metalloproteases are the most abundant protease class in ticks (Ali et al. 2015); they are secreted in tick saliva at the tick host interface (Francischetti et al. 2003; Perner et al. 2020), and are recognised as immunogens inherent to blood feeding (Decrem et al. 2008; Becker et al. 2015; Ali et al. 2015; Jarmey et al. 1995; Perner et al. 2020). However, most of the members of the M13 gene family lack SignalP prediction and only some are expressed by the tick salivary glands. It is therefore tempting to speculate that these proteases play multiple functions including regulation of physiological processes, development, and modulation of bioactive regulatory peptides, and that they are important for the tick parasitic lifestyle.

The most spectacular gene family expansion concerned the family of cytosolic sulfotransferases (SULTs). The research on SULTs is relatively new compared to the research on detoxification enzymes like cytochrome P450s and UGTs (Suiko et al. 2017). The characterization of SULTs in model organisms like humans, zebrafish, and the house fly has shown the importance of these enzymes in the detoxification of exogenous compounds, and in the sulfation of key endogenous compounds. The number of tick SULTs (∼130 genes in Metastriata, and ∼200 genes in *Ixodes*) was considerably higher compared to any other Chelicerata species, vertebrates (∼20 genes), and the housefly (4 genes), suggesting that these genes have acquired an important function in tick biology. As discussed above, this expansion of the SULT genes in ticks appears to result from a combination of retroposition and tandem duplications, as observed in other tick gene families. Our observations of many additional SULT gene fragments (i.e., genes that were not annotated due to partial sequences and no transcription support) shows that SULT gene duplication is highly dynamic, is possibly mediated by nearby transposable elements, and often leads to the degradation of the gene copy. This observation agrees with an analysis of presence-absence variation (PAV) (Rosani et al. 2023) based on tick genomes sequenced in another study (Jia et al. 2020). The study of Rosani et al. identified sulfotransferases among sequences with a strong signal of PAV, along with transposable elements. We hypothesize that with such a dynamic SULT toolbox, ticks may differ at the population level or even at the individual level in the number of SULT copies. Different SULT genes may be adapted to detoxify different substrates, such as the host molecules ingested with the blood. Thus, the function of SULTs could be the digestion and/or detoxification of these vertebrate host blood components. Alternatively, the function of SULTs could be the modification and enhancement of some of the compounds in the tick saliva, creating a cocktail of molecules that modulates the host immune response. We also observedtissue-specific gene expression of SULTs; many SULT genes had higher expression in the Malpighian tubules, an excretory organ that has several functions in ticks, such as the excretion of nitrogenous wastes, osmoregulation, water balance, and detoxification. Other groups of SULT copies were primarily expressed in eggs or larvae, indicating also specialization of SULT gene expression by tick developmental stage.

Ticks do not synthesize juvenile hormone (JH), although JHs and their precursors play crucial roles in molting and reproduction in insects and crustaceans. The last enzyme in the JH pathway, CYP15A1, is absent from tick genomes. It is therefore surprising that a study in *I. scapularis* found a large gene expansion of juvenile hormone acid methyltransferases (JHAMTs), which are the preceding enzymes in the JH pathway (Gulia-Nuss 2016). Another study on *I. scapularis* found that Gln-14 and Trp-120, which are residues critical for interactions with farnesoic acid and JH acid, respectively, were absent (Zhu et al. 2016). Another study found that methyl farnesoate (MF) does not occur in ticks (Neese et al. 2000). Together, these data raise doubts whether these sequences actually function as JHAMTs in ticks. Our study found that this gene family has one of the largest expansions in ticks (with ∼40 genes in Metastriata tick, and ∼80 genes in the *I. ricinus* species complex), although curation showed that a substantial proportion of these gene copies are incomplete and might be non-functional. In contrast with the study of Zhu et al. (2016) our alignment of 45 curated JHAMT sequences in the *I. ricinus* genome showed that Gln-14 and Trp-120 were conserved in the majority of the tick sequences (data not shown). Interestingly, independent expansions of JHAMTs have been recorded in spiders (Yang et al. 2021), and the presence of a transcript of *CYP15A1* may indicate the presence of juvenile hormone and/or methyl farnesoate in this group (Nicewicz et al. 2021). JH synthesis occurs in the corpora allata in insects, and tick synganglia are partly homologous to this tissue (Zhu et al 2016). However, the expression profiles of the JHAMT genes in *I. ricinus* do not show synganglion specificity; most JHAMT genes are more expressed in eggs, larvae, and ovaries, while a few genes have a broad pattern of expression. In conclusion, although ticks lack JH, they have kept most genes of the JH pathway and the large expansion of JHAMTs could indicate that MF is produced by ticks. More studies are needed to elucidate the processes that control molting in ticks, and if tick molting still relies on the JH biosynthesis pathway and on JH precursors.

## Conclusions

In conclusion, our comparative analysis of the genomes of four species within the genus *Ixodes*, namely *I. ricinus*, *I. pacificus*, *I. persulcatus*, and *I. hexagonus*, shed new light on the genomic characteristics of ticks. Through genome sequencing and assembly, and by emphasizing the chromosome-level assembly in *I. ricinus*, we achieved a detailed understanding of the genomic architecture in *Ixodes* ticks. The macro-syntenic analysis highlighted the conservation of genomic organization between *I. ricinus* and *I. scapularis*, with few structural rearrangements. Our annotation efforts, including manual curation for *I. ricinus*, revealed that a high proportion of tick genes have an unusual, intronless structure, indicating frequent retroposition events. We have highlighted the significant role of gene family expansions in the evolution of tick genomes which have undergone highly dynamic gains and losses of genes, alongside expansions and contractions of gene families, showcasing a remarkable adaptation to their parasitic lifestyle. Our comprehensive analysis of the genomes of four *Ixodes* species offers a rich understanding of tick genomics and sets the stage for future functional studies.

## Materials and Methods

### Tick sampling

For *I. ricinus*, we used a laboratory population originally derived from wild ticks in the region of Neuchâtel, Switzerland, and maintained in the laboratory since 1980. This population was maintained in small numbers (∼30 adults) and sexual reproduction was conducted on an annual basis, conditions expected to have favored inbreeding. Unfed adult females were isolated for sequencing. For *I. pacificu*s, ticks were collected from the vegetation on November 9, 2017 in Del Valle Regional Park California (37° 48’ 15.71’’ N, 122° 16’ 16.0’’ W). Adult female ticks were put in RNAlater and shipped to the Nantes laboratory under cold conditions (∼4°C) for DNA extraction. Ticks from the species *I. persulcatus* were sampled in the Tokachi district, Hokkaido, Japan. Adult female ticks were fed on Mongolian gerbils, and their salivary glands (SG) were dissected. *Ixodes hexagonus* ticks were isolated from a live hedgehog collected at Oudon, France (47° 20′ 50″ N, 1° 17′ 09″ W) and sent to a wildlife recovery center (CVFSE, Oniris, Nantes, France). Fully engorged *I. hexagonus* nymphs were collected, and maintained in the lab until they molted into adult ticks.

### DNA extraction

For the four species of *Ixodes*, a single adult female was used for genome sequencing. For *I. persulcatus*, DNA from salivary glands (SGs) was extracted following a standard SDS/ProK and phenol/isopropanol protocol, and stored at –30°C. For the three other tick species, DNA extraction of the whole body was performed following the salting-out protocol recommended for DNA extraction from single insects by 10X Genomics.

### 10X library preparation and sequencing

For each species, linked read sequencing libraries were constructed using the Chromium Gel Bead and Library Kit (10X Genomics, Pleasanton, CA, USA) and the Chromium instrument (10X Genomics) following the manufacturer’s instructions. Prior to DNA library construction, tick DNA fragments were size-selected using the BluePippin pulsed field electrophoresis system (Sage Science, Beverly, MA, USA); size selection was adjusted according to the initial size of the extracted DNA fragments (ranging in size from 20 to 80 Kb). Approximately 1 ng of high molecular weight (HMW) genomic DNA (gDNA) was used as input for Chromium Genome library preparation (v1 and v2 chemistry), which was added on the 10X Chromium Controller to create Gel Bead in-Emulsions (GEMs). The Chromium controller partitions and barcodes each HMW gDNA fragment. The resulting genome GEMs underwent isothermal incubation to generate 10X barcoded amplicons from which an Illumina library was constructed. The resulting 10X barcoded sequencing library was then quantified by Qubit Fluorometric Quantitation; the insert size was checked using an Agilent 2100, and finally quantified by qPCR (Kapa Biosystems, Wilmington, MA, USA). These libraries were sequenced with 150 bp paired-end reads on an Illumina HiSeq 4000 or NovaSeq 6000 instrument (Illumina, San Diego, CA, USA).

### Hi-C library preparation and sequencing

Chicago and Hi-C libraries were prepared by Dovetail Genomics (Dovetail Genomics, Scotts Valley, CA, USA) and sequenced at the Genoscope on a HiSeq 4000 instrument (Illumina, San Diego, CA, USA).

### Illumina short-reads filtering

Short Illumina reads were bioinformatically post-processed as previously described (Aury et al. 2008; Alberti et al. 2017) to filter out low quality data. Low-quality nucleotides (Q < 20) were discarded from both read ends, Illumina sequencing adapters and primer sequences were removed, and only reads ≥ 30 nucleotides were retained. These filtering steps were done using an in-house-designed software based on the FastX package (https://www.genoscope.cns.fr/fastxtend/). Genomic reads were then mapped to the phage phiX genome and aligned reads were identified and discarded using SOAP aligner (Li et al. 2008) (default parameters) and the Enterobacteria phage PhiX174 reference sequence (GenBank: NC_001422.1). Standard metrics for sequencing data are available in supplementary Table S18).

### Genome sizes and heterozygosity rate

Genome sizes and heterozygosity rates were estimated using Genomescope2 (Ranallo-Benavidez et al. 2020) with a kmer size of 31 (supplementary Fig. S17). Genome size ranged between 1.8 Gbp (*I. pacificus*) and 2.6 Gbp (*I. ricinus* and *I. hexagonus*). Heterozygosity rates varied between 0.82% (*I. ricinus*) and 3.17% (*I. pacificus*).

### Tick genome assemblies (10X Genomics) and scaffolding

Genomes were assembled using the Supernova software from 10X Genomics. *I. ricinu*s was assembled using the 1.2.0 version while *I. hexagonus*, *I. pacificus* and *I. persulcatus* were assembled with the 2.1.1 version of the assembler. Hi-C scaffolding of the *I. ricinus* genome assembly was performed by Dovetail using both Chicago and Hi-C libraries. The RagTag (Alonge et al. 2022) software (version 1.0.1) was used to scaffold the genomes of *I. hexagonus*, *I. pacificu*s and *I. persulcatus* by using the chromosome-scale assembly of *I. ricinus* as an anchor. RagTag was launched with default options and with Minimap 2.17 (Li 2018) for the mapping step.

### Gene prediction

The genome assemblies of *I. ricinus*, *I. hexagonus*, *I. pacificus* and *I. persulcatus* were masked using RepeatMasker (http://repeatmasker.org/, default parameters) with Metazoa transposable elements from Repbase (version 20150807 from RepeatMasker package) and RepeatModeler (Flynn et al. 2020) with default parameters (version v2.0.1). The proteomes of *Varroa destructor*, *Centruroides sculpturatus* and *Stegodyphus mimosarum* were used to detect conserved genes in the four tick genome assemblies. In addition, a translated pan-transcriptome of 27 tick species (Charrier et al. 2019) was aligned on the four tick genome assemblies. The proteomes were aligned against genome assemblies in two steps. BLAT (Kent 2002) with default parameters was used to efficiently delineate a genomic region corresponding to the given protein. The best match and matches with a score ≥ 90% of the best match score were retained and alignments were refined using Genewise (Birney et al. 2004) with default parameters, which allows for accurate detection of intron/exon boundaries. Alignments were kept if more than 50% of the length of the protein was aligned to the genome.

We also used RNA-Seq data to allow the prediction of expressed and/or specific genes. Two transcriptome libraries of synganglia (Rispe et al. 2022) from *I. ricinus* (PRJEB40724), a library from the whole body of *I. ricinus* (GFVZ00000000.1), and a pan-transcriptome of 27 different tick species (Charrier et al. 2019) were aligned on the four genome assemblies. As for protein sequences, these transcripts were first aligned with BLAT where the best match (based on the alignment score) was selected. Alignments were then refined within previously identified genomic regions using Est2Genome (Mott 1997) to define intron/exon boundaries. Alignments were retained if more than 80% of the transcript length was aligned to the genome with a minimum percent identity of 95%.

Genes were predicted on the four genome assemblies by integrating protein and transcript alignments as well as *ab initio* predictions using a combiner called Gmove (Dubarry et al. 2016). Single-exon genes with a CDS length smaller or equal to 100 amino acids were filtered out. Additionally, putative transposable elements (TEs) were removed from the predicted genes using three different approaches: (i) genes that contain a TE domain from InterPro; (ii) transposon-like genes detected using TransposonPSI (http://transposonpsi.sourceforge.net/, default parameters); (iii) and genes overlapping repetitive elements. Finally, InterProScan (Jones et al. 2014) (version v5.41-78.0, with default parameters) was used to detect conserved protein domains in predicted genes. Predicted genes without conserved domains and covered by at least 90% of their cumulative exonic length by repeats, or matching TransposonPSI criteria or selected IPR domains, were removed from the gene set.

### Genomic web portal, Apollo server and manual curation

A genomic portal was set up at the BIPAA platform (https://bipaa.genouest.org/), providing access to raw data and to a set of web tools facilitating data exploration and analysis (BLAST application (Camacho et al. 2009), including a JBrowse genome browser (Buels et al. 2016), GeneNoteBook (Holmer et al. 2019)). Automatic functional annotation of genes was performed using Diamond 2.0.13 (Buchfink et al. 2021) against the nr databank (2022-12-11), EggNog-Mapper 2.1.9 (Cantalapiedra et al. 2021), InterProScan 5.59-91.0 (Jones et al. 2014) and Blast2Go 1.5.1 (2021.04 database) (Götz et al. 2008). Results were then made available into GeneNoteBook. The BIPAA Apollo (Dunn et al. 2019) server was also used in order to improve the annotation for *I. ricinus*, based on expert knowledge of several functional groups of gene. Based on the automatic annotation, alignments of RNA-Seq data (reads of selected libraries and contigs of reference transcriptomes), localization of TEs, and of non-coding RNAs, experts were able to perform manual curation of the annotation (gene model edition and functional annotation refinement, including gene naming). Manual curation data was merged with OGS 1.0 (automatic annotation) to produce a new reference annotation named OGS1.1. This merging was performed using ogs_merge (https://github.com/genouest/ogs-tools version 0.1.2). The OGS1.1 was functionally annotated using the same procedure as OGS1.0 and made available into GeneNoteBook.

### Synteny analysis

Synteny analyses between *I. scapularis* and *I. ricinus* were performed using CHRoniCle (January 2015) and SynChro (January 2015) (Drillon et al. 2014), which use protein similarity to determine syntenic blocks across these two genomes. The results were parsed and then plotted into chromosomes using chromoMap R package (v0.4.1 (Anand & Rodriguez Lopez 2022)). Parity plots were built using a homemade R script (R version v4.2.2) based on hit tables, for two comparisons: *I. ricinus* versus *I. scapularis*, and *I. ricinus* versus *D. silvarum*. Hit tables were generated using BlastP (NCBI Blast+ v2.13.0 (Camacho et al. 2009)) with the following options: e-value cutoff = 1e-5, max HSPs = 1 and max target sequences = 1.

### Transposable element annotation

Repeated elements (REs) and transposable elements (TEs) were first annotated by RepeatModeler (v 2.0.2a) (Smit, AFA, Hubley, R. *RepeatModeler Open-1.0*. 2008-2015 http://www.repeatmasker.org) using NCBI BLAST for alignment. Predicted TEs or REs matching with gene sequences from the OGS1.1 prediction were removed from the RepeatModeler results. Finally, a second round of annotation was performed by RepeatMasker (v 4.1.1) (Smit, AFA, Hubley, R & Green, P. *RepeatMasker Open-4.0*. 2013-2015, http://www.repeatmasker.org) using the filtered results of RepeatModeler combined with sequences from arthropods contained in the Dfam database (from RepeatMasker v 4.1.1).

For Bov-B LINEs, sequences annotated as RTE/Bov-B by RepeatModeler were extracted and blasted by tblastx against curated Bov-B LINE sequences contained in the Dfam database (downloaded on 5 December 2023). Only hits with an e-value lower than 1e-3 were kept and sequences aligning with curated Bov-B sequences were considered as BovB LINE retrotransposons.

### Gene clustering, definition of gene families

In addition to the four genomes newly sequenced, the genomes of other tick species and other arachnids were included for a study of phylogeny and evolutionary dynamics of gene repertoires, as listed in supplementary Table S19. Genome completeness was assessed by BUSCO (v5.4.4) for each species (Manni et al. 2021), using the Arachnida database (OrthoDB10) (Kriventseva et al. 2019), with an e-value threshold of 1e-5. For each species, the longest protein isomorph of each gene was extracted using AGAT tools (v0.8.1, https://zenodo.org/records/5834795) in order to retain a single sequence per gene. Three clustering methods were then compared, using several well-annotated gene families as benchmark tests. We found that SiLiX (v1.2.11) (Miele et al. 2011) generated larger gene families that typically matched well the curated gene families, whereas OrthoMCL (v2.0.9) (Li et al. 2003) and OrthoFinder (v2.5.2) (Emms & Kelly 2019) defined gene families at a much smaller grain (clades with fewer genes, of more closely related sequences, as shown for two examples of gene families in supplementary Fig. S18). We therefore decided to use SiLiX for the clustering. Pairwise comparisons between all arthropod species were carried out using BlastP (NCBI Blast+ v2.13.0 (Camacho et al. 2009). Parameters for clustering were a minimum percentage identity of 0.30 and a minimum length of 75 residues. The SiLiX output was then parsed to obtain a contingency table, while a BlastP search against the Swiss-Prot database (12th of January 2023 download), using an e-value threshold of 1e-5, was performed to attribute gene ontology (GO) terms to each gene family.

### Phylogeny of the Chelicerata

The SiLiX output was then parsed to obtain a contingency table, while a BlastP search against the Swiss-Prot database (12th of January 2023 download), using an e-value threshold of 1e-5, was performed to attribute GO terms to each gene family. For gene families containing a single gene sequence in each species, protein sequences were aligned using MAFFT (v7.487 (Kuraku et al. 2013)) with the “genafpair” alignment algorithm and 1000 iterations. The generated alignments were trimmed with TrimAl (v1.4.1 (Capella-Gutiérrez et al. 2009a)), using a gap threshold of 0.6, and concatenated using goalign (v0.3.2 (Lemoine & Gascuel 2021, NAR Genomics and Bioinformatics)). A phylogenetic tree was then generated with IQ-TREE (v2.1.3) (Minh et al. 2020). The best model was identified by the software using BIC and branch supports were estimated with 1000 bootstrap replicates and SH-like approximate likelihood ratio tests. Non-concatenated alignments were also used to construct trees for each family in order to check branch length and topology for all single-copy families. Owing to the low completeness of the *Hae. longicornis* and *Hya. asiaticum* genomes, illustrated by BUSCO metrics and lower number of gene families compared to other ticks (see Results), we removed these species from further analyses. Our phylogeny thus included 107 shared single-copy orthologs, for 19 species of Chelicerata.

To further evaluate the phylogenetic relationships among *Ixodes* species included in our data set, some of them being very closely related, the nucleotide sequences of the single-copy genes of the five species included in our study were aligned using MAFFT (with the “genafpair” alignment algorithm, 1000 iterations and using the LOSUM 80 matrix). Aligned sequences were trimmed with TrimAl (using a gap threshold of 0.8) before being concatenated with goalign. Finally, a phylogenetic tree was built with IQ-TREE. As described previously, the best model was identified by IQ-TREE using BIC and branch supports were estimated with 1000 bootstrap replicates and SH-like approximate likelihood ratio tests.

### Global study of the evolutionary dynamics of gene families

To analyze changes in gene family size across the phylogeny, CAFE5 (v5.0) (Mendes et al. 2021) was run using the previously generated contingency table and the species phylogenetic tree, rooted and previously transformed into an ultrametric tree using phytools (v 1.5-1) (Revell 2012) and the ape R packages (v5.7-1) (Paradis & Schliep 2019) with R version v4.2.2. The horseshoe crab, *Limulus polyphemus*, was used as an outgroup to root the tree. The error model was estimated before running the base model with a dataset cleaned of highly divergent families. The base model predicted a lambda value of 0.451, which was used for a second run of CAFE5 with the full family dataset. This second run was then used to predict gene family expansion/contraction.

### Metabolic pathways

KEGG orthology (KO) numbers were assigned to each protein sequence of the Chelicerata species used in this study using eggNOG-mapper (v2.1.10) (Cantalapiedra et al. 2021). KO numbers of each species were regrouped into five taxonomic groups (*Ixodes* spp., Metastriata ticks, Mesostigmata+Acariformes, spiders and finally a group including *C. sculpturatus* and *L. polyphemus*). Assessment of gene presence/absence was made based on a majority rule in each group (a gene being considered present in each group if present in the majority of its species), to avoid both the effects of incomplete or spurious annotation, or of contamination. Metabolic maps and reactions were then reconstructed using the Reconstruct tool of KEGG Mapper (Kanehisa & Sato 2020). Maps and detailed patterns of presence/absence in each species are available in the BIPAA webpage for the *I. ricinus* genome (see https://bipaa.genouest.org/sp/ixodes_ricinus/download/).

### Expression atlas

Transcriptomic data of *I. ricinus* were downloaded from the NCBI SRA archive (see supplementary Table S20 for detailed information) using the SRAtoolkit (v3.0.0 (https://trace.ncbi.nlm.nih.gov/Traces/sra/sra.cgi?view=software)). A preliminary study of read quality, quantity, and homogeneity between replicates was first conducted, which led us to retain only a sample of the many data sets already published for this species. Whenever possible, we retained two replicates for a given combination of tissue and condition (typically either unfed ticks, or half-replete ticks), to include a minimal level of replication. The transcriptome datasets were filtered and trimmed using Trimmomatic (v0.39) (Bolger et al. 2014). To filter rRNA sequences from the datasets, paired-reads were mapped on an RNA-Seq contig from *I. ricinus* described previously (Charrier et al. 2018); this sequence of 7,065 bp was found to represent a cluster with complete 18S, 5.8S and 28S subunits of rRNA. Mapping was performed with Hisat2 (v2.2.1) (Kim et al. 2019). Unmapped paired-reads (non-rRNA) were extracted using bamUtil (v1.0.14) (Jun et al. 2015) and Samtools (v1.16.1) (Danecek et al. 2021) and read quality was checked using MultiQC (v1.14) (Ewels et al. 2016) on FastQC (v0.11.7) outputs. Another run of Trimmomatic was then performed on the retained reads, which were then mapped on the *I. ricinus* genome with Hisat2. Mapped reads were finally sorted with Samtools and the number of mapped reads per gene was retrieved by the FeatureCount R function contained in the Rsubread package (v3.16) (Liao et al. 2019). Counts were then converted into Transcripts per million (TPMs) – supplementary Table S21. The Spearman correlation heatmap was built for each gene family using the heatmap.2 function (gplots packages v3.1.3) and tree/gene model/heatmap figures were built using a homemade script using treeio (v1.22.0) (Wang et al. 2020), ggtree (v3.6.2) (Yu et al. 2017) and ggplot2 (v3.4.2 ((Wickham 2016) https://ggplot2.tidyverse.org)) packages.

### Annotation of structural and regulatory non-coding elements

To ensure reliable and accurate annotation of structural and regulatory non-coding elements, we used several approaches, software and databases. Initially, we used Infernal and the latest version of the Rfam database to identify ncRNAs and cis-regulatory elements in the *I. ricinus* genome (Nawrocki & Eddy 2013; Kalvari et al. 2021). Subsequently, we used tRNAscan-SE to annotate transfer RNAs in the *I. ricinus* genome (Chan & Lowe 2019) and sRNAbench to identify the most accurate set of miRNAs and their genomic positions (Aparicio-Puerta et al. 2019).

For the annotation of long non-coding RNAs (lncRNAs), we relied on the lncRNA dataset compiled and analyzed by Medina et al. (Medina, Jmel, et al. 2022; Medina, Abbas, et al. 2022). These studies resulted in a consensus set of lncRNAs, which we considered to be high confidence lncRNAs. First, we confirmed the absence of coding properties in these lncRNAs using CPC2 (Kang et al. 2017). Next, we aligned the lncRNAs to the *I. ricinus* genome using Blat and retained alignments with a score above 90. Finally, to eliminate potential assembly artifacts and avoid interference with the set of coding RNAs, we used gffcompare to remove any lncRNAs that overlapped with coding RNAs (Pertea & Pertea 2020).

### Annotation and phylogeny of protease inhibitors in *I. ricinus*

To annotate proteins containing the Kunitz domain (KDCP) and cystatins, we combined *I. ricinus* mRNA sequences from different sources: our initial prediction of coding sequences for the *I. ricinus* genome (version OGS1.0), the National Center for Biotechnology Information (NCBI), and the transcriptome assembled by Medina et al. (Medina, Jmel, et al. 2022). We used TransDecoder to extract coding sequences (CDSs) from the mRNA sequences. To eliminate redundancy, we used CD-HIT (Fu et al. 2012): sequences with a CDS showing 98% identity and at least 70% coverage were identified as redundant, and the longer sequence in each cluster was chosen. Next, InterProScan and BlastP were used to identify proteins belonging to the Cystatin and Kunitz families (Jones et al. 2014; Blum et al. 2021; Altschul et al. 1990). These sequences were then aligned with the *I. ricinus* genome using Blat (Kent 2002). The automatic annotation was then manually curated on the basis of expression and junction data using the Apollo annotation platform (Lee et al. 2013), the result of all our annotations being present in the OGS1.1 version of the genome prediction.

For phylogenetic analysis, we included gene sequences from *I. scapularis* (De et al. 2023), as well as from the four genomes sequenced in the present study. For the three species other than *I. ricinus* sequenced in our study, cystatins and Kunitz domain-containing proteins were annotated using the same method as described for *I. ricinus*, but no manual curation was performed. SignalP was used to identify the signal peptide of cystatins and Kunitz domain-containing proteins (Almagro Armenteros et al. 2019). Clustal Omega was then used to perform a multiple sequence alignment of the mature protein sequence (Sievers et al. 2011). Spurious sequences and misaligned regions from the multiple alignment were removed using trimAl (Capella-Gutiérrez et al. 2009b). The phylogenetic tree was then constructed using FastME (Lefort et al. 2015), and visualized with the R software ggtree (Yu et al. 2017). Detailed lists of KDCPs and cystatins are given in Supplementary Table 7.

### Serpins

The search for serpins in the genome was performed by BLAST, either with blastn algorithm (protein query against translated nucleotide sequences) or tblastn (translated nucleotides query against translated nucleotide sequences), and sequences from the original prediction were manually curated. Sequences were aligned and edited as proteins by using the ClustalW algorithm and Maximum likelihood method and the JTT matrix-based model and bootstrap method with 1000 replications was used to calculate the reliability of tree branches.

### Metallobiology and Ferritins

The search for heme synthesis and degrading enzymes in the *I. ricinus* genome was performed by BLAST, with the tblastn or blastp algorithms, using the sequences of *Dermanyssus gallinae* (Ribeiro et al. 2023) as queries. The Alphafold2 algorithm was used to predict the monomeric structure of ferritin-1 and ferritin-2 identified in the *I. ricinus* genome. The resulting PDB files were used for Swiss Homology Modelling (Waterhouse et al. 2018) to predict the structure of their multimeric assemblies, using human Ferritin heavy chain as a template (3ajo.1.A; seq identity 67%, CMQE 0.89). Measures of the external diameters of the protein multimers were performed in PyMol.

### Chemoreceptors: annotation and phylogenetic study

The annotation of the two major chemosensory gene families of *I. ricinus* was based on known sequences from the closely related species *I. scapularis* (Josek et al. 2018). The *I. ricinus* scaffolds with significant blast hits (cutoff value 1e-30) were retrieved to generate a subset of the genome for each chemosensory gene family. Gene models were obtained after amino acid sequences were aligned to this subset of the genome using Exonerate (Slater & Birney 2005). All gene models thus generated were manually validated or corrected via Apollo. Based on homology with *I. scapularis* sequences, matching parts were joined when located on different scaffolds. The classification of deduced proteins and their integrity were verified using blastp against the nonredundant (NR) GenBank database. When genes were suspected to be split on different scaffolds, protein sequences were merged for further analyses.

Following alignment and phylogenetic proximity, *I. ricinus* sequences were named after their *I. scapularis* orthologs according to Josek et al. 2018. The abbreviations Iric and Isca are used before the gene names to clarify the species, *I. ricinus* and *I. scapularis*, respectively. The gene family was defined by IR or GR for ionotropic or gustatory receptors, respectively and were followed by a number designating a different gene sequence. A supplementary number was given for closely related sequences. For example, if a receptor was named IscaIRX for *I. scapularis* the closest *I. ricinus* receptor was named IricIRX. If other sequences were closely related they were named IricIRX-1, IricIRX-2 etc. The IRs and GRs of the phytoseid predatory mite, *G. occidentalis* (Hoy et al. 2016), were ultimately added to the phylogenetic analysis as well as the *Drosophila melanogaster* IRs. On the GR dendrogram we used the Mocc abbreviations while for the IR dendrogram we did not display the name abbreviations for reasons of clarity. Finally, for the phylogenetic analysis of the conserved IRs, we included the two *Ixodes* spp. (Ir and Is), *G. occidentalis* (Mo)*, D. melanogaster* (Dm)*, Spodoptera littoralis* (Sp) and *Heliconius melpomene* (Hm).

Multiple sequence alignment was performed by MAFFT v7 (Katoh & Standley 2013) and maximum-likelihood phylogenies were built using the Smart Model Selection (Lefort et al. 2015) in PhyML 3.0 (Guindon et al. 2010) (http://www.atgc-montpellier.fr/phyml/) which automatically selects the best substitution model. Node support was estimated using the approximate likelihood ratio test via SH-like aLRT (Anisimova & Gascuel 2006). Trees were retrieved with FigTree v1.4.4 (https://github.com/rambaut/figtree) and images were edited with PowerPoint software.

### Defensins

Identification of genes encoding defensins in the *I. ricinus* genome was performed by tBlastn search (default parameter) using annotated *I. ricinus* prepro-defensin transcripts from a variety of *I. ricinus* transcriptomes deposited in GenBank (NCBI) as queries. Phylogenetic analysis of the prepro-defensins and defensin-like proteins (DLPs) of *I. ricinus* was performed using sequences including other *Ixodes* sp. (*I. scapularis*, *I. persulcatus*, *I. hexagonus*, *I. pacificus*) and constructed as above for Kunitz-type and cystatin protease inhibitors

### Detoxification

We used different sets of tick CYP, CCE, GST, and ABC transporter proteins from published studies (for *I scapularis*) or from the NCBI website automatic annotation databases to search the *I. ricinus* genome using TBLASTN with Galaxy (Giardine et al. 2005), Exonerate and Scipio to align protein sequences to the genome and define intron/exon boundaries (Keller et al. 2008). All generated gene models were manually validated or corrected in WebApollo based on homology with other tick sequences and aligned with RNA-seq data when available. In addition, a direct search was performed using the keyword search on the NCBI website to search for specific protein domains in the databases as well as a search in the automated annotation for *I. ricinus*. The classification of the deduced proteins and their integrity were checked using BlastP against the non-redundant GenBank database. When genes were suspected to be split into different scaffolds, the protein sequences were merged for further analysis. All active sites were confirmed using the NCBI CD search program. Phylogenetic trees were constructed using PhyML (Guindon et al. 2010) based on the best substitution model determined by the SMS server (Lefort et al. 2017), and both SPR (Subtree-Pruning-Regrafting) and NNI (Nearest-Neighbor-Interchange) methods were applied to improve tree topology. Branch supports were estimated by a Bayesian transformation of aLRT (aBayes) (Anisimova et al. 2011). Dendrograms were constructed and edited using the FigTree software (http://tree.bio.ed.ac.uk/software/figtree/). RNA-Seq data, as described on the expression atlas section, were used to construct heatmaps for each enzyme family using TBtools (Chen et al. 2020).

Manual curation of cytosolic sulfo-transferases (SULTs) was carried out for *I. ricinus*, adding several new genes that had not been predicted in the initial automatic annotation, in particular, mono-exonic genes. For a phylogenetic study of this expanded gene family, we included sequences from the horseshoe crab *P. polyphemus* and a spider, *Parasteatoda tepidariorum*, as well as from two model organisms where SULTs have been well characterized, humans and the housefly (sult1-4). For human sequences, we chose one sequence in each of the major clades of SULTs, as previously characterized (Suiko et al. 2017). Sequences assumed to be incomplete based on the predicted gene model and sequence length were discarded. Sequences were aligned with Muscle. The alignment was cleaned with Gblocks, using the following options: –b2=112 –b3=20 –b4=2 –b5=h, implying a low-stringency (small blocks and gaps in up to half of the sequences being allowed). A ML phylogenetic tree was obtained with IQ–tree (Nguyen et al. 2015), using Model Finder to determine the best model of substitution (Kalyaanamoorthy et al. 2017). Branch support was assessed using 1000 ultrafast bootstrap replicates (Hoang et al. 2018) and the resulting tree was graphically edited with ITOL (Letunic & Bork 2019).

### Ethical statement

For the feeding of *I. persulcatus* ticks on gerbils, animal experimentation was carried out according to the Laboratory Animal Control Guidelines of National Institute of Infectious Diseases (institutional permission no. 215038).

## Data availability

Illumina and Hi-C data, assemblies and annotations are available in the European Nucleotide Archive under the following project PRJEB67793. A genome page for each of the four species of this study is available at https://bipaa.genouest.org/sp/ixodes_ricinus/: it contains blast forms, a GeneNoteBook page providing detailed information on each individual gene, a genome browser, and gives the possibility to download sequences, annotation files, expression data (RNA-Seq based atlas, for *I. ricinus*) and maps of metabolic pathways.

## Acknowledgements

We thank Olivier Lambert from the Centre Vétérinaire de la Faune Sauvage et des Écosystèmes (CVFSE) at Oniris, Nantes, France, for providing rescued hedge-hogs, on which we sampled *I. hexagonus* ticks.

Thanks to Joyce Kleinjan (U. Berkeley, USA) for providing *I. pacificus* samples.

Thanks to the Genotoul platform for access to its bioinformatics and calculation resources, and Fabrice Legeai (INRAE, BIPAA) for help with the Apollo database.

We thank Céline Baulard from the Centre National de Recherche en Génomique Humaine (CNRGH) at Evry, France, for providing help with the preparation of 10X libraries.

