## Supplementary material for "Genome sequences of four *Ixodes* species expands understanding of tick evolution": Table and figure legends

Figure S1: K-mer distribution and estimation of genome sizes and heterozygosity rates from short reads for **A** *I. ricinus* **B** *I. hexagonus* **C** *I. persulcatus* **D** *I. pacificus*.

Figure S2: Synteny between the genome assembly of *I. ricinus* and two other tick species. **A** (Left panel) Parity plot between the largest scaffolds of *I. scapularis* (version of (De et al. 2023)) and *I. ricinus* (this study). Inversions may be the result of assembly errors (inversion of blocks) during super-scaffolding. The haploid chromosome number is 14 in both species. **B** (Right panel) Parity plot between the largest scaffolds of *D. silvarum* (Jia et al. 2020) and *I. ricinus* (this study) - haploid chromosome numbers are respectively 11 and 14.

Figure S3: Phylogenetic tree of five species of the genus *Ixodes*, based on their complete genome. The tree was built by IQ-TREE 2 using a concatenation of 2,373 single-copy protein sequences. Branch support is shown by bootstrap values.

Figure S4: Predicted structures of *I. ricinus* neuropeptide precursors. Neuropeptide sequences were predicted and the 45 precursors are presented. Signal peptides are in italic and with a magenta background, while predicted mature neuropeptides are in yellow. Processing (cleavage) sites for mature peptides are shaded in green and putative glycine-derived C-terminal amidation sites in red shade. Cysteines are in bold/red font.

Figure S5: Phylogenetic tree of cystatins in different tick species with a clear duplication event observed for Iristatins shown in dark blue. The tree was constructed using FastME and visualized using the R software ggtree.

Figure S6: Expression heatmap of cystatins genes in *I. ricinus* tissues (salivary glands (SG), Ovaries (OV), Malpighian tubules (MT), midgut (MG), Fat body/trachea (FB_T), and hemocytes) and stages (Eggs, larvae). The suffixes F and UF respectively correspond to half-fed and unfed ticks. To the left: phylogenetic tree of SiLix family FAM006825, comprising most cystatins and iristatins, number of exons, and drawing of the structure of each gene in different exons.

Figure S7: Phylogenetic tree of the Kunitz domain-containing proteins illustrating the monolaris (**A**), bilaris (**B**), trilaris (**C**), tetralaris (**D**) and penthalaris (**E**). Trees were constructed using FastME and visualized using the R software ggtree.

Figure S8: KEGG map of the porphyrin metabolism pathway. The biosynthesis pathway of heme is highlighted by a light orange background (color legend on the top right). Presence/absence profile of metabolic blocks was determined using KO numbers associated to each taxonomic group. Only KO numbers present in the majority of the species composing each group were kept. For example, the gene *cpox* (HemF enzyme) is absent from Metastriata tick species (white background) but present in Ixodes ticks (green background) and also in other groups of Chelicerata. Genes upstream of *cpox* in the pathway are absent from both *Ixodes* and Metastriata.

Figure S9: Evolution of the M13 metalloproteases in the genome of I. *ricinus*. Phylogenetic tree of the *I. ricinus* sequences. Gene structure (number of exons and graphical representation of the gene model). Heatmap of expression (color scale on the right) for different tissues or stages: salivary glands (SG), Ovaries (OV), Malpighian tubules (MT), midgut (MG), Fat body/trachea (FB_T), and hemocytes) and stages (Eggs, larvae). The suffixes F and UF respectively correspond to half-fed and unfed ticks. Presence of a signal peptide (cells with a red background), Number of transmembrane domains (TMDs), Amino acid sequence of conserved motifs required for catalytic activity, if present. On the right: scales for SH-aLRT and Bootstrap tests on the nodes of the phylogenetic tree, and scale of expression level in the heatmap, in Transcripts per Million (TPM).

Figure S10: Phylogenetic tree of the conserved ionotropic receptors between Acari and Insects. The phylogeny was built using five *I. ricinus* (IrIRs, bold black) sequences and their *I. scapularis* (IsIRs, red) orthologs including IR25a and IR93a co-receptor IRs. Homologs from *G. occidentalis* (MoIRs, green) as well as *D. melanogaster* (DmIRs, blue) IR sequences and in two Lepidoptera species, *Spodoptera littoralis* (SlIRs, pink) and *Heliconius melpomene* (HmIRs, purple) were included for comparison. The IR93a clade was used as an outgroup. The two coreceptor clades are pictured as grey colored ranges. Pictograms designate *D. melanogaster* receptors' function in humidity, temperature and sweat molecules such as ammonia and amines. Clades supported by an aLRT value over 0.9 are indicated by a black dot.

Figure S11: Phylogenetic analysis of defensins proteins in Ixodidae ticks. Cladogram illustrating the evolutionary relationships of Defensins proteins in five distinct Ixodidae tick species: *I. ricinus*, *I. scapularis*, *I. persulcatus*, *I. pacificus*, and *I. hexagonus* - respectively colored in dark blue, orange, green, light blue and red.

Figure S12: Phylogenetic tree of Cytochrome P450 genes in I. scapularis and *I. ricinus* genomes. The tree was rooted at the midpoint. Dot size on branches correspond to bootstrap values. Only values above 80 are represented. Gene labels for *I. ricinus* are labelled in red and for *I. scapularis* in black. Clan 2 members are highlighted in light pink, clan 20 in yellow, mitochondrial clan in green, clan 4 in light blue and clan 3 in dark blue.

Figure S13: Heatmap of expression levels for Cytochrome P450 genes in *I. ricinus* (CYPs), based on RNA-Seq data. Tissues or stages were eggs, larvae synganglion, salivary glands (SG), Ovaries (OV), Malpighian tubules (MT), midgut (MG), Fat body/trachea (FB_T), and hemocytes. The suffixes F and UF respectively correspond to half-fed and unfed ticks. The color scale is given with more highly expressed IricCYPs in red and lower expressed IricCYPs in blue. Hierarchical clustering is given on the left side of the heatmap.

Figure S14: Heatmap of expression levels for CCEs in *I. ricinus*, based on RNA-Seq data. Tissues or stages were eggs, larvae synganglion, salivary glands (SG), Ovaries (OV), Malpighian tubules (MT), midgut (MG), Fat body/trachea (FB_T), and hemocytes. The suffixes F and UF respectively correspond to half-fed and unfed ticks. The color scale is given with the most highly expressed genes in red and the least expressed ones in blue. The hierarchical grouping is shown on the top of the heatmap.

Figure S15: **A** Phylogeny and **B** expression heatmap of glutathione S-transferases (GSTs). In the heatmap, tissues or stages are: salivary glands (SG), Ovaries (OV), Malpighian tubules (MT), midgut (MG), Fat body/trachea (FB_T), and hemocytes) and stages (Eggs, larvae). The suffixes F and UF respectively correspond to half-fed and unfed ticks.

Figure S16: Phylogenetic tree of ATP-binding cassette (ABC) transporters.

Figure S17: Heatmap of expression levels for IricABCs. Expression based on RNA-Seq data in eggs, larvae and various tissues of fed and unfed adult females. The color scale is given with the most highly expressed genes in red and the least expressed ones in blue. The hierarchical grouping is shown on the left-hand side of the heatmap.

Figure S18: Comparison among three clustering methods, for two large multigenic families of *I. ricinus*. **A** Acyl-coenzyme A synthetases (ACSs). Gene names as annotated in the *I. ricinus* Apollo database are indicated, and grouped by clades recognised in the literature (Watkins et al. 2007) distinguished by different color ranges. **B** Cytosolic sulfotransferases (SULTs). The outer circles indicate the family ID of each gene as determined by SiLiX (FAMXXX), OrthoMCL (OG1.5_XXX), and OrthoFinder (OGXXX) respectively. This analysis was based on a preliminary run of SiliX run. The subsequent analyses presented in the paper were made on second run, with practically the same results, but different family numbers (e.g. here near all SULTs are clustered in FAM000432, while in the definitive run, they belong to FAM000226).

Table S1: Statistics on predicted genes issued from our automatic prediction pipeline (OGS1.0) and after manual curation (OGS1.1), for the tick *I. ricinus*.

Table S2: Manual curation of the *I. ricinus* gene predictions. Merging statistics for the OGS1.1 prediction, as the result of manual curation: number of genes for each category.

Table S3: Results of a clustering analysis of protein sequences with SiLiX. Columns are species name, total number of gene sequences, number of gene families where a given species is included, and number of gene sequences included in families for this species, respectively before and after filtration of sequences of putative transposable elements (TEs). For each genome, only the longest isoform per gene was retained.

Table S4: Contingency table resulting from protein sequence clustering with SiLiX. The number of genes clustered per family is reported for each species. Gene ontology terms are given in the last column. Families identified as putative transposable elements (TEs) or derived from TEs were counted separately (sheet "putative_TEs") from other families (sheet "non_TEs").

Table S5: List of gene families significantly expanded in the common ancestor of ticks. Columns: Family ID (SiLiX clustering), predicted function, number of genes in each species, GO terms associated with genes of this family, significant expansions at different nodes following CAFE statistical analysis: common ancestor of ticks, ancestor of non-Ixodes ticks (Metastriata), ancestor of *Ixodes* ticks, ancestor of the "ricinus group". Median number of genes respectively in non-ticks (other Chelicerata) and tick species.

Table S6: List of neuropeptides annotated in the genome of the tick *I. ricinus*. Name of the peptide, abbreviation (if existing) and corresponding sequences (cDNA, CDS, protein).

Table S7: Lists of cystatins and Kunitz-domain proteins in the genome of *I. ricinus*. Sheet **A**: Detailed list of cystatins. Sheet **B**: Detailed list of Kunitz-domain proteins. Columns for Sheets A and B: Gene ID, Symbol, Full gene name, Type, and Family assigned through SILIX clustering). Sheet C: Number of known Kunitz-domain proteins (as reviewed in (Jmel et al. 2023)) and number of genes of each class detected in the genomes of I. scapularis, and in the four new genomes sequenced for different species of the genus Ixodes (after this study).

Table S8: List of *I. ricinus* Iripins and IRIS-serpins with names, gene codes, number of exons and their position on scaffolds. Putative pseudogenes are also indicated as well as their family clustering according to SiLix analysis.

Table S9: Heme biosynthesis pathway in the Chelicerata. Gene counts for each gene in the pathway, for the different species included in our whole-genomes comparison. The last column corresponds to the mining of transcriptomic data, for the soft tick Ornithodoros turicata.

Table S10: Detailed list of Gustatory receptors (GRs) in the genome of *I. ricinus*. Gene name, completeness of the gen model, exon number, localization in the genome (scaffold and position), and protein sequence.

Table S11: Detailed list of Ionotropic receptors (IRs) in the genome of *I. ricinus*. Gene name, completeness of the gene model, exon number, localization in the genome (scaffold and position), and protein sequence.

Table S12: Detailed list of genes of the defensin family (prepro-defensins and defensin-like peptides). Genomic location, gene name, Exon number.

Table S13: Detailed list of Cytochrome P450 genes in *I. ricinus*. Nelson name, Size in amino acids, exon number, clan, Scaffold and position, strand.

Table S14: Detailed list of CCEs

Table S15: Detailed list of GSTs

Table S16: Detailed list of SULTs

Table S17: Detailed list of ATP-binding cassette (ABC) transporters.

Table S18: Metrics of sequencing data. Read sizes in base pairs (bp).

Table S19: List of the genome sequences included in our comparative genomics analysis of ticks and other Chelicerata. Columns indicate the species name, taxonomic group, assembly name, accession, size of each genome assembly in Mbp, number of genes, level of assembly, and main reference or project name.

Table S20: List of RNA-Seq datasets obtained for the species *I. ricinus* and used to generate an expression atlas. All data sets retrieved from SRA NCBI archives. The columns contain the NCBI accessions of each data set, technical characteristics of each run, number of spots (one spot is two reads, all data sets being paired), type of isolate (stage, sex and feeding condition of ticks), tissue or organ, and a short sample name.

Table S21: Expression counts for the predicted genes of *I. ricinus*, on a selection of datasets used to generate an expression atlas. Only the longest transcript of each gene was retained. Normalized read counts, in Transcripts per million (TPM). Datasets acronyms correspond to the following: SYN = synganglion pools, SG = salivary gland pools, OV = ovaries pools, MT = Malpighian tubule pools, MG = midgut pools, FB.T = Fat body / trachea pools, UF = unfed, F= fed and WB = whole body.
