## Supplementary figures and images for "Genome sequences of four *Ixodes* species expands understanding of tick evolution"

### Figure S01

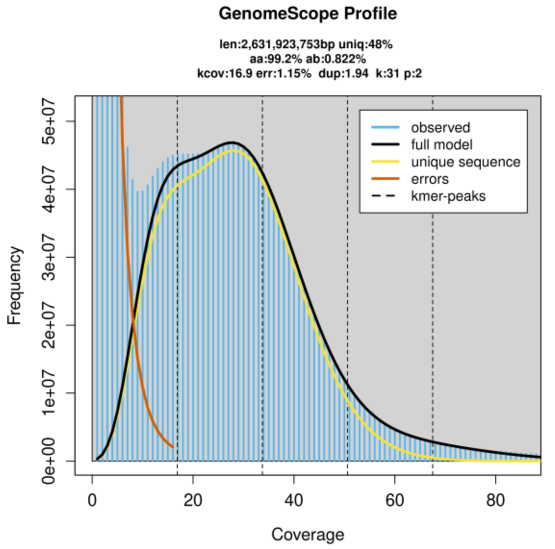

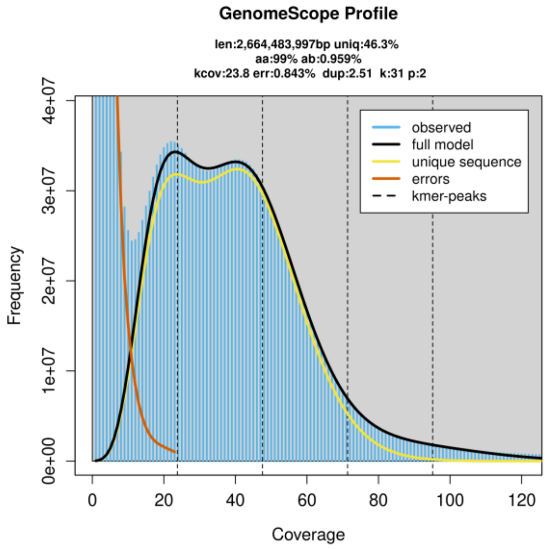

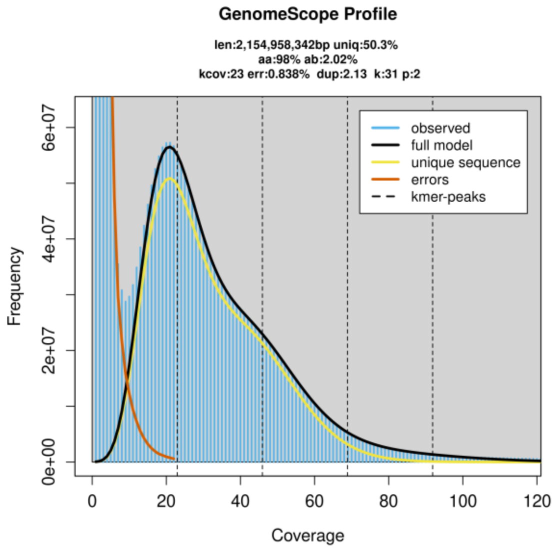

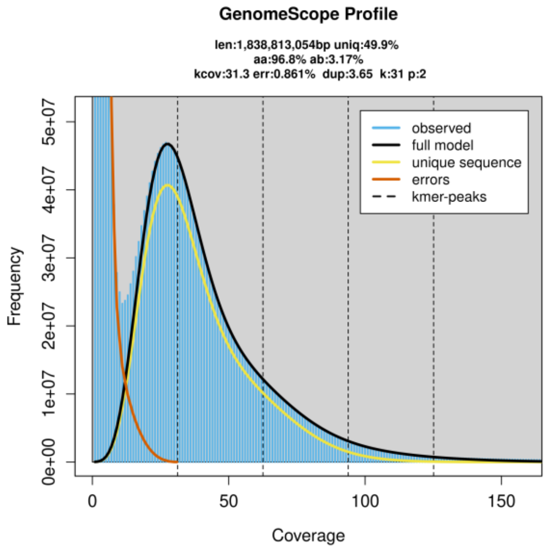

### Figure S02

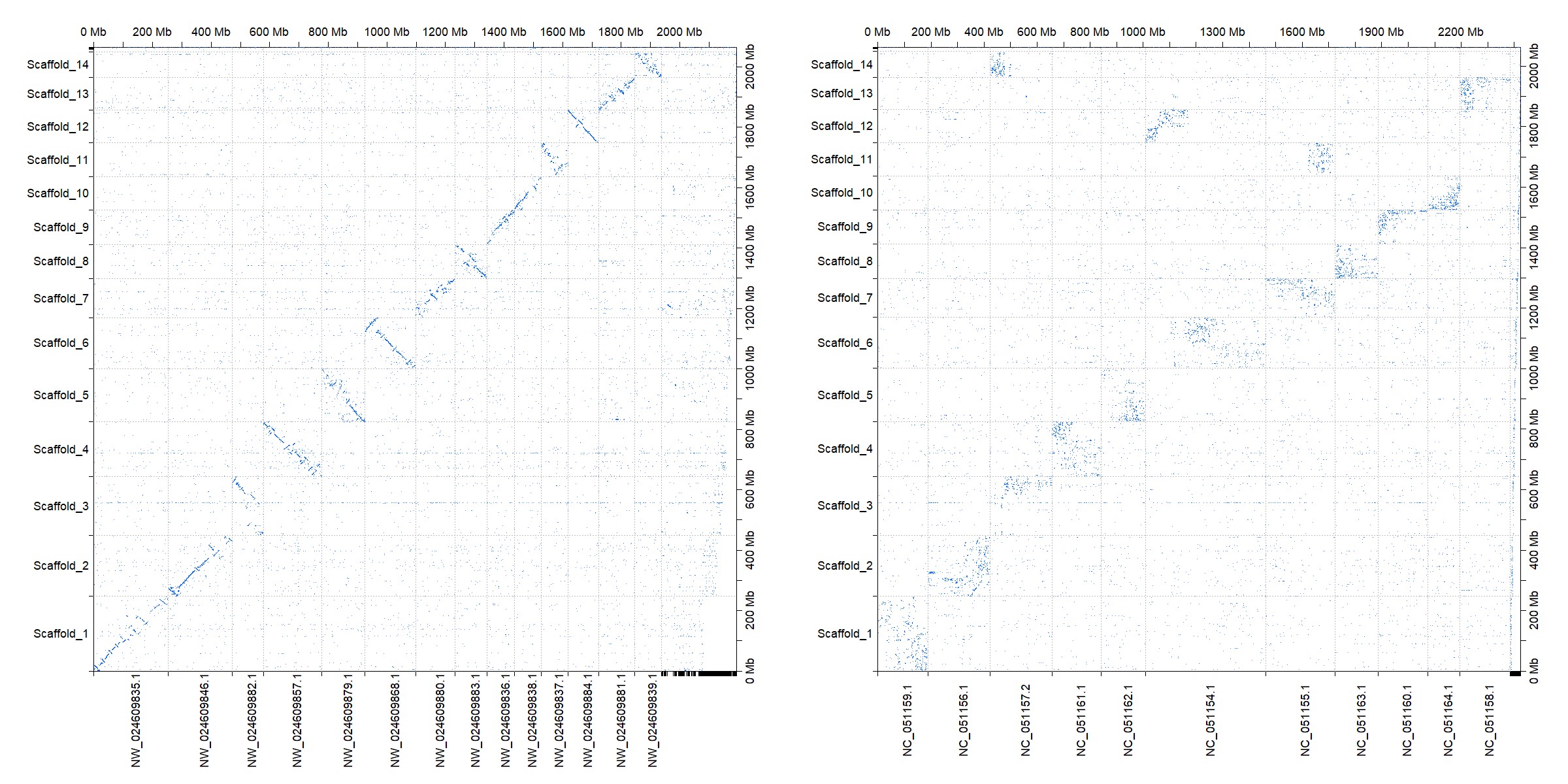

### Figure S03

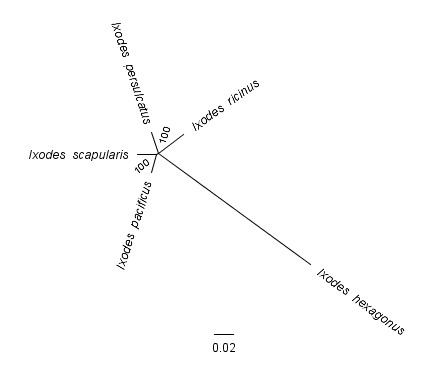

### Figure S05

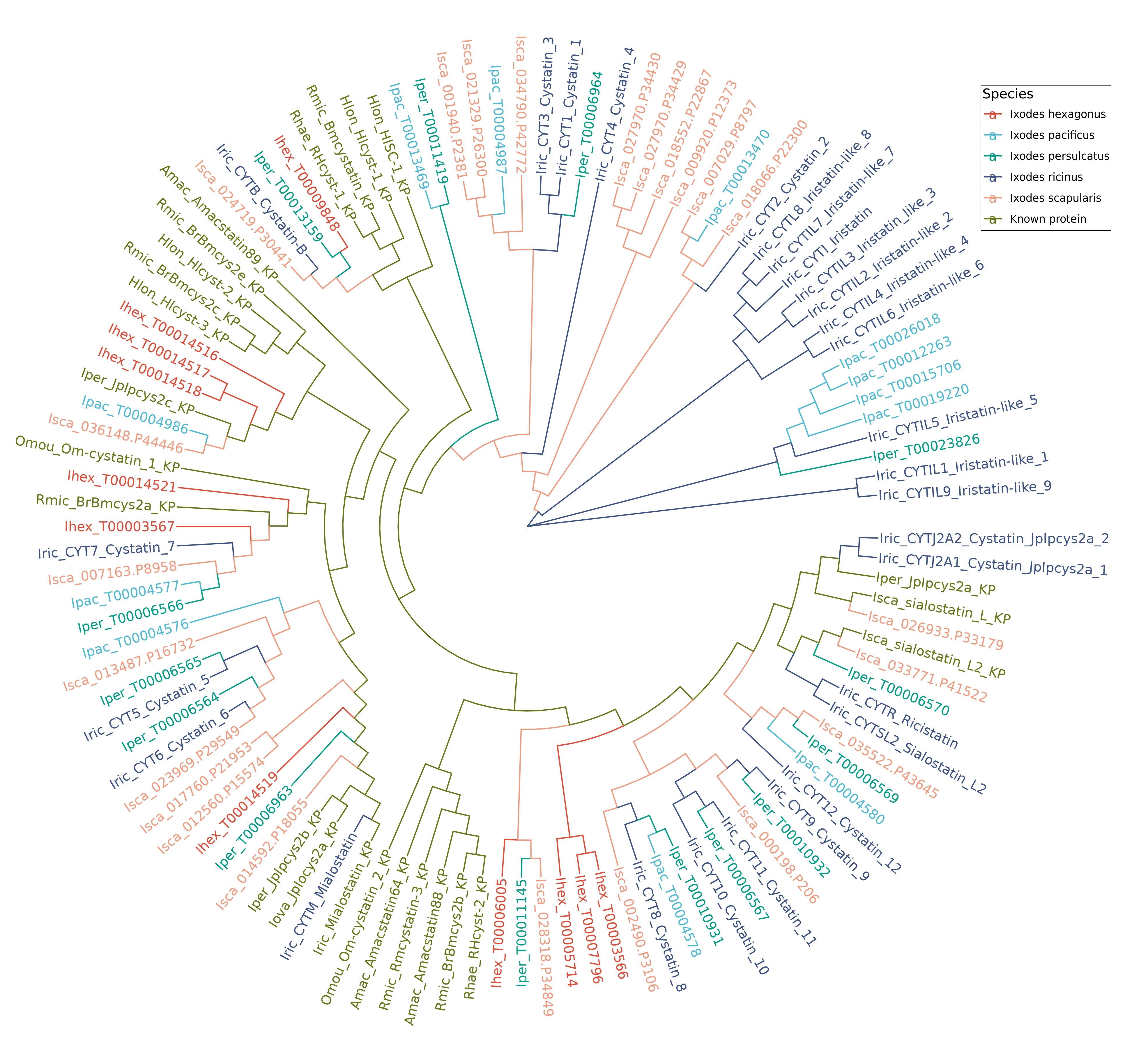

### Figure S06

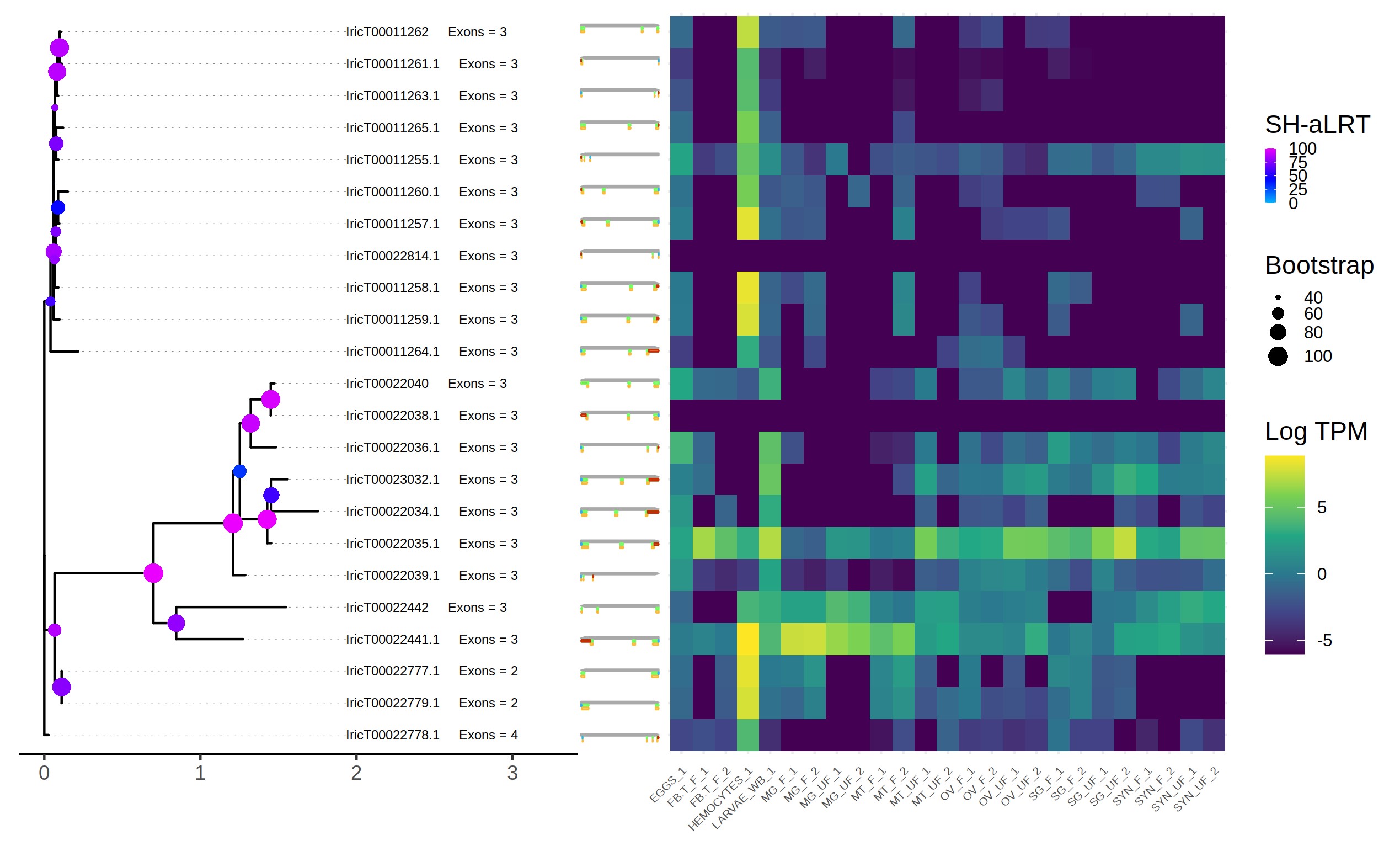

### Figure S07

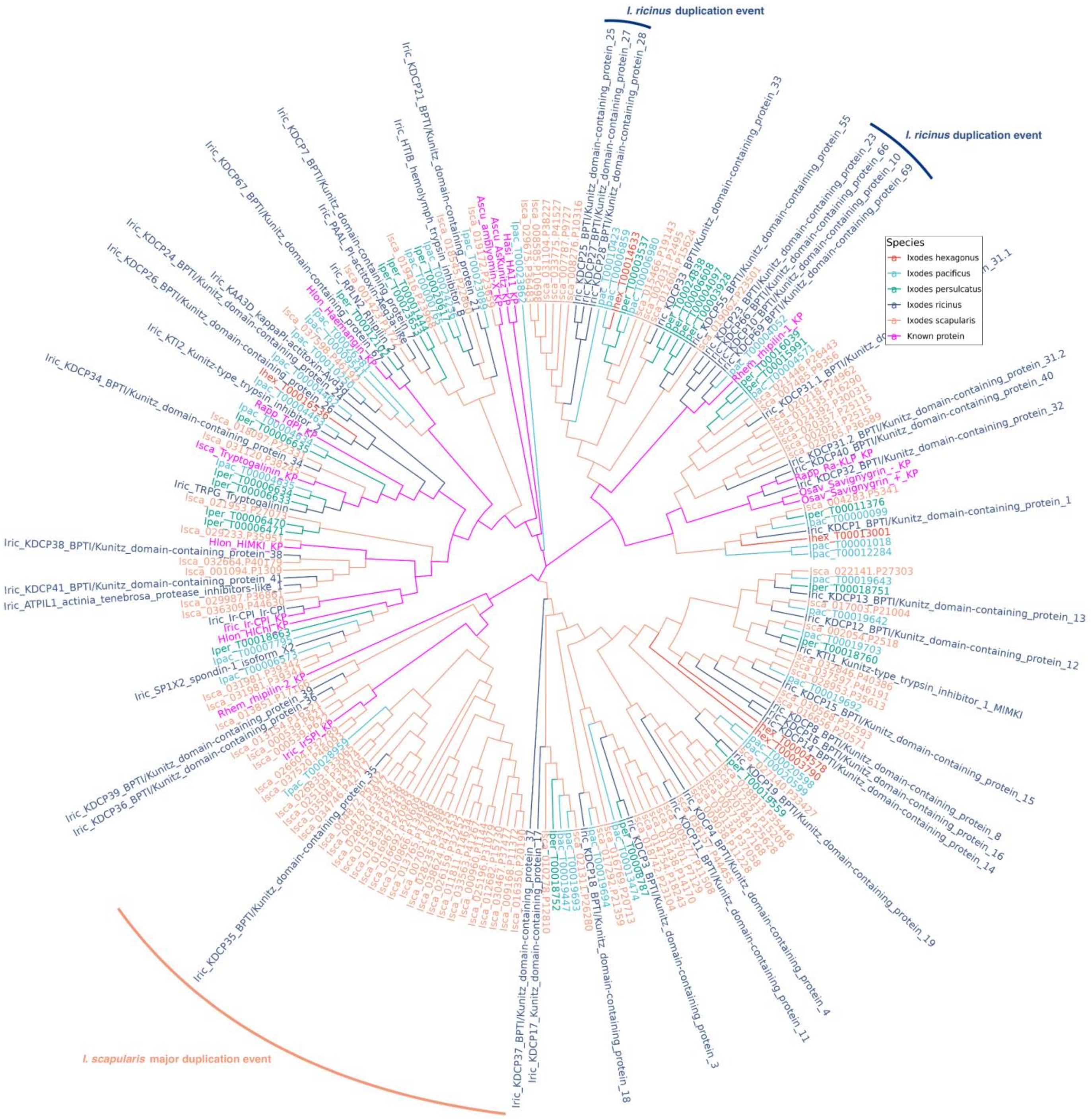

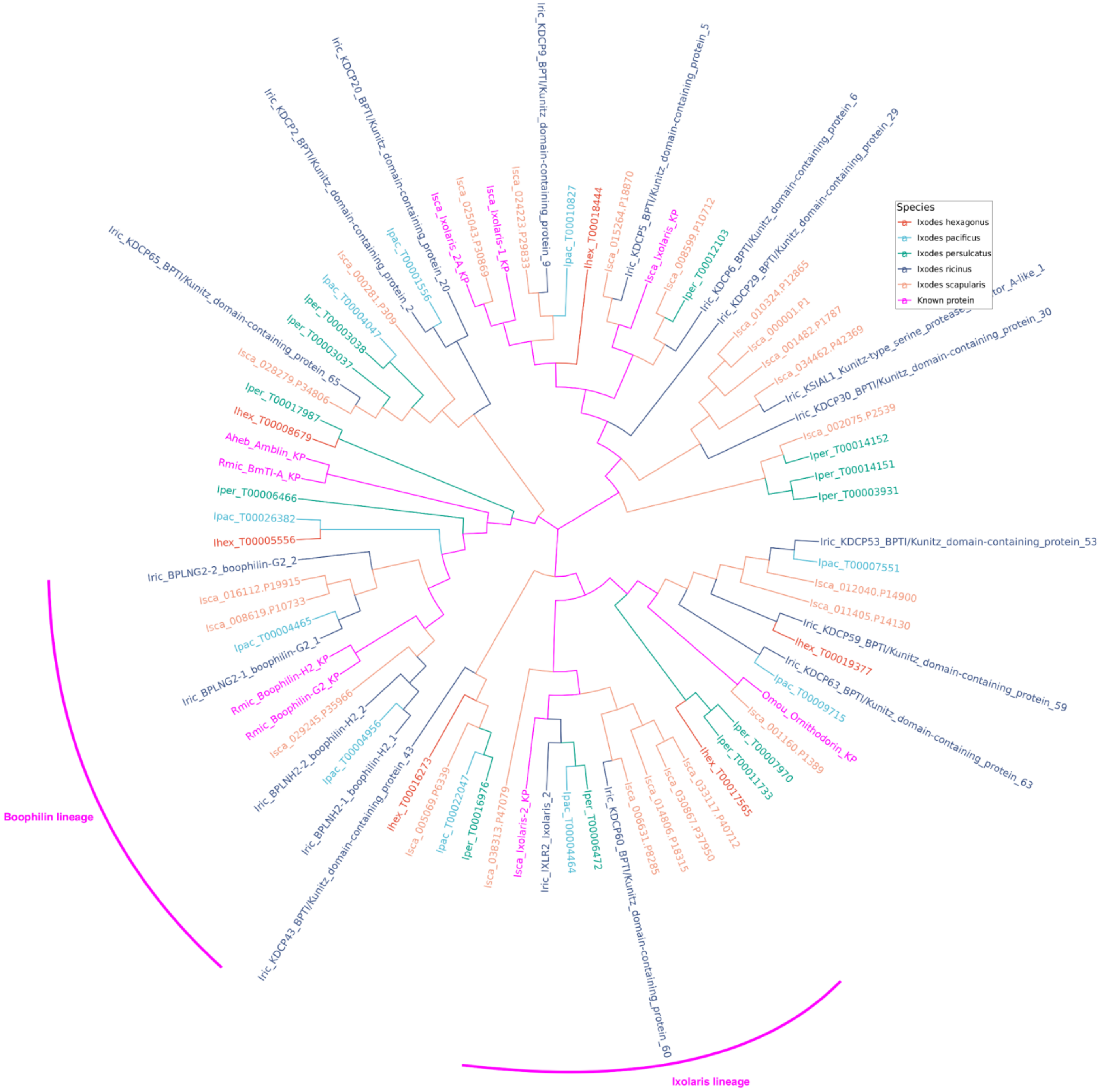

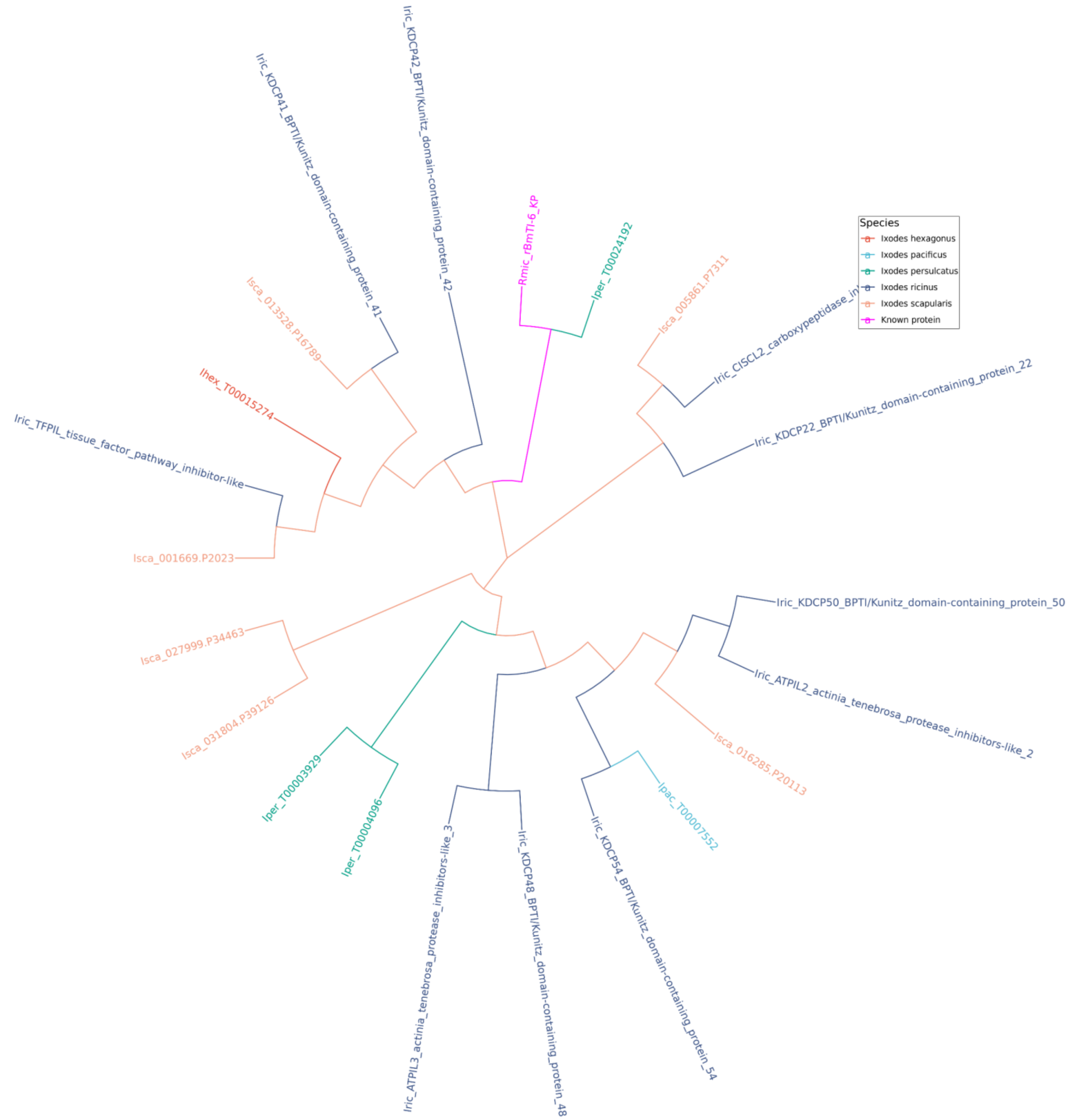

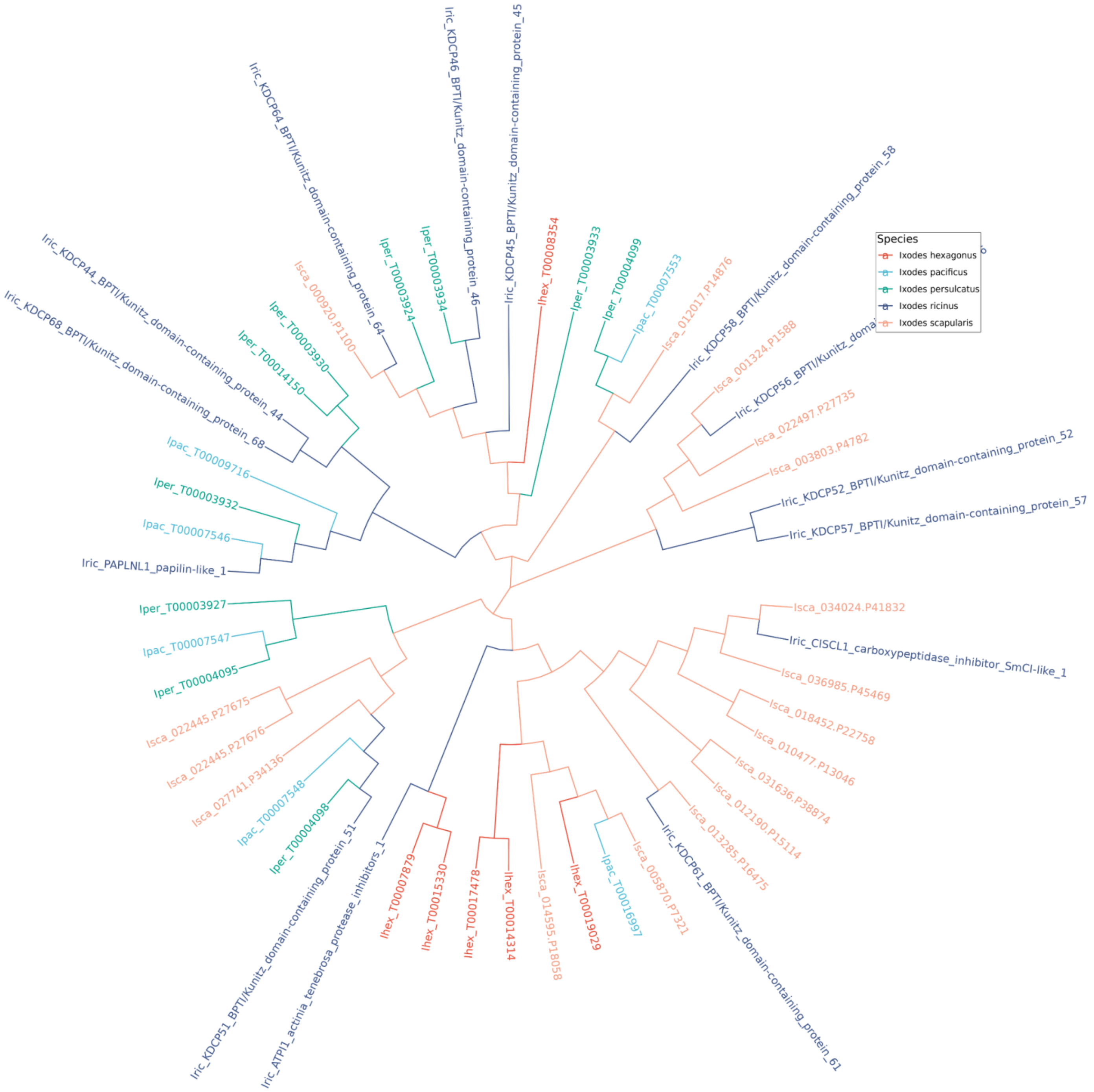

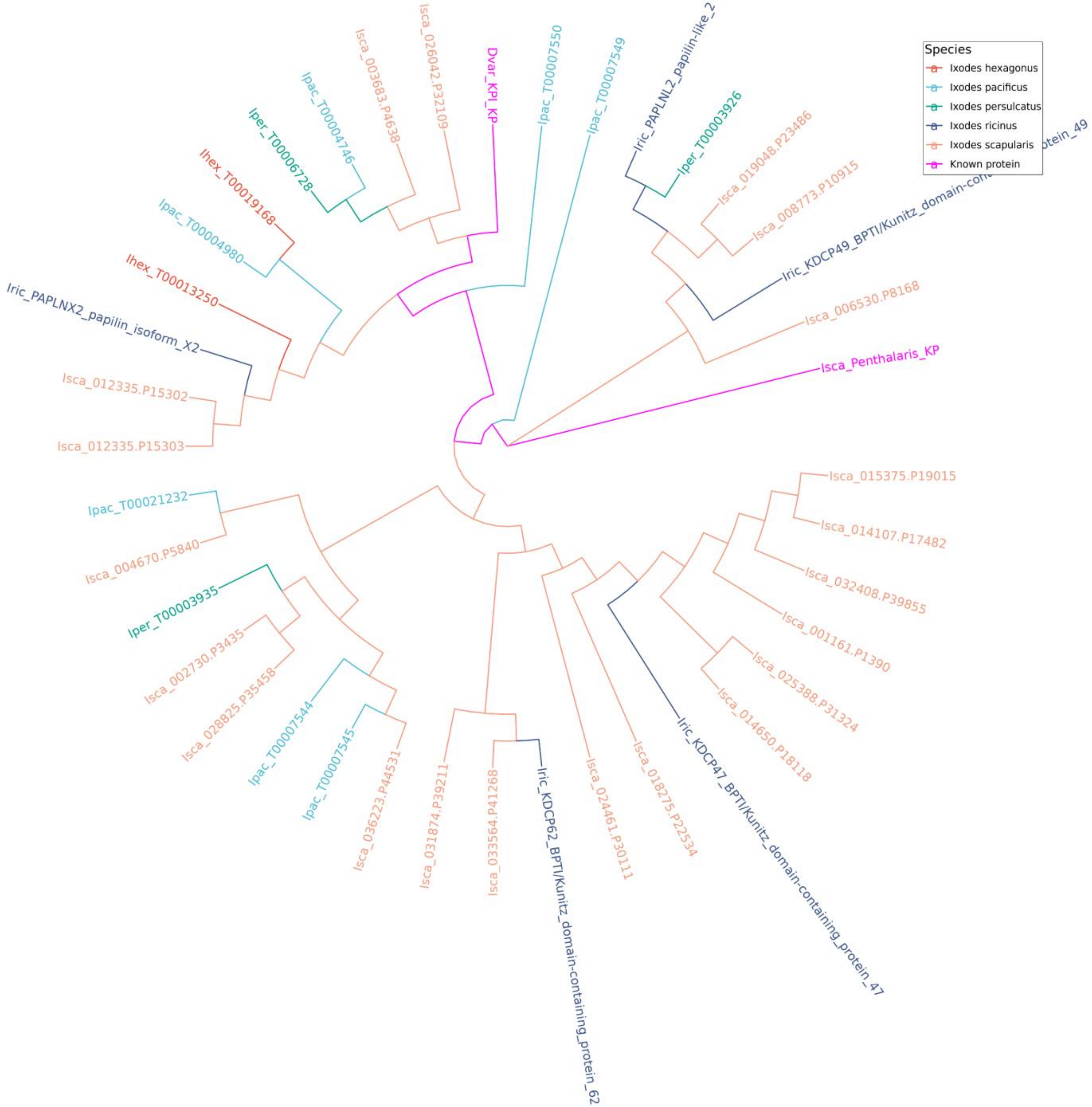

### Figure S08

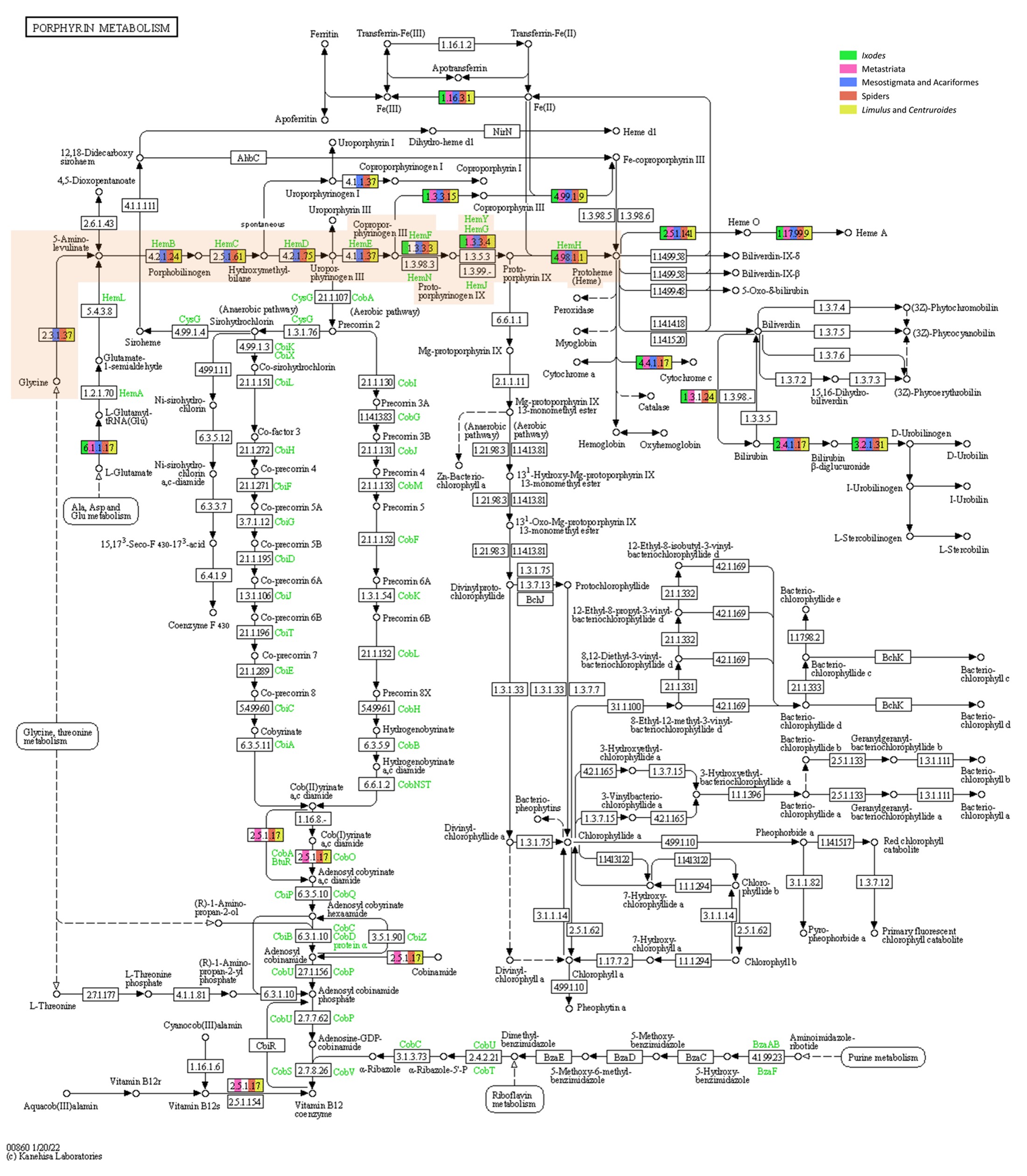

### Figure S09

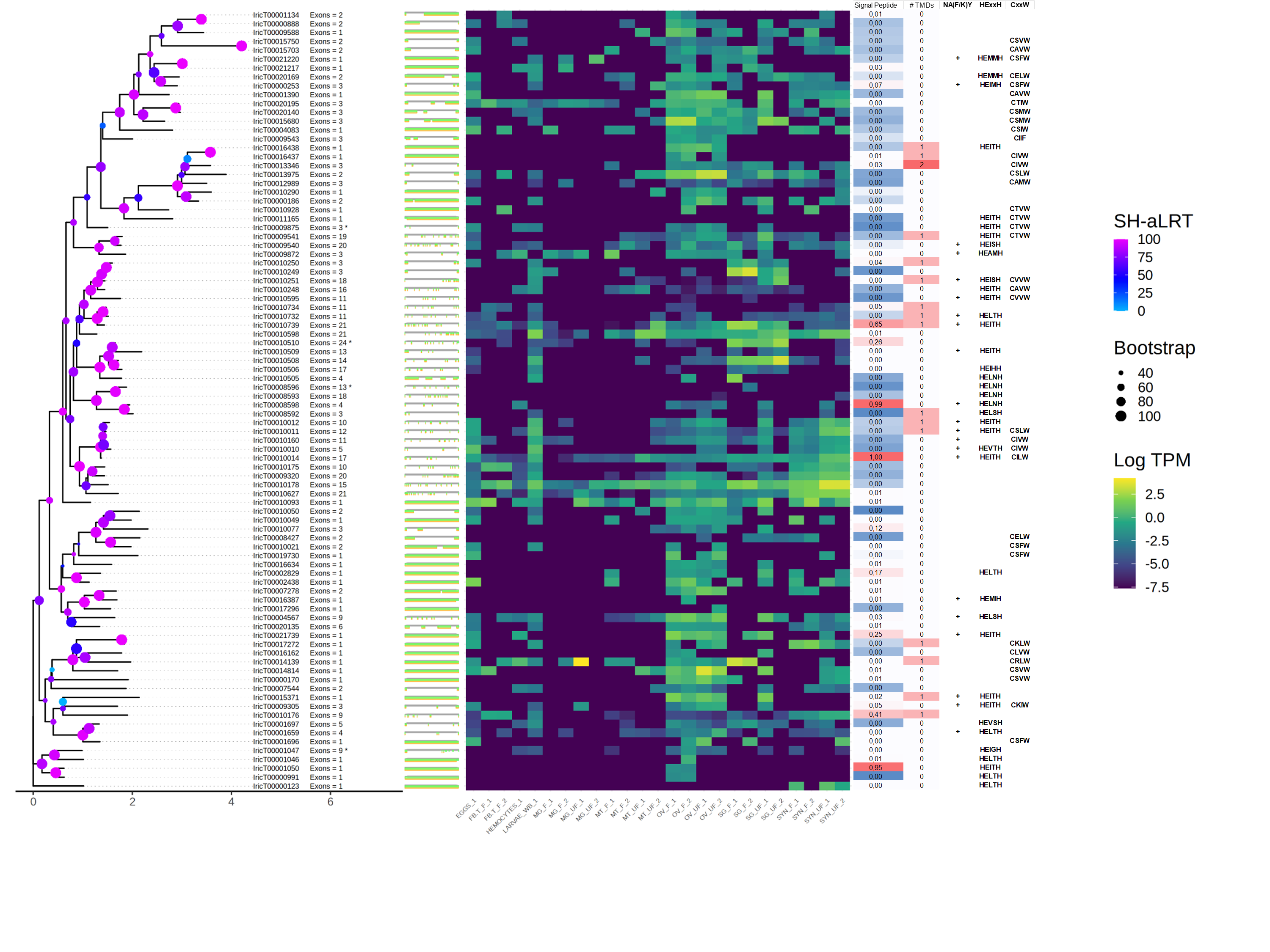

### Figure S10

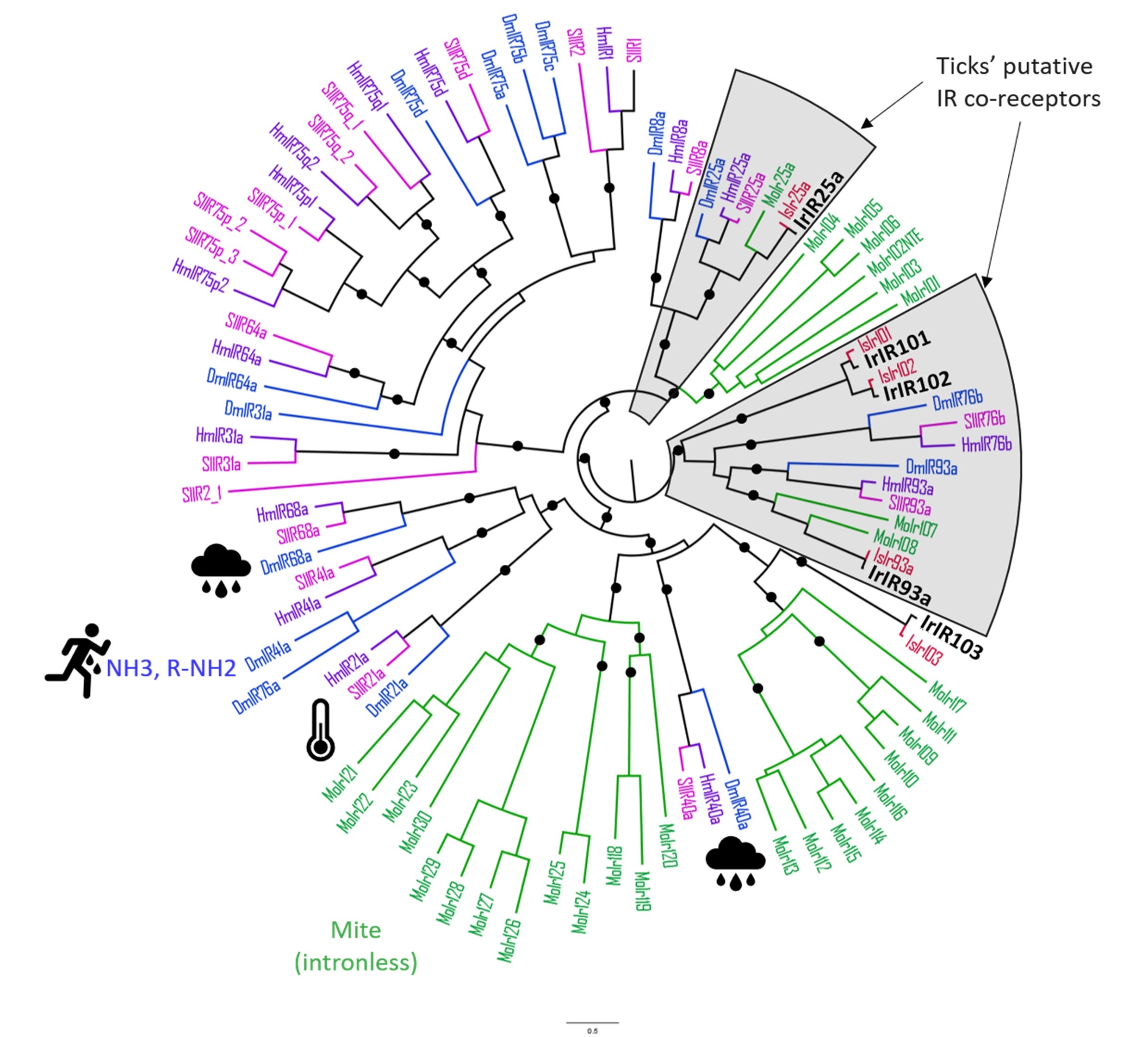

### Figure S11

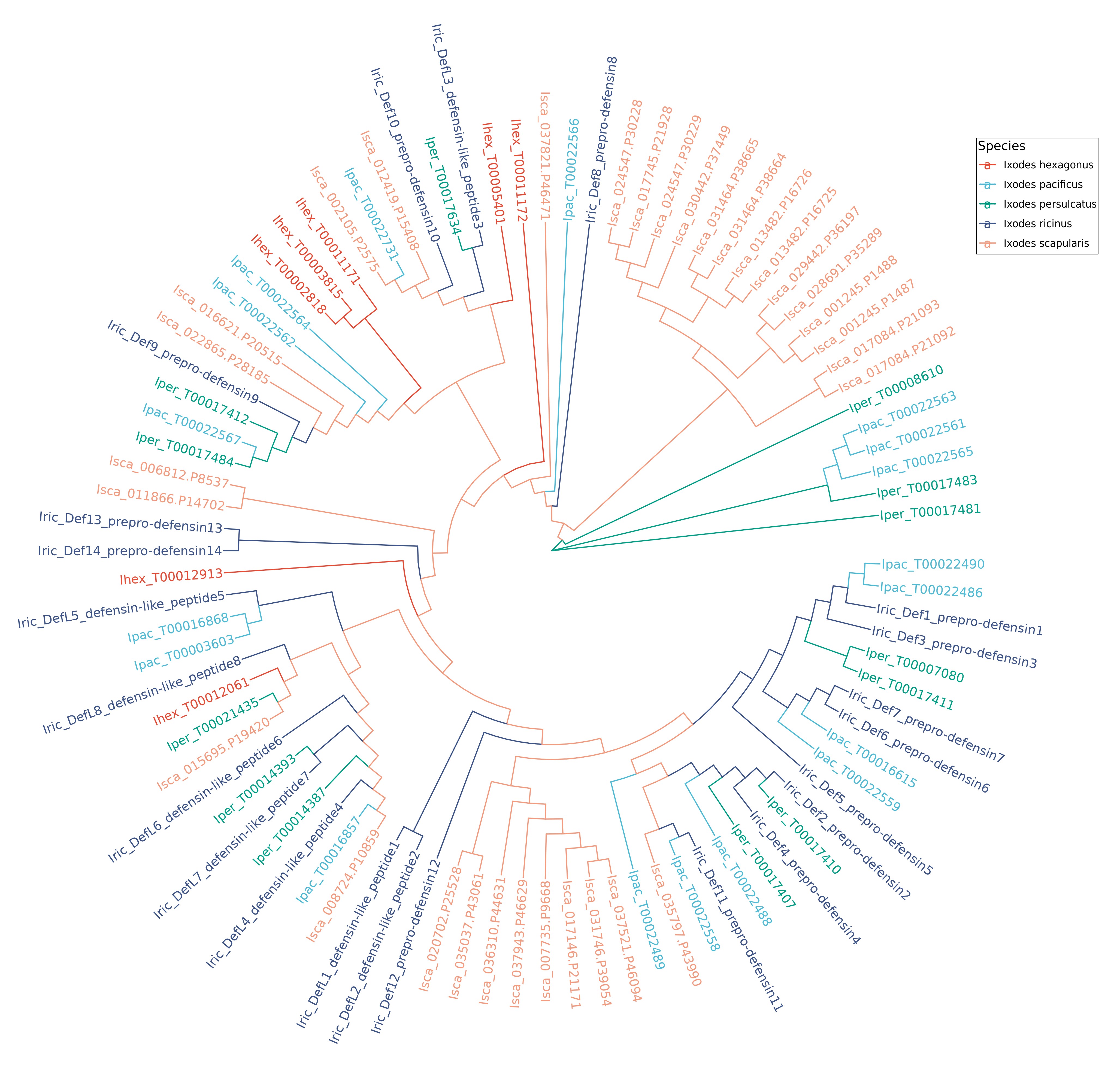

### Figure S12

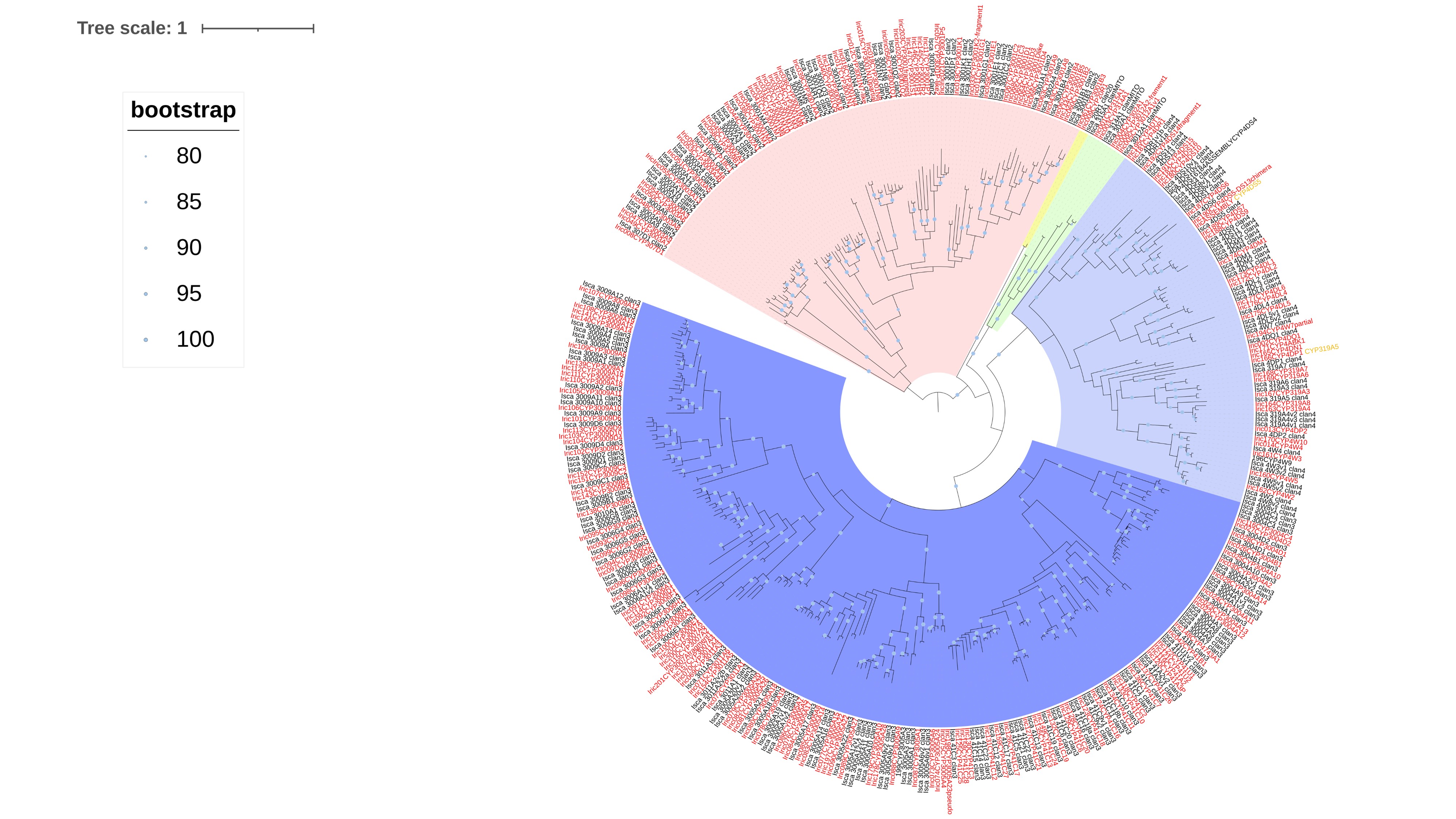

### Figure S13

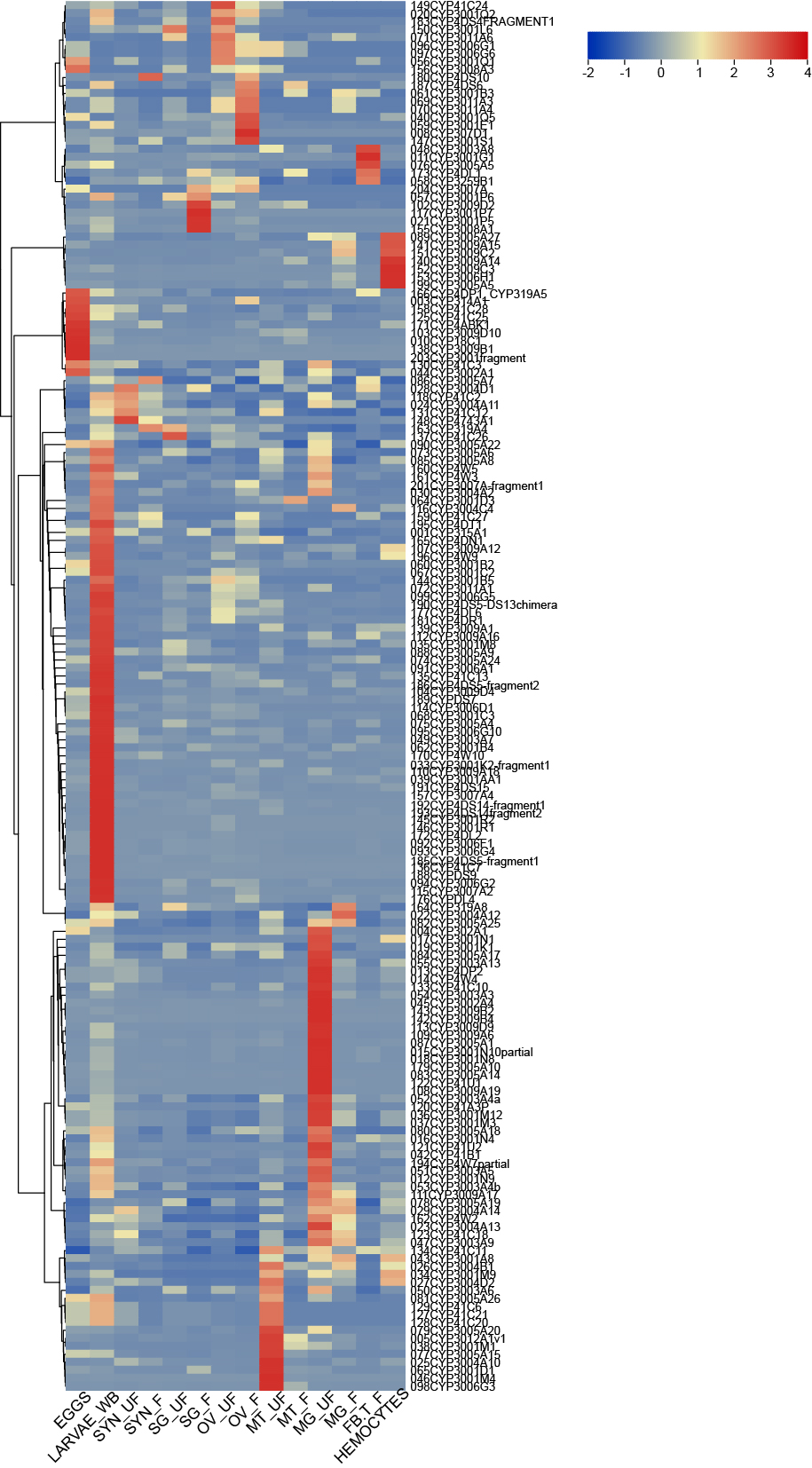

### Figure S14

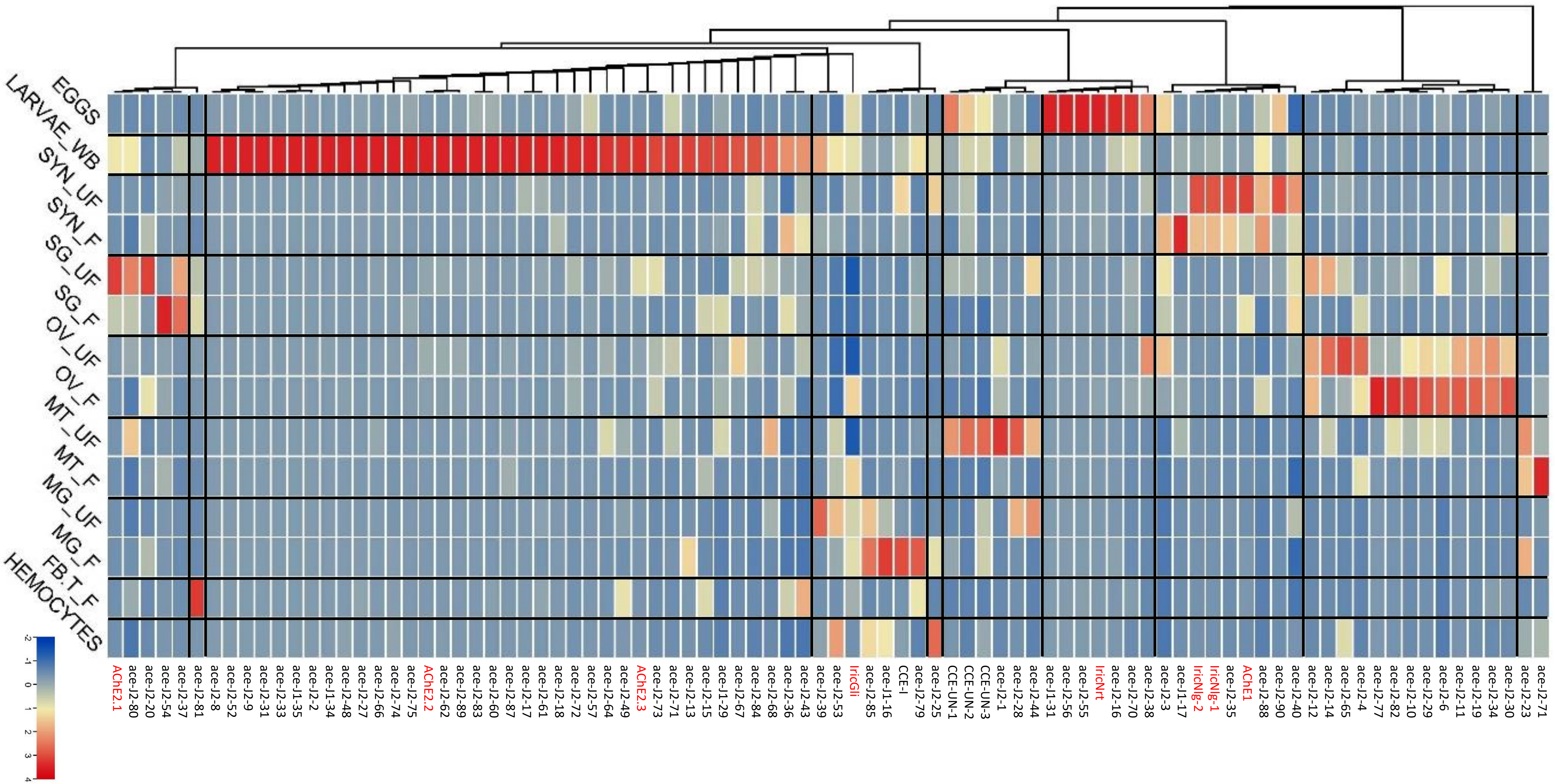

### Figure S15

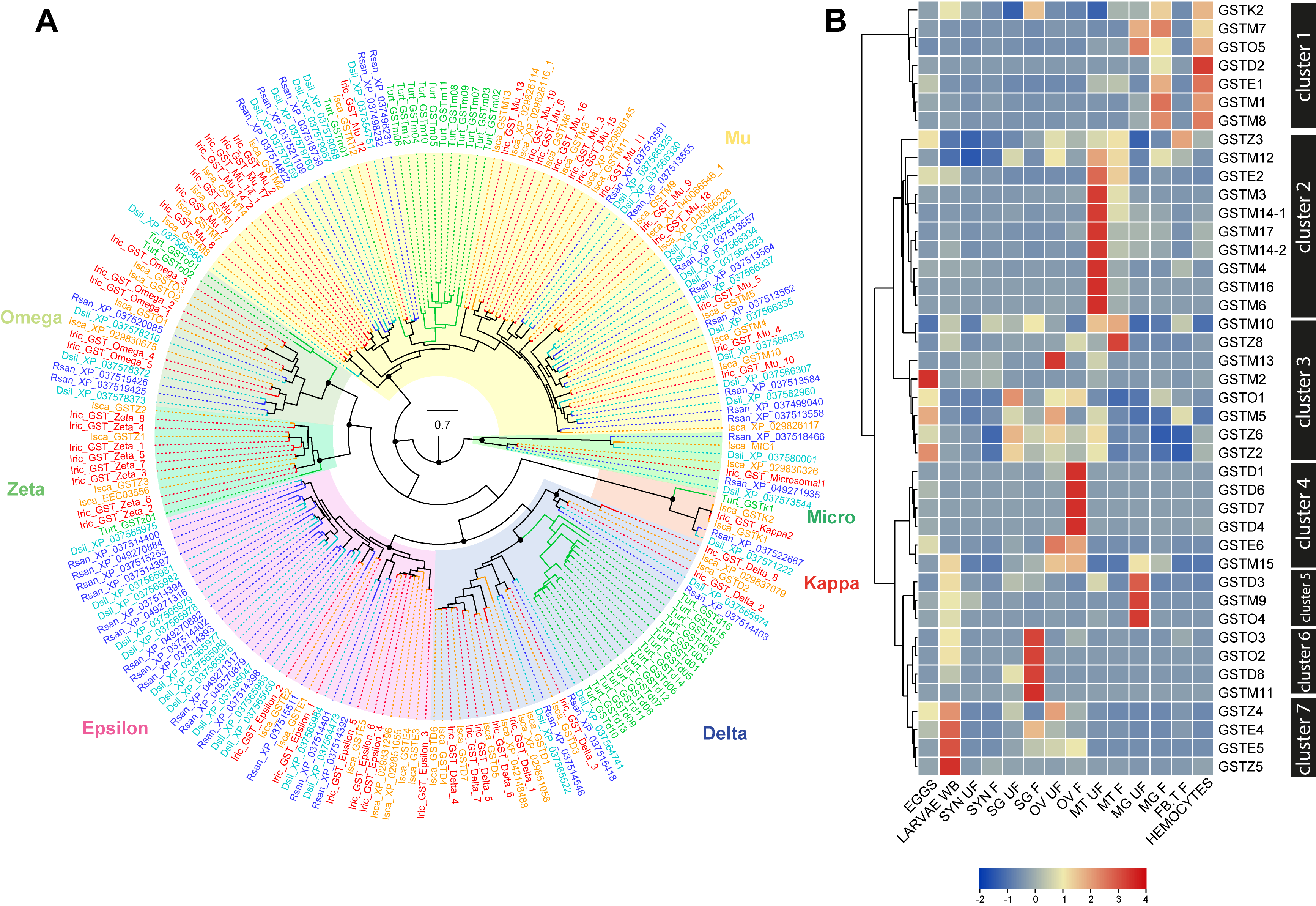

### Figure S16

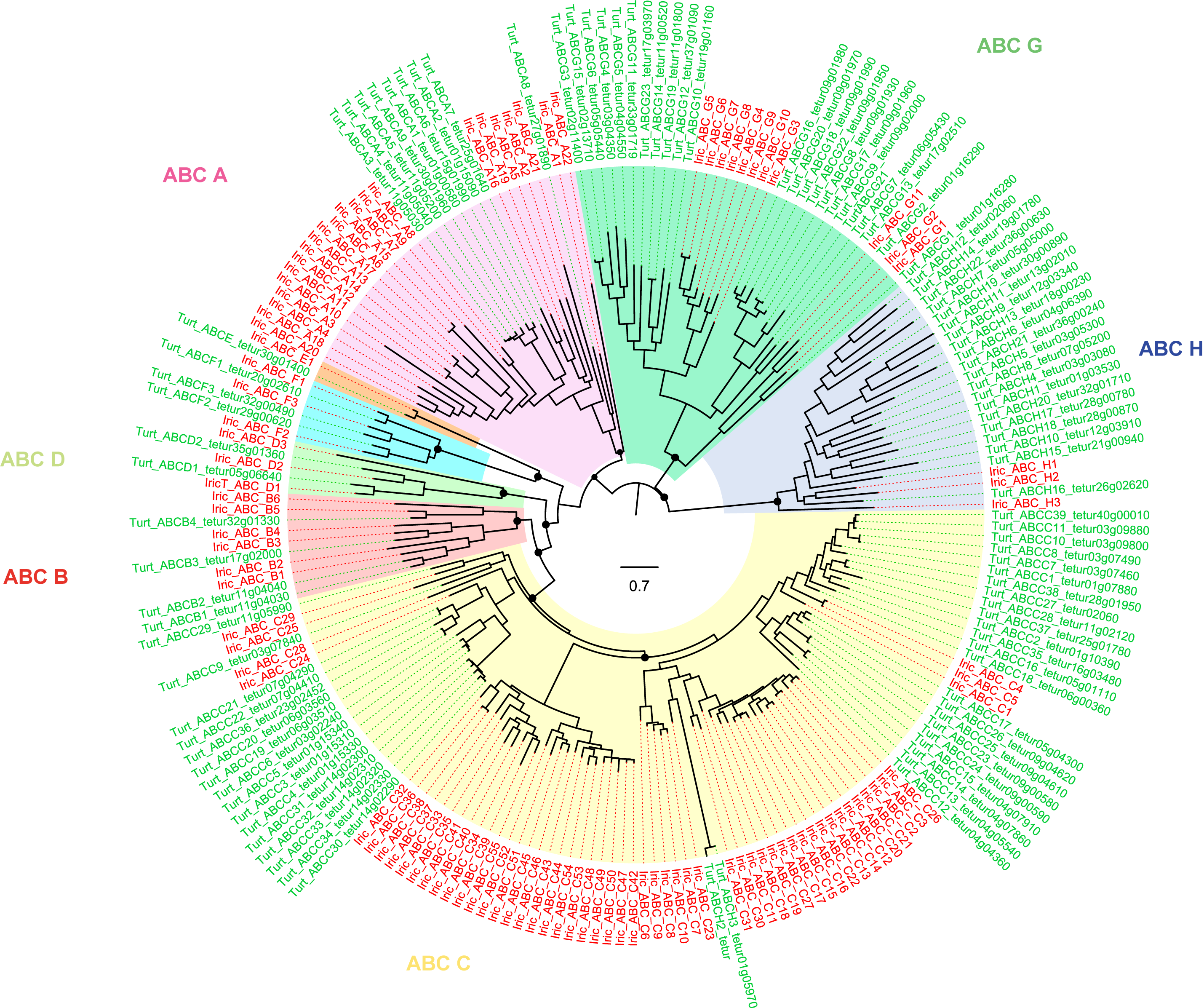

### Figure S17

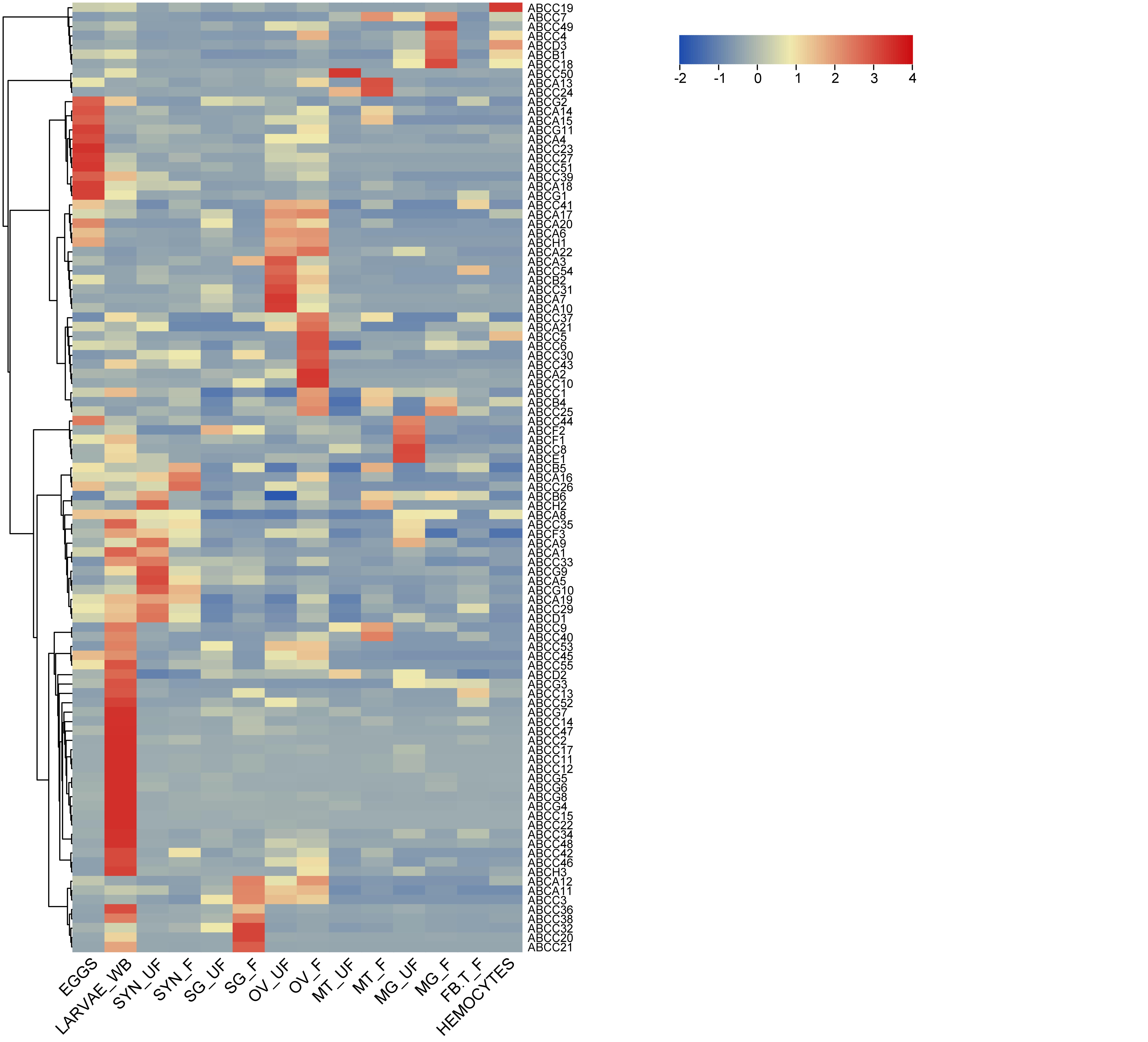

### Figure S18

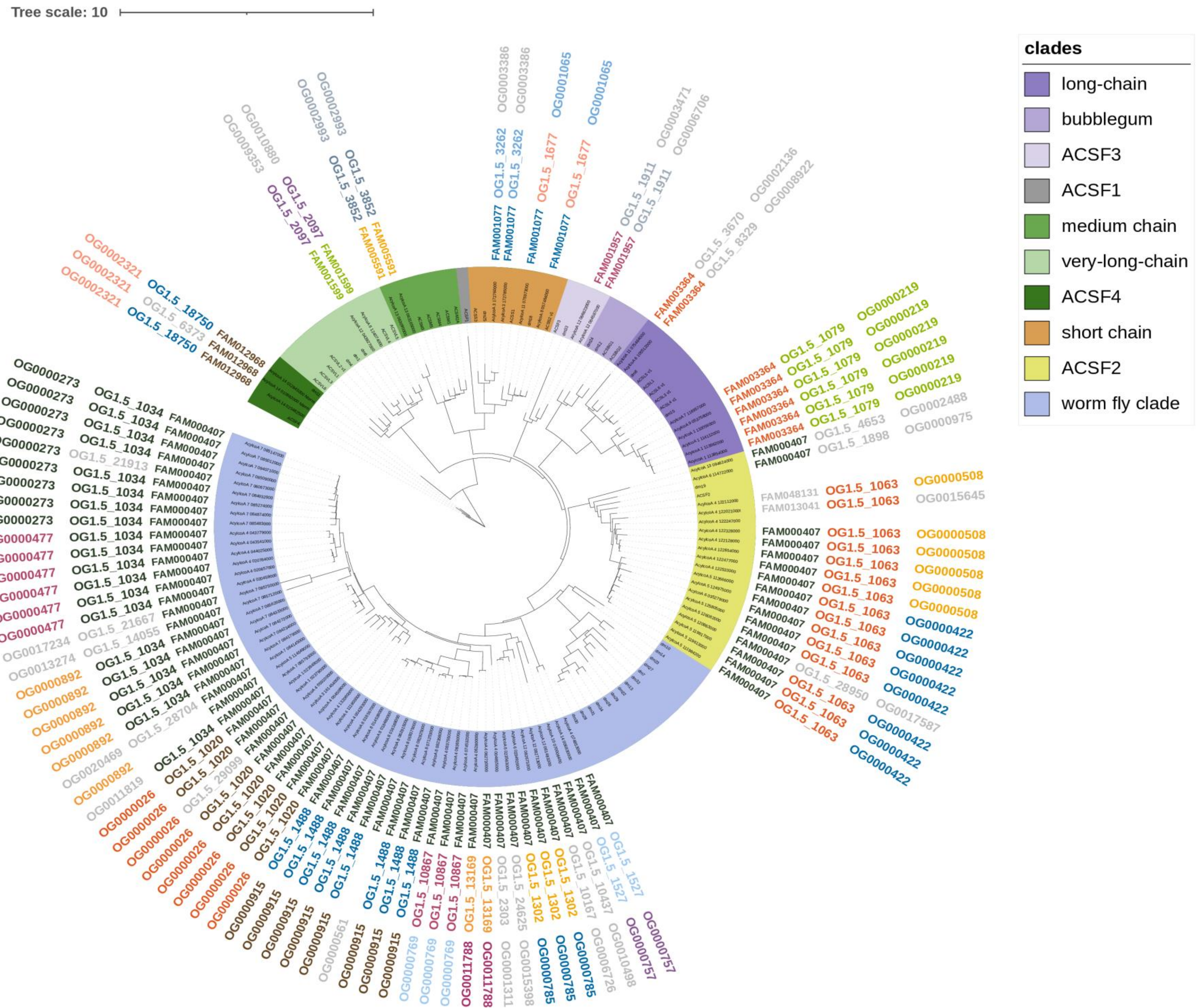

**B**

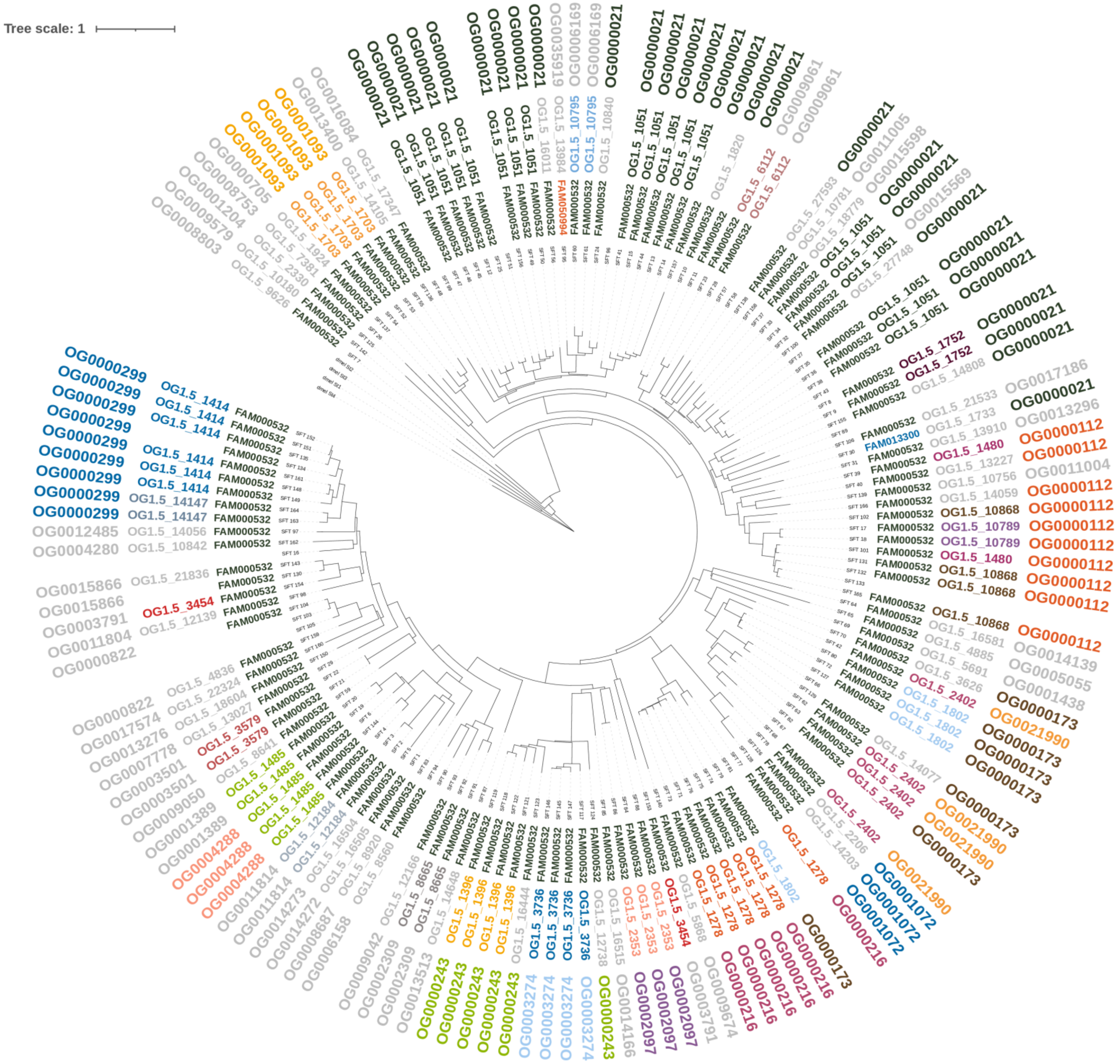
