## Supplementary material for "Genome sequences of four *Ixodes* species expands understanding of tick evolution": Figure S04

**>Achatin**
*MGRAEGFLVLFVLVVVERVALN*SGEGSPAVENQEVFVLNPEAGVRHFRQGKEDLNVDYGDDEPLLVEDLVNKRGFGEKRGFGE
KRGFGEKRRFNERRRLLENSDVIGRGPYLMPSSLVSPWARVYGRR

**>Adipokinetic hormone/corazonin (ACP) related peptide isoform A**
*MALVSKAVLVFLLLGLLVVQHAQC*QITFSKNWQPGKRGEMCSQREAQAVIKLRQVL

**>Adipokinetic hormone/corazonin (ACP) related peptide isoform B**
*MALVSKAVLVFLLLGLLVVQHAQC*QITFSKNWQPGKRGEMCSQREAQAVIKLRQVLLYRYFSS

**>Agatoxin-like peptide (ALP)**
*MNSQAIAMFLVVAVLAGAVLAV*PFNNYDDSFLDEYKEKLENYLMSADKRSCIKRNGICDGRPNDCCHQSACRCNLWGTNCRCM
RAGLFQSLGKK

**>Arginine-vasopressin-like peptide (AVPL, Inotocin)**
*MALSHMLLLAVVGGTSA*CFITNCPPGGKRSSEPSPARLCPRCGPAGRGVCYSADVCCTGSMCVLNDALATLSCRAEALHGVAC
HVPGKRCGTDGRCAIRGYCCGPDGCTKDSSCSGGVPTDQFGSPVDILEYGMSER

**>Allatostatin A (FGLa-related allatostatins)**
*MRPCPVTCLLLLFMLAAQYCRA*DDASPAQLQENDKRRPPAAMYGFGLGKRAPFLFLADDAAERAAERAEAEEEDPDLNYLDKR
GERPQHPLRYGFGLGKRLDREGNYPGSIDHNRRERHRFGFGLGKRGKKSEIEDFMKRRYNFGLGKRSAYGGDDGERWKRSLAS
DHN

**>Allatostatin CC (PISCF-related allatostatins)**
*MGYALSQAVGGLAVLVLLVAASEVSG*QEAWSNLQRGGTASEDGPGTSSSASSEDRTHELLAFKMTSRNIHPDFKTASAKRSTM
LLSKLMQPLFKAFKNDADGGFSSQMELQRRGEGKMYWKCYFNAVSCFRRK

**>Allatostatin CCC (PISCF-related allatostatins)**
*MASTMKIFVLTGAVLLVFGLTGA*QVADESGVRSSGVGLGAGGNGISLENMLWNYFMAKQMSRQAQLQRDGEITTELQRKRSGW
KQCSFNAVSCFGRK

**>Allatotropin (AT)**
*MAALGRTSALVAAALFLCMAAAG*SETPEASDRQHRGGFQKLRLSTARGFGKRIPPGLAFLRQRNQEPADPIIKKGFRKMKIST
ARGFGKREDPDLSFLLENEDMDPVDLK

**>Bursicon A**
*MLICVRSPCSLLASWLILAVLAAS*MGPEESCQLRPVIHVLKQPGCQPKPIPSFACQGSCSSYVQVSGSRYWQVERSCMCCQEM
GEREATKAVFCPKGPGPKFRKLITRAPVECMCRPCTAPDEASILPQEFVGL

**>Bursicon B**
*MKCWTCGVLWLVLLAAAKS*DTLDGGSSCRLQPTTIRITRDQNDELGSLTRTCEGTVLVSRCEGTCISQVQPSITLPHGFLKEC
NCCRETYMNKREIQLQDCFDPNGQKLYGAEGSMTIFLEEPQDCSCHKCGG

**>Calcitonin-like diuretic hormone (CT-like DH) isoform A**
*MQSLLVGFATTLLLAMAVTSA*SSQGHLPKRDVYYDSSLDAGEDYLLNLLRNRVSSHPGLSPALLEQQQKRAGGLLDFGLSRGA
SGAEAAKARLGLKLANDPYGPGKR

**>Calcitonin-like diuretic hormone (CT-like DH) isoform B**
*MMLPVNAGIFVFLGVVLLVEA*SVPAQATYDTESWDRDQQNTAYDSALTPDEDYLLSLLRSRSGSDSLPKRSRGMLDFGMTRGA
SGAKAAKARLGLKLANDPFGPGR

**>Corticotropin releasing-factor-related diuretic hormone (CRF-related DH, DH44)**
*MRCAQSSHPLLRPWLLLALLVTDAALG*ARLGGSKASLDLSDLWKRGPLSLRSRGPVPSLSMHSRDHMPSLSIVSPLDVLRDKM
MQDIIERSIKNKIQANDKILKDLGKRSAPPAAIGEDV
AVSLLRRDDDRSSFSPQESFSDELL

**>CCHamide isoform A**
*MCFKPATPFLLFMLALLVACTQCTAFA*DSCKMYGHSCLGGHGKRSEEGAVSSDVMQMLRKSAAEDDAGAMLAQARLDRPALLE
KTLLQILRNRLQ

**>CCHamide isoform B**
*MCFKPATPFLLFMLALLVACTQCTA*FGDSMEEGLRDKKIFTILRRNTDSCKMYGHSCLGGHGKRSEEGAVSSDVMQMLRKSAA
EDDAGAMLAQARLDRPALLEKTLLQILRNRLQ

**>CCRFamide**
*MLLTYLCLVSSLTVLLLTPSWA*DAQPPRCRALCQNSYAEDACQRCRLRIPMRFGKRDSSRAVALIRDPILDVRSMVLDDELGT
SMSSSREVLETRPSTRTSDQYGLLLRAIRDAAGRSYD

**>Crustacean hyperglycaemic hormone-related iontransport peptide (CHH-related ITP) isoform A**
*MSPSCPVHRRWLAALLLVSGTLVSHQGAQA*RNLHKRSFLELGCRGNFEQSYLARLERVCEECFGLYREPQVYNICRANCFKNE
NFDLCADALLLKDEMSDLRRMINYVYG

**>Crustacean hyperglycaemic hormone-related iontransport peptide (CHH-related ITP) isoform B**
*MSPSCPVHRRWLAALLLVSGTLVSHQGAQA*RNLHKRSFLELGCRGNFEQSYLARLERVCEECFGLYREPQVYNICRENCFKNE
NFLKCAEALLLHDEMESLKGKVDYLYSR

**>Crustacean cardioactive peptide (CCAP)**
..DDTADDSYLVEQKRPFCNAFTGCGGKRSSPNRIDLLARLQNRLLSEIRNMELRTRLEEGPSRRHDGYTDNLRSNGLLMDLL
TPSFRKRKFPMEGP

**>Corazonin (CRZ)**
*MSHYLGLSAVLLVCLAVTAYS*QTFQYSRGWTNGKRRVAEMPLVGVPVRASNDHRALDEVLSKFTPRDRIVLERLGHMVRVLDH
AEEEQQEY

**>EFLamide**
*MSLPQLLLLVVLVTCGFLCVANA*SMEAKRMGSEFLGKRSRMDAADLISGLWEEDGQPRPSAARVRFLRTAAALNRLGALEDDD
DALEASPLMSDLSAAALYAKQKRIGSEFLGKRSSADLVQKRMGSEFLGRRKRDAVHGDFPPPVVGPLDA

**>Elevenin**
*MKRTCIQAVVGVLFFAALVHQLHA*ELDCRKYPFYYRCRGISAKRSFAPITKMEAMSLKELYEDDDGWKNRRRPADAVLGWVRN
KYGDDIFDPDEPLDTTRGSFERKELY

**>Enterin**
..SIRPFAKRTTPYSGYRKIFYVGKRATAGQHLP..

**>Eclosion hormone (EH)**
*MARIFDFTIFLVVSTATFAVLLSLSSA*THTYPSDPVLVCINNCGQCKMIYGEYFNGRQCAEECLSTAGFIQPDCDEADSIVKY
LRRKP

**>FMRFamide_Myosuppressin-like**
..MAIMGPRRHRYLHFGRKRALPLYSDVPVDQVEGSDDYIGDDYDASSAESLEDAMRWAKPYLGDGLHGDVVLSGALEDGQVI
RYKRDVSMASVRDDELDTNTDQKRRALIEVHDELARDGNAGPYLDWQGREKKSQNRILHFGKREGQHESTQLGSDEVQGDIKR
AMNRILHFGKRVRDDATSDFQGDDYGWYSSGDKRATNRIMHFGKRQPEEILISDESSGPQIQIHGNDKKSINRILHFGKREGN
EAFDTDLVESGYRKRNARSNRIMHFGKRTDGGLTSDFDPESHTALKRATNRIMHFGKRESALSSSLEDQLKRDFFEWKKRYTN
RMLYFGKKRPQDRYTDKRITNRIMHFGKRGVIFPLSDETDDSSGKQKRQLKNSILHFGKRDDEKSIEKRTRNRIMHFGKREEG
YPYENRLTSDKHLGDRILHFGKQEPHHQAADFLNKRSTANADLQFDNEDNESPYLVDKKITNRILHFGKRLDDSAEDPGKVSG
KPKQHVPSVNSDIKFKDSFLFEEHKPHNRRKRSLGFDQYDLDETLERVVHQLMDAGYPKRVALGHPGIPGHLHLPHAFVAAHV
YGSELPRMLSRPSRSDRFFPVPYSGEHREAPKGPSRNVFLHFG

**>Glycoprotein hormone alpha**

*MSRVNQFVAVFVAVLVLGQAGA*NFWEHPGCHKVGHTRRVSIPDCVEFDMTTNACRGFCTSYSIPSPEYTLRMNPNQGVTSFGQ
CCNIMDTEDVKVQVRCLDGHKDLTFKSAKSCSCFHCKKN

**>Glycoprotein hormone beta**

*MNTSRMLRGALAVAVVVLCPLFGDA*SVGVVDTQTTLDCHRREYTFKATRVDAKGNRCWDDVTAMSCWGRCDSGEIADWRFPFK
KSFHPVCTYDSRRLVTTQLRNCEPEDLDEEDELRTYDYFEALSCSCQVCDSTWTSCEGFRHP

**>IDLSRF-like peptide**

*MLTRATQHLFFLGVVLVGLAQG*AYLINFSHVLNRNLGNVKREVDTERCHPTKPFRCPSSATVCISIQYLCDGAPDCPDGYDED
PRLCTAAKRPPVEETANFLQTLLSNHGPNYLEKLFGAKARDALAPLGGVQKVAVALSEGETLEDFGKALHLMRSDLEHLRSVF
LAVESGDISLLRSLGIRDSELADVKFFLDKLVATGFMD

**>Insulin-related peptide 1**

*MVSWALNTVVVALVAASALVAPAAA*GSGRRCGKILLEFMEFVCEGEFYDPYENTGPKRSLIGQRLYPLVSPGIENTDKAPASG
FLRAESASQLLRKRNFQGGIVFECCYKACSIMEAQSYCPS

**>Insulin-realted peptide 2**

*MNAAVLLLLCATALLSSHRGASA*RSGVEKRNNRYCGNNLNRVLEFLCEEYYDPTQKRHTGYRPAHDVPATLPVWFPVLDANGD
SKMGFMEAKAALQLLRPSVRYGRQTRGIVEECCHKSCSTLELLAYCKTPRNNADLQVSSDDNTA

**>Kinin**

..MSMGGLCQDVDSGSGDLGRSSRVGESFIRWNISPATLQHMRSESFKRQFSPWGGKRGVLDQALPTAHRLSGPLYLYKALHS
PGAMGERRGDKQPDDETFNPWGGKRENDKDKELSFNPWGGKRGTFSSWGGKRDTFGPWGGKRDTFGAWGGKRDTFGPWGGKRD
TFGPWGGKRDTFGPWGGKRDTFGPWGGKRDTFGPWGGKRQDKESGFNPWGGKREDPFNPWGGKKEDKNAFSPWGGKREQNFNP
WGGKKTSKDSTFSPWGGKREGPFNPWGGKKGDSDTAFAPWGGKRDNNFNPWGGKRDNGNKDSSFSPWGGKRESFGVQASDPDS
LEDHSPSRNKRDSSSRVPKTKNSARSAISSVAKTF

**>Myoinhibitory peptide (MIP)**

*MSPVESSRHVGRRPVVATYGESGRTATSAVVLSRSLLVLLVLAALLCCGA*AEPQPQGGDWNALSGMWGKRASDWNRLSGMWGK
RAGAYGPYQALLLRAEESNDGAGHGISARAAPPGSPRENHWNDLSGYWG

**>Natalisin (NT)**

*MSDQLLWISFLLMMGCLA*DPLRAQLLQLYSGDVDPRTTGEGTQLVGLMQRLLRDAALLRKRSPDGDTLPPGFVGARGRRQEAD
GPPGFVGARGKRPSLVIRGLADGISSAAGDAY

**>Orcokinin (OK)**

*MTSLLFVFLLAGVSLCSA*LIDGHEAEPGKGVRTLDKLSGGEYIRALHRLGGRRLDKISGGELLRAMPESQDRSSGEVLRSMGG
PYALRRLTVPRGLDRISGGEYIRAMGSSGFPAGPASAKRFDSLSGLTFGGDQHAARKRWYGHGDFDEIDNVGWPGFTKRNFDE
IDRTGFEGFYKRAANAAAAAPAARED

**>Pyrokinin diapause hormone pheromone biosynthesis activating neuropeptide (PBAN)**

*MGANRQLLMRAFWLQLLFSTLTLGVGG*HEAYYEEDGGLWGPSEVKRQGLIPFPRVGRSAGRDLGRGPEELLTLDTELGSDWAF
LLLPYKRRSNNFTPRIGRKRRSVSEDGGHGDSSDMRALSRHSWPGLEWSYPRMSQQMIPVPRNGRGSFVPRLGKRRMGYDDPE
SWDSREYSASGDPKRGSFTPRIGRAAFTPRIGRTPFTPRIGRSGDSNKDTMSMDDKTQSASGSGSNSRSTV

**>Prothoracicotropic hormone (PTTH)**

*MLVTPRVAARLTRSLLVILLVAAGVET*SPKPACSRATREDLERLLGSAFNARYMAIDKPDPEPTSSRLTARHSEDPLSDQADL
EAGGFSVDADFRQDLPGERRRVREKRSETAKPWGCASRLEWEDLGDDRFPRYLRTVKCLGGDCWFGKFRCKARAFTVKVLRRK
TAKEKDGDCVMSASTADLPAELREHWEFEERAVAFCCDCSVDD

**>Proctolin**

*MVSQTRLLALALMSTLMLLVVDA*RYLPTRSDDLQKDHIRDILRGLFEKAEFEKSASNLLADLGSAYTLRGGAPLGSDMGAYAG
SRAGAVRSGLLSRDMGA

**>Relaxin**

*MLRCATVWLVVSAVMDLATG*TADTPNWEEIFRNRNDEDWARVWHVERHRR**C**YHQLVSHMNLV**C**REDIYKLNRRRRDVAADKDP
EMTDLFLKPEAALGVLTGKLSKRRATQHNIRTRSIIDE**CC**DTEVG**C**SWEEYAEY**C**PANRRMRNRRR

**>RYamide**

*MLHCRAWAVALIALLVLSVASA*AKGTPQFVPNGRYGRRSVTPPLAGVSRDITVNFFGDSSISCTHTGFADIYRCTR

**>SIFamide**

*MNSWKAFFMFGTLLVMAVMMNMACA*AYRKPPFNGSIFGKRSRADLNNADVKYAMCEAVWDTCTQWFPITQDGAQ

**>Short neuropeptide F (sNPF)**

*MVSLRRRPASFSIACVFVVLLVAGTFA*AAYADFNGERDMRDLVELLLKSEQESQLSHTMERKGGRSPSLRLRFGRRSDPAWSD
SLHRFLAAGNAAGDSGHSAPAA

**>Sulfakinin (SK)**

*MRASSWFLICLLAALVYGSWS*SPTSMQQRHRMAMGKWLKSVLPGAPSGGDAGSRNSGDIDTDMIDPVILANGFAKRQDDDYGH
MRFGRSDDYGHMRFGRK

**>Tachykinin-related peptide (TRP)**

*MDHPEMKVVALWSLVLLLTLSSVRT*ASFQSSEVGNELQSPEVGEVSLVEKLGWDGGLDGGDDLEIAAAADDEKRAFHAMRGKK
DDPSLDWDEADKRAFHAMRGKRLLAPASVDSFIAQLRRAVLQGKRGSGFFGMRGKRMSRTPNKEHPRSTFVATRGRRSVLSEA
ESRPYY

**>Trissin**

*MASQVAQILRYAPRMALLLLLCGLEGQA*SRA**C**NA**C**GPE**C**VTA**C**GTAMFRA**CC**FNYNRKRSVDQRDALTPAPLDEAAFETPVDP
PQRSLDAWTLLWVPRRAAAAMPTNLF
